# Functional Decomposition of Metabolism allows a system-level quantification of fluxes and protein allocation towards specific metabolic functions

**DOI:** 10.1101/2022.10.22.513080

**Authors:** Matteo Mori, Chuankai Cheng, Brian Taylor, Hiroyuki Okano, Terence Hwa

**Author notes:** These authors contributed equally to this work (Matteo Mori, Chuankai Cheng).

## Abstract

Quantifying the contribution of individual molecular components to complex cellular processes is a grand challenge in systems biology. Here we establish a general theoretical framework (Functional Decomposition of Metabolism, FDM) to quantify the contribution of every metabolic reaction to metabolic functions, e.g. the biosynthesis of metabolic building blocks such as amino acids. This allows us to obtain a plethora of results for *E. coli* growing in different conditions. A detailed quantification of energetic costs for biosynthesis and biomass growth on glucose shows that ATP generated during *de novo* biosynthesis of building blocks almost balances the ATP costs of peptide chain polymerization, the single largest energy expenditure for growing cells. This leaves the bulk of energy generated by fermentation and respiration (consuming 1/3 of the glucose intake) during aerobic growth unaccounted for. FDM also enabled the quantification of protein allocated towards each metabolic function, unveiling linear enzyme-flux relations for biosynthesis. These results led us to derive a function-based coarse-grained model to capture global protein allocation and overflow metabolism, without relying on curated pathway annotation or clustering of gene expression data.

## Introduction

Living cells perform thousands of distinct metabolic reactions in order to grow, maintain homeostasis, and respond to environmental stimuli. Understanding the coordination of these reactions, their associated costs, and their contribution to cellular fitness are grand challenges of systems biology. Protein allocation has been established to be a key factor determining bacterial growth, owing to constraints in protein synthesis for rapidly growing bacteria [1–4]; accordingly, the abundance of proteins catalyzing these metabolic reactions have been shown to follow simple rules of allocation [3–7]. In order to integrate genome-scale data of metabolic fluxes and protein abundances, metabolic models are increasingly used as multi-omics platforms [8–13]. The inference of intracellular fluxes based on mass-balance constraints and optimization principles is a mature subject [14], with Flux Balance Analysis (FBA) being the most celebrated framework for study the metabolic capabilities of a wide variety of organisms [15]. Extensions of FBA have been developed to predict how global protein allocation impacts metabolic activities of the cell and vice-versa [11, 16–21]. Despite the need of inferring a large number of unknown molecular parameters [22–24], these models were able to recapitulate known metabolic features, e.g. the emergence of overflow metabolism [21, 25], or the global utilization of the proteome [26], in relation to cell growth.

Given the global constraint in protein synthesis, the amount of protein needed for specific metabolic functions, e.g. the synthesis of a specific amino acid, is important for quantifying the link between metabolism and cell growth. However, the deeply interconnected nature of the metabolic networks also complicates its decomposition into individual components [27]. For example, central carbon pathways such as glycolysis and the TCA cycle are not only tasked with the production of metabolic precursors for biomass building blocks, but also with balancing the “currency” metabolites, e.g. ATP and NAD(P)H [28], consumed by each biosynthetic pathway. Thus, it can be difficult to associate reaction fluxes, and the corresponding enzyme concentrations, to individual metabolic functions. Indeed, the intuitive notion of individual metabolic pathways is largely concocted, as the production and consumption of currency metabolites have to be balanced across conditions, thus effectively coupling the fluxes through all pathways. Thus, the calculation of costs and yields for the production of individual metabolites are often performed by making use of coarse grained metabolic networks in which the balance of currency metabolites or pathway byproducts is simplified or neglected [29, 30].

Here, we sought to develop a systematic computational method to define the metabolic costs and the enzyme amounts associated to each metabolic functionality, thus cutting through the complexity of the network. We introduce a Functional Decomposition of Metabolism (FDM) based on the decomposition of metabolic fluxes into a set of flux components, each associated to a *metabolic function*. Being based on properties of optimal flux pattern such those obtained with FBA, FDM is generally applicable to any metabolic network, and does not require additional parameters. We applied FDM to *E. coli* cells grown in carbon minimal media, under translational-limiting antibiotics, and in anaerobic growth. The resulting functional characterization of the metabolic reactions allowed us to analyze in detail how cells allocate nutrients towards biosynthesis and energy generation, as well as the metabolic costs and yields of the production of different biomolecules. Together with experimental protein abundances, FDM allowed us to quantify the total amount of enzymes allocated to each function. Finally, FDM enabled a genome-wide classification of the proteome according to metabolic function, and the formulation of a coarse grained model of protein allocation which quantitatively captures the global changes of the proteome across conditions.

## Results

### Functional decomposition of metabolic fluxes

*In silico* genome-scale model of metabolism (GEMs) enable the quantitative modeling of cellular metabolism by including information on thousands of cellular metabolites and reactions, as well as on the cellular biomass composition and the enzyme-reaction assignation [31]. The prototypical use of GEMs is the inference of intracellular fluxes by using a combination of empirical constraints and optimization, an approach that is generally termed Flux Balance Analysis (FBA) [15]. In brief, growing cells accumulate biomass building blocks (e.g. amino acids and nucleotides) at rates set by the cellular biomass composition and the growth rate, while also regenerating the ATP to sustain homeostasis and growth (the so-called “maintenance” energy flux). Both biomass-associated and energetic demand fluxes are fixed empirically by the biomass composition and the fluxes of metabolite uptake or excretion. Once the information have been determined empirically, intracellular fluxes can be estimated by taking the flux pattern(s) that maximize a given objective function. Such optimal flux patterns are generally in good agreement with experimental data on intracellular fluxes for carbon-limited growth [32]. Our implementation of FBA, with a detailed description of all constraints, is given in Supp. Note 1.

Given a flux pattern obtained via FBA, we sought a general method to quantify how much a given metabolic reaction contributes to another metabolic process, e.g. how much of the carbon intake flux is used for the production of a given amino acid, or how much of the flux through a given glycolytic reaction is associated to the regeneration of ATP. Answering these questions is tantamount to expressing the metabolic fluxes in the network in terms of the set of demand fluxes *J*_*γ*_ corresponding to the consumption of biomass building blocks or the production of energy in each growth condition.

It is possible to provide an explicit expression relating the FBA-derived flux vector **v**, with entries *v*_*i*_ for each reaction *i*, to the demand fluxes *J*_*γ*_. As described in depth in Supp. Note 2, the set of the demand fluxes is not arbitrary, but is determined by the set of non-dimensionless constraints in the metabolic model. Because of the linear properties of the optimization problem, and as long as the flux solution is unique, the nonzero fluxes can be expressed [21] as a linear combination of the demand fluxes *J*_*γ*_:

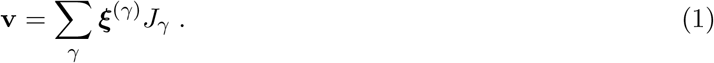

This expression represents a parameterization of the optimal fluxes in terms of the demand fluxes *J*_*γ*_, and allows to analyze how the flux **v** changes in response to perturbations of the demand fluxes. The terms ***ξ***^(*γ*)^ determine how variations in the demand fluxes *J*_*γ*_ affect each reaction. To determine these coefficients we note that they match the derivatives of the fluxes with respect to the demand fluxes. Thus, they can be obtained numerically by computing the optimal fluxes upon a small perturbation of each demand flux *J*_*γ*_. Taken to face value, Eq. (1) shows that the flux pattern **v** can be partitioned into the sum of several “flux components”

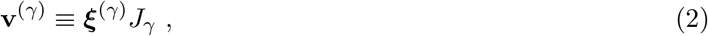

where each component **v**^(*γ*)^ satisfies the mass-balance constraints of the network and is associated to a single demand flux *J*_*γ*_. In the context of metabolic networks, the production of biomass and energy represent natural definitions of biological functions which the cell has to perform in order to survive and grow. This offers a natural biological interpretation of the linear relation between cellular and demand fluxes encapsulated in Eq. (2): it represents a *functional decomposition* of the metabolic fluxes, where each reaction *i* contributes to the function *γ* (with associated demand flux *J*_*γ*_) by a fraction 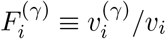 of the total flux *v*_*i*_. This is illustrated in Fig. 1A, where the flux of each active reaction (blue in the top diagram) is split into several components (in different colors in the bottom diagram), and therefore to each reaction (arrows) is assigned a breakdown into different biological functions (piecharts).

**Figure 1.**
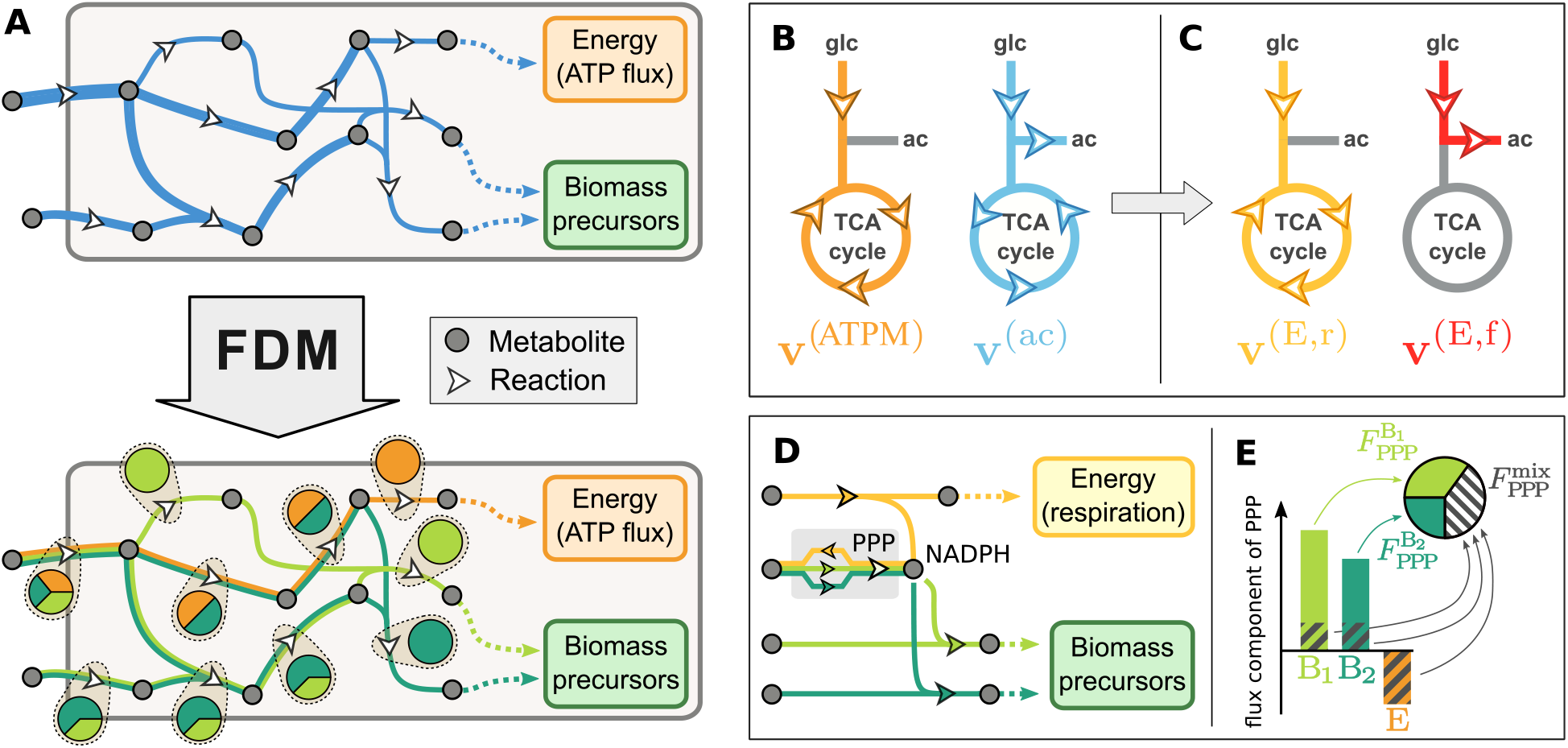
General approach to the functional decomposition of metabolic fluxes. **(A)** Functional Decomposition of Metabolism (FDM) is a method to decompose metabolic fluxes **v** (top diagram, in blue) estimated via Flux Balance Analysis (FBA) into several components **v**^(*γ*)^ (bottom diagram, colored lines). In this simple example, fluxes are split into three components: energy production (orange) or the biosynthesis of two biomass precursors (shades of green). These flux components allow to quantify for each reaction (white arrows) the fraction of flux that is associated with each metabolic function, indicated here by the pie charts. **(B)** The application of this method to *E. coli* cells grown aerobically in minimal glucose media requires overcoming two main problems. First, the presence of a constraint on acetate excretion, necessary to obtain a good agreement with experimental data, leads to an acetate-associated flux component **v**^(ac)^ (cyan) which has no clear interpretation in terms of biological functions. Second, the flux components associated to TCA reactions are found to have both positive terms, e.g. in the energy-associated component **v**^(ATPM)^ and negative terms, e.g. in the acetate-associated component. This sign mismatch does not allow us to define the functional shares of these reactions as ratios of flux components, *v*^(*γ*)^*/v* (see text). **(C)** By linearly combining the flux components associated to energy and acetate production, it is possible to define two new flux components associated to energy production via respiration (**v**^(E,r)^, yellow) and fermentation (**v**^(E,f)^, in red). These flux components have a clear biological interpretation, and do not lead to inconsistent flux directions for TCA reactions. **(D)** In some cases, sign mismatch among flux components is an inevitable consequence of the network structure. In this simplified example, NADPH is produced along with energy (by the ICD reaction in the TCA cycle) and consumed in biosynthetic reactions. In order to preserve a steady concentration of NADPH, the flux through the penthose phosphate pathway (PPP) has to adjust upon changes in energetic or biosynthetic cellular demands, thus leading to negative flux components. **(E)** A functional decomposition can be defined even in presence of negative flux components. Negative components partially “cancel out” the positive components; the magnitude of the cancellation defines a “mixed” fraction *F* ^mix^ not associated to any specific biological function. The rest of the flux components are then associated to the remainder, 1 *− F* ^mix^ (see Supp. Note 2.3 for details).

We term the definition of function-specific shares for individual metabolic reactions (and, later, metabolic enzymes) *Functional Decomposition of Metabolism* (FDM). In the next sections we will discuss the application of FDM to the concrete case of exponentially growing *E. coli* cells in carbon minimal media. However, we will first highlight two challenges that emerge when applying the method on realistic networks, and how both issues are solved by adjusting the set of biological functions used to define the functional decomposition.

### Additional constraints and mixed flux components

While the simple example in Fig. 1A illustrates the general idea of the method, it does not capture two important obstacles that prevent the straightforward application of FDM to realistic networks. Firstly, additional constraints are often needed to correctly model the cellular fluxes. Such additional constraints have to be accounted for in the flux decomposition, Eq. (2), but the biological interpretation of the corresponding flux components can be opaque. For example, acetate is excreted by fast-growing *E. coli* cells even in presence of oxygen, but FBA fails to model overflow metabolism without additional constraints. This constraint leads to the appearance in Eq. (2) of a flux component **v**^ac^ = ***ξ***^ac^*J*_ac_, proportional to the acetate production flux *J*_ac_ (Fig. 1B, cyan). However, the biological interpretation of this term is unclear, since the vector **v**^(ac)^ corresponds to the production of neither energy nor biomass components. A closer look reveals a second problem: reactions belonging to the TCA cycle presented negative entries 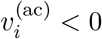 (indicated in figure by the counterclockwise arrows), in contrast to the positive direction of the overall fluxes dominated by energy production (Fig. 1B, orange, clockwise arrows). Such sign mismatch obstructs a consistent functional decomposition for the TCA reactions: a ratio 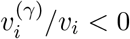 can hardly be interpreted as the share 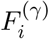 of flux associated to the function *γ*. This problem is not unique to acetate excretion, but is generically expected to arise even in absence of external constraints.

Fortunately, the linear structure of Eq. (2) allows us couple different biological functions, expressing the flux vector as a different linear combination of flux components (see Supp. Note 2). In this example, the negative components in the TCA flux can be eliminated by linearly combining the energy- and acetate-associated components into a new component 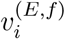 (Fig. 1C, red) which describes aerobic fermentation, leading to new vectors with no negative entries and a clear biological interpretation. The example of acetate excretion shows how negative flux components can be an indication that two metabolic activities (e.g. energy generation and acetate production) are tightly coupled, and it is more sensible to consider them as a unique process (e.g. fermentation). However, in the general case, the opportunity of coupling different metabolic processes, as well the biological interpretation of the resulting function, have to be evaluated on a case-by-case basis. Furthermore, not all sign-mismatched components can, or need, to be removed by coupling different biological functions. In fact, this phenomenon is generically expected in association with intermediate metabolites which are differently produced or consumed in association to different metabolic functions. This is illustrated in Fig. 1D where we show a simplified picture of NADPH balance in the cell. The electron donor NADPH is consumed in biosynthetic processes, and produced during aerobic respiration (by isocitrate dehydrogenase). The flux through the Penthose Phosphate Pathway (PPP) is regulated to keep the concentration of NADPH constant upon variations in the energetic and biomass-associated fluxes [33]. This particular role for PPP is reflected in the sign-mismatch among the flux components associated to energy and biomass biosynthesis. To account for such role, we introduced an additional “mixed” biological function, proportional to the amount of cancellation between positive and negative flux components:

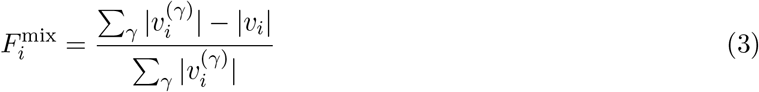

This quantity is equal to zero in absence of negative flux components, and approaches one in presence of large flux components of opposite signs. As shown in Fig. 1E, we associated the remainder of the functional share 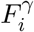 to the predominant metabolic functions (see also Supp. Note 2). This allowed us to consistently define a genome-wide functional decomposition 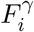 even in presence of sign-mismatched flux components.

### Application to *E. coli* growing on glucose minimal medium

To model the intracellular fluxes, we made use of the most recent *E. coli* genome-scale model of metabolism, iML1515 [11], including 2719 reactions and associations to 1515 protein-coding genes. We assembled physiological data for exponentially growing *E. coli* K-12 NCM3722 cells across conditions, including a “reference” condition corresponding to glucose minimal media, and slower growth conditions attained by either titrating the uptake of glucose (C-limitation), or by inhibiting protein synthesis using sublethal doses of chloramphenicol (R-limitation). (See Methods and Supp. Table S1 for the strains used in this study.) For each of these conditions, we used the macromolecular composition of the cell (Fig. 2A and Supp. Fig. S1), and determined the demand fluxes for individual amino acids, nucleotides, lipids and other small molecules (Supp. Fig. S1). In particular, the abundance of cysteine and glycine residues were found to be quite different compared to the values in iML1515 across all conditions (Supp. Fig. S1K). Three versions of the iML1515 models with modified biomass composition are available in Supp. File S1. For each growth condition, the metabolic model was constrained using the condition-specific biomass composition, the measured acetate excretion fluxes (Methods, Supp. File S2), Fig. 2B and the ATP maintenance flux. The latter was estimated separately for the two growth limitations by matching the minimal glucose intake allowed by the metabolic model to the measured glucose intake flux (Methods), shown in Fig. 2C. This allowed us to define maintenance fluxes specific for each of the two growth limitations explored (Fig. 2D). Finally, the metabolic fluxes were then computed with “parsimonious”FBA, by first minimizing the glucose uptake, and then minimizing the *L*_2_-norm of the fluxes, which guarantees the uniqueness of the solution (see Supp. Note 1). This approach allows to model the intracellular fluxes in *E. coli* with remarkable accuracy [34].

**Figure 2.**
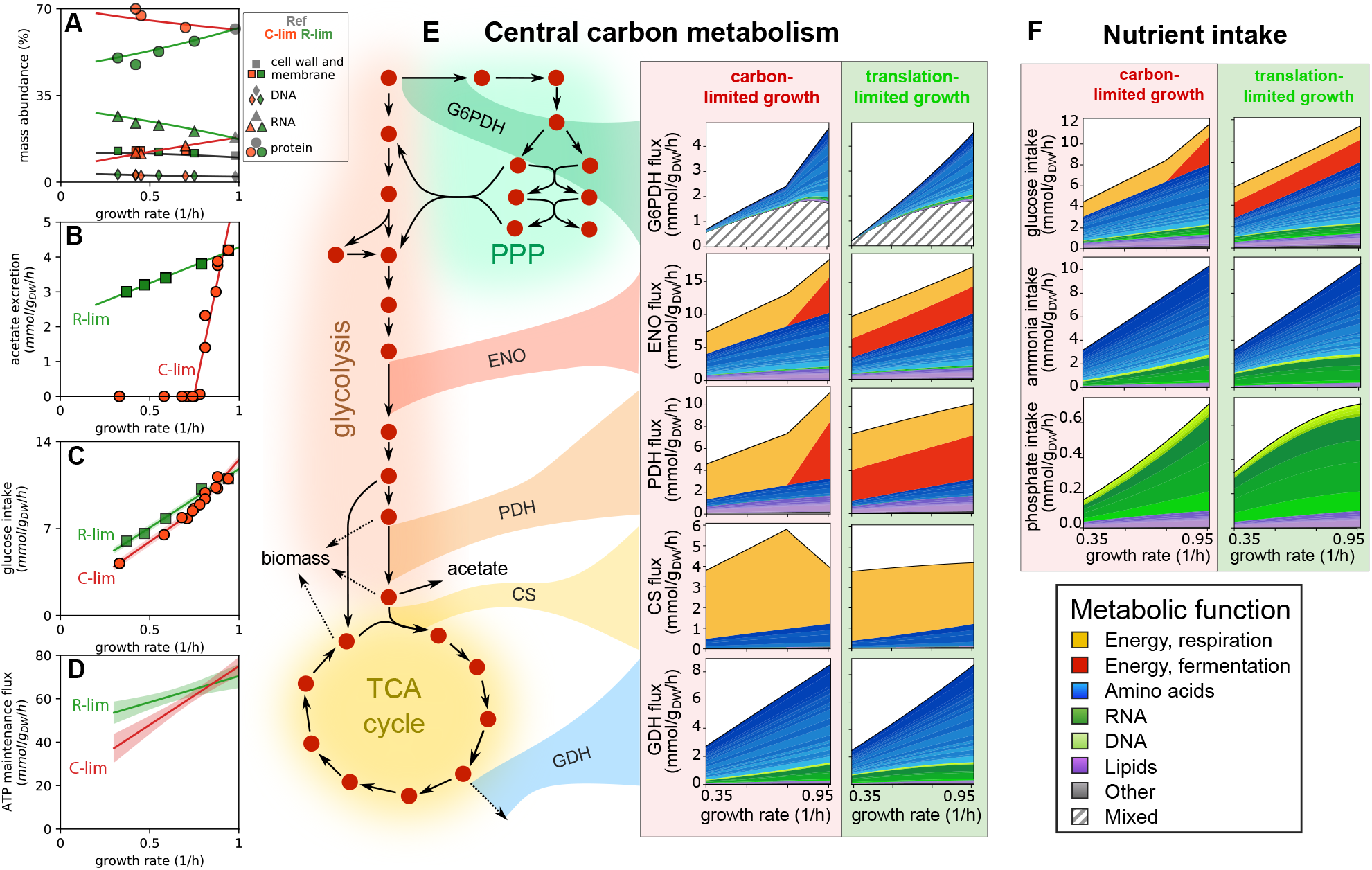
Application of FDM to carbon- and translation-limited *E. coli*. **(A)** Summary of the growth-dependent biomass composition of carbon-limited (red) and translational inhibited (green) *E. coli* cells. See Supp. Fig. S1 for a detailed breakdown of the biomass composition. **(B)** Acetate excretion rate against growth rate in C-limitation (red) and R-limitation (green). The solid lines are best fit to the data, and have been used to prescribe the acetate demand flux *J*_ac_ as a function of growth rate in the two growth limitations. **(C)** Glucose uptake rate. Filled circles and squares indicate measurements (see Methods). Solid lines indicate the flux prediction corresponding to the best fitting ATP maintenance flux parameters (see next panel); shaded areas represent 68% confidence bands. **(D)** The ATP maintenance flux is modeled as a linear function of the growth rate *µ, J*_ATPM_ = *σ*_0_ + *σ µ*. The solid lines indicate the best fits, and the shaded areas representing 68% confidence bands. The best fit values are shown in Supp. Table S2. **(E)** Flux decomposition of the central carbon metabolic pathways across growth rates for C-limited (left) and R-limited (right) conditions. The reactions shown are G6P dehydrigenase (G6PDH), enolase (ENO), pyruvate dehydrogenase (PDH), citrate synthase (CS) and glutamate dehydrogenase (GDH). **(F)** Flux decomposition of glucose, ammonia and phosphate intake fluxes.

Applying FDM, we separated the metabolic fluxes into individual components (Supp. File S3). Respiration and fermentation fluxes obtained by coupling the flux components associated to ATP maintenance and acetate production, as illustrated before (Fig. 1BC); the production of several biomass components was also found to be coupled to the ATP maintenance flux component, and the corresponding flux components were similarly combined (see Supp. Note 2.4 for details). We illustrate in Fig. 2E the flux decomposition for some representative reactions from the central carbon pathways across growth rates. As the carbon flows from upper to lower glycolysis and the TCA cycle, precursors are diverted towards biosynthetic pathways, and therefore reactions are increasingly focused toward energy production. Biosynthetic fluxes, e.g. the glutamate dehydrogenase reaction (GDH) are similar between the two growth limitations. The decomposition of the intake fluxes (Fig. 2F) reveals the fraction of nutrient sources associated to energy production and to the synthesis of biomass precursors. The functional decomposition of the carbon uptake flux mirrors that of upper/mid glycolysis (e.g. enolase, ENO): depending on the condition, 25% to 50% of the carbon is consumed for energy production (*∼*30% in reference condition), while the remainder is allocated to the biosynthesis of biomass precursors, mostly amino acids (shades of blue). Nitrogen and phosphate are instead used exclusively for the biosynthesis of biomass components, in particular amino acids (nitrogen) and nucleotides (phosphate).

As mentioned above, the flux to the penthose phosphate pathway (PPP) is mostly associated to biomass production, but has a large mixed components 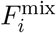, as shown for the G6P dehydrogenase (G6PDH) reaction. To explore the prevalence of sign-mismatched flux components across the entire network, we computed 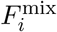 before (Supp. Fig. S2A-C) and after (Supp. Fig. S2D-F) the flux coupling procedure. Mixed functional fractions for TCA and oxidative phosphorylation reactions are greatly reduced by the coupling. Instead, large mixed functional components for reactions in the penthose phosphate pathway (PPP), upper glycolysis and nucleotide metabolism persist after the coupling. We studied the origin of such mixed components by dissecting the synthesis and consumption fluxes of each metabolite in terms of flux components (Supp. Fig. S2G-I), and identified key metabolites such as NADPH and DHAP that are produced and consumed in association with different metabolic functions, thus necessitating anaplerotic reactions with mixed functional assignation.

### Energetic metabolism

The functional decomposition of the network fluxes allowed us to study in detail the energetic metabolism of the cell. In *in silico* metabolic models, the overall ATP flux produced by the metabolic network (the “maintenance” flux) models the cellular requirements due to homeostasis and growth. Charging of tRNA and protein synthesis is generally thought to be the major energetic burden to the cell [35], at 4 ATP equivalents per residue. The estimated ATP demand is shown as green bar in Fig. 3A-C for reference and slow Cor R-limited growth conditions. The impact on the cellular budget of other ATP-consuming processes is either much smaller (e.g. mRNA turnover, at most 1.5 mmol ATP/g_DW_/h in nutrient-limited conditions [36], too small to be seen in Fig. 3), or is difficult to quantify (e.g. chemotaxis and metabolic futile cycles) [37].

**Figure 3.**
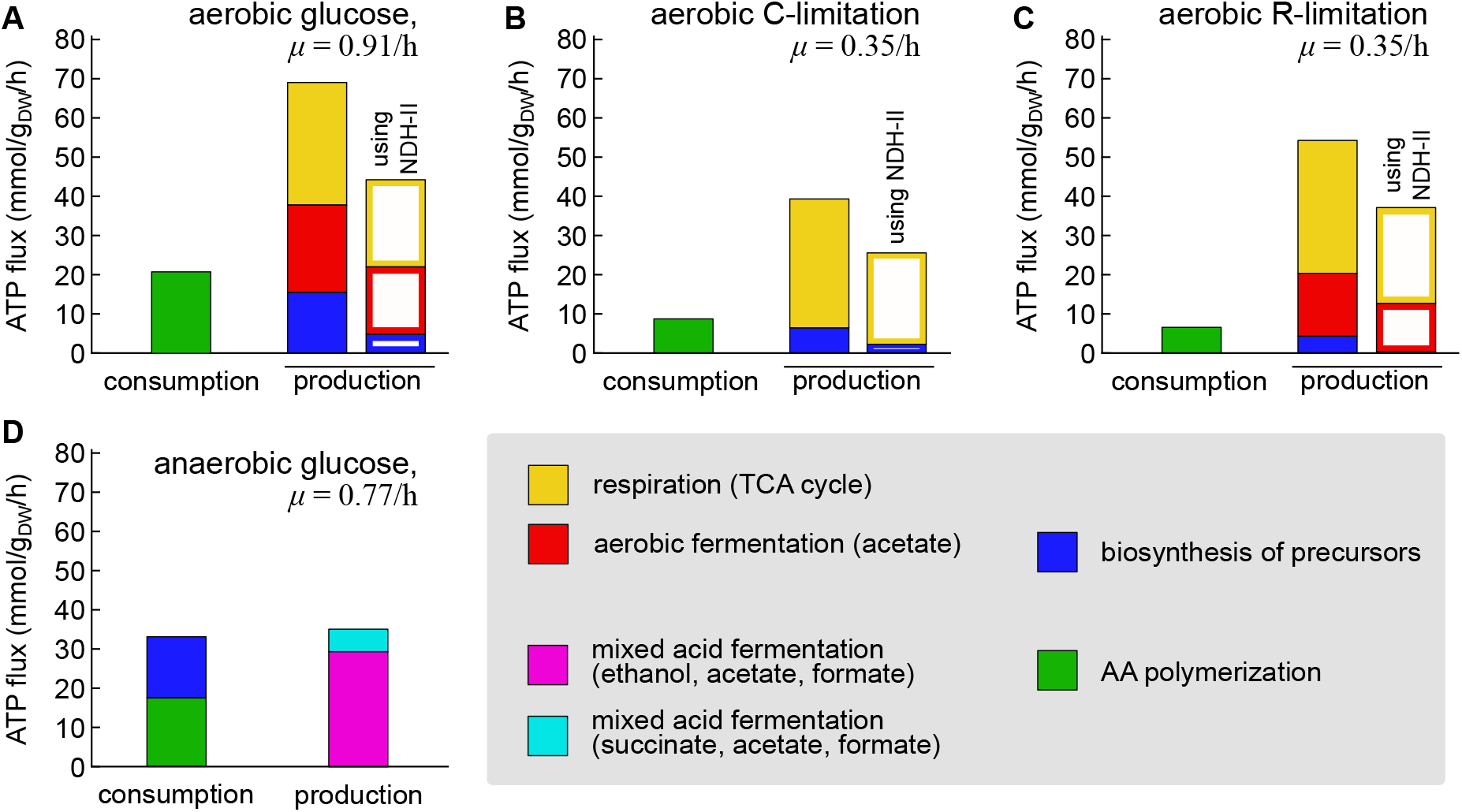
Energetic balance in *E. coli* cells. **(A)** Comparison of the cellular ATP consumption (dominated by protein synthesis, green) and production in aerobic glucose minimal medium. The ATP production is broken down according to the source: respiration (yellow), aerobic fermentation (red) and the synthesis of biomass precursors (blue) sources. Filled bars indicate the estimated ATP production using the FBA solution with maximum yield (using the efficient NADH dehydrogenase I, encoded by the *nuo* genes). Empty bars are obtained in flux solutions using the less efficient NADH dehydrogenase II (*ndh* gene; see text). **(B-C)** Same as the previous panel, but for slow, C-limited and R-limited growth. **(D)** The same breakdown of energy production and consumption was obtained for anaerobically-grown cells. Application of FBA and FDM to this case is analogous to the aerobic case, and is described in Supp. Fig. S3 and Supp. Note S2), and allowed us to determine the ATP production flux from two distinct modes of mixed acid fermentation (purple and cyan, see Supp. Fig. S3G) and the ATP consumption associated to the biosynthesis of biomass precursors (blue). The estimated production and consumption fluxes are close to each other, around 35 mmol ATP/g_DW_h.

When comparing the ATP production fluxes to the estimated consumption due to AA polymerization, we found that the total production flux was several-fold higher than the expected consumption flux (compare the filled bars in Fig. 3A-C; see also Supp. Fig. S4A). Most of the energy production, about 70 mmol ATP/g_dw_h in reference condition, is associated to the respiration and fermentation pathways (yellow and red, respectively, yielding *∼*55 ATP/g_dw_h combined). In slow, C-limited growth, only the respiration pathway is active, and the ATP flux is reduced (Fig. 3A). However, in the case of R-limitation, the energetic fluxes appear to be similar to those observed in reference condition, with a roughly equal share for flux associated to respiration (yellow) and aerobic fermentation (red) (Fig. 3B). This is due to the combination of acetate production (Fig. 2C) and high total ATP production (Fig. 2D) observed for slow-growing R-limited cells.

Counter intuitively, the biosynthesis of biomass building blocks also has a positive net contribution to the cellular energy budget. In fact, the ATP produced in conjunction with the biomass building blocks (filled blue bars) almost compensates the energetic demand associated with protein synthesis (in green). Therefore, the ATP fluxes associated to respiration and fermentation are apparently not required by known energy-consuming processes.

It is possible that the ATP production flux might have been overestimated if *E. coli* operated the electron transport chain (ETC) at reduced efficiency, which the cell can be modulate by expressing different NADH dehydrogenases [38] (Supp. Fig. S4BC). By using available quantitative proteomics data [7, 39], we found that, while the concentration of the efficient NADH dehydrogenase I (NDH-I, encoded by the *nuo* genes) is much higher than that of the inefficient NADH dehydrogenase II (*ndh* gene) in carbon-limited conditions, the two generally have opposite dependencies on the growth rate, and become comparable at either fast or slow, R-limited growth (Supp. Fig. S4D). FBA fluxes are obtained under the assumption of minimal carbon consumption, which leads to the use of the efficient NDH-I protein complex. When assuming that NDH-II is used instead of NDH-I (i.e. modeling a *nuo*^*−*^ strain), the predicted energy production fluxes are reduced by about 40%, thus partially reconciling the estimated energy production flux with the theoretical cellular demand (Fig. 3A-C, open columns; see also Supp. Fig. S4E), but still exceeding the energy consumption flux by 2-4 fold.

Energy metabolism was further examined by studying the energetic balance in anaerobic conditions, where ATP production only relies on substrate-level phosphorylation and both NADH dehydrogenase enzymes are inactive. We measured the growth rate and main metabolic fluxes of glucose-limited cells in anaerobic growth (see Methods), and used the data to model the intracellular fluxes with FBA (Supp. Fig. S3A-F). The estimated maintenance ATP flux is much lower compared to the aerobic case, and almost growth-independent at *∼*20 mmol ATP/g_dw_h. This lower value is also consistent with data from an earlier report [11] in which, however, the difference between aerobic and anaerobic growth was not emphasized.

Application of FDM to anaerobic growth is similar to that of aerobic growth, except the constraint on acetate excretion is substituted by a constraint on succinate production, leading to the presence of two distinct mixed acid fermentation functional modes (Supp. Fig. S3G). After minimizing the prevalence of mixed functional components across the network (Supp. Fig. S3HI), we obtained the functional decomposition for all fluxes in anaerobic conditions (Supp. File S5). The total ATP synthesized by the energetic pathways is about *∼*35 mmol ATP/g_dw_h in reference condition (Fig. 3D, cyan and purple), while the biosynthesis of biomass precursors consuming about *∼*15 mmol ATP/g_dw_h (in blue). Most importantly, the total production and consumption of ATP are much closer to each other compared to the aerobic case, further suggesting that *E. coli* might be operating at reduced energetic efficiency in aerobic conditions.

### Global definition of functional modules and costs

We then turned to the analysis of the global structure of the flux and functional decomposition. (From this point on, we will only consider the flux solutions obtained with the high-efficiency NADH dehydrogenase only.) We first considered the functional decomposition in reference condition, *µ ≈* 1*/*h, and performed a hierarchical clustering of the functional shares 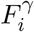. The analysis, summarized in Fig. 4A and reported in Supp. File S4, uncovered a modular structure of the metabolic network: most reactions are highly specialized, and contribute to few biological functions. Groups of functionally similar reactions can be obtained as a function of a threshold on their mutual distance (see Methods), thus allowing to define a hierarchy of functional modules in the metabolic network of *E. coli*. These functional modules were associated to the synthesis of individual (proline, histidine, arginine, lysine) or groups of amino acids (aromatic AA, alanine/valine/leucine, cysteine/methionine); these modules match well known biosynthetic pathways and superpathways [40]. As expected, TCA cycle reactions are mostly associated to energy production, while other reactions from the central carbon pathways, such as those included in upper/lower glycolysis, electron transport chain and penthose phosphate pathway, have more broadly distributed functions associated to the synthesis of several amino acids and/or energy. In sum, these results indicate that our approach is able to recapitulate the known biological functions of the metabolic network of *E. coli*.

**Figure 4.**
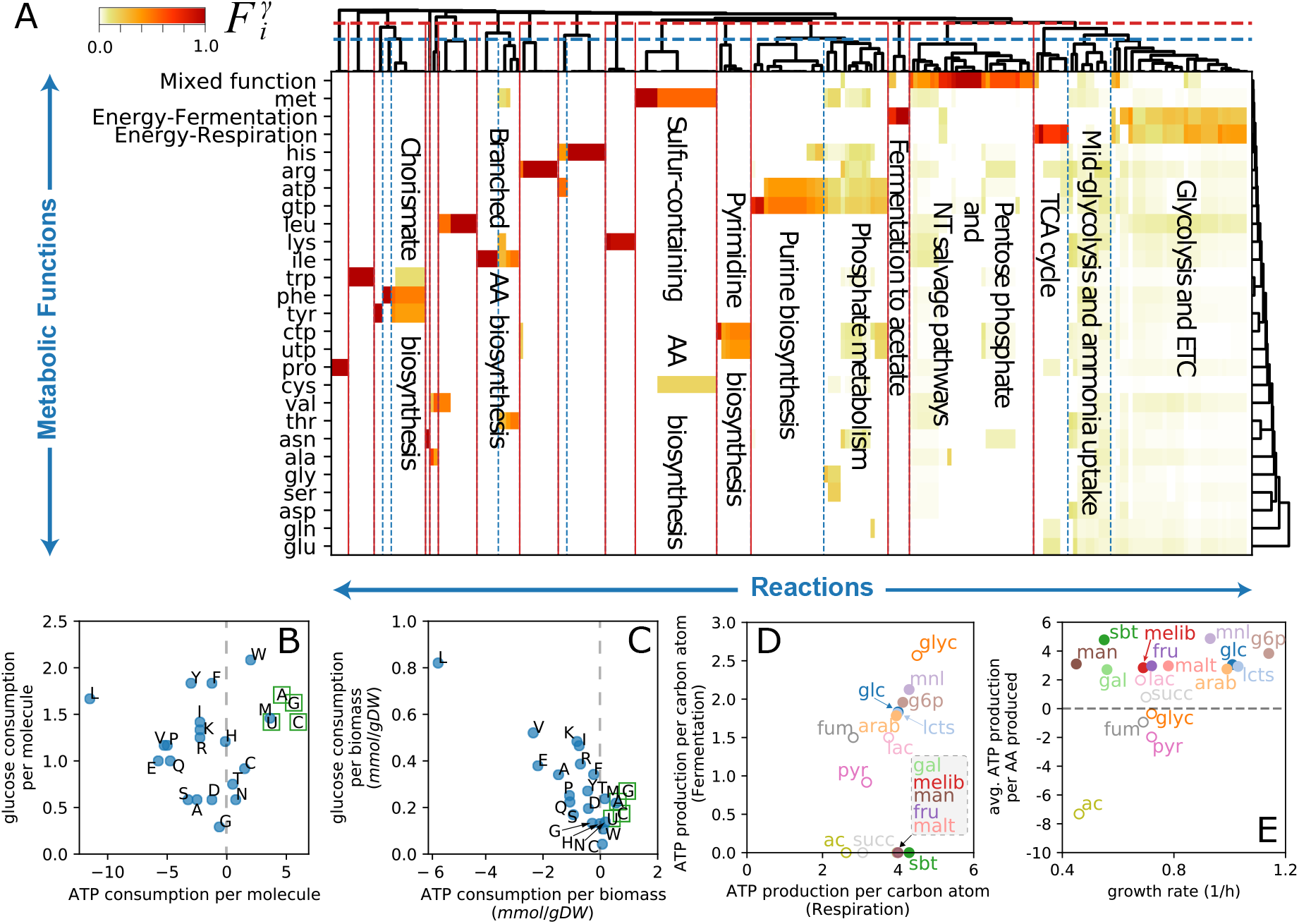
Metabolic costs for biosynthetic activities. **(A)** Hierarchical clustering of the functional de-composition 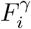, restricted to the production of energy, amino acids and nucleotides, in reference condition (214 reactions). Clustering reactions based on the similarity of the functional profiles allows to define a hierarchy of functional modules. Here we show the modules arising from two different choices of the similarity threshold (horizontal lines in the top dendrogram); the resulting functional modules are summarized in Supp. File S6. **(B)** Carbon and energy (ATP) costs associated to the biosynthesis of amino acids and nucleotides, in glucose media. Nucleotides are more expensive that most amino acids, both in terms of carbon substrate and energy. Instead, the biosynthesis of several amino acids is associated to the production of energy (i.e. a negative cost). Such coupling emerges from flux-balancing the pathway intermediates (since the flux components ***ξ***^(*γ*)^ are flux-balanced) and from the removal of sign-mismatched flux components (see Supp. Note 2). **(C)** Same as before, but weighting the costs by the demand of the building blocks (in mmol per gram of dry weight). Leucine production is associated with a large glucose uptake, but also to a large associated ATP production. **(D)** Scatter plot of respiration and fermentation ATP yields per unit of intaken carbon atom, for a variety of carbon sources (data in Supp. Files S5 and S6). The yields were calculated by repeating the FBA and FDM calculations for cells growing on different substrates, constraining the growth rate and fermentation fluxes to the experimental data from Ref. [5], and further assuming the same ATP maintenance flux parameters and growth-dependent biomass composition as for carbon-limited growth. Open symbols indicate the yields for the glycolytic carbon sources, while filled symbols indicate the yields for gluconeogenic substrates. An ATP yield of 0 is shown for the fermentation pathway if cells are not producing acetate. **(E)** The average net ATP production per AA (weighted by AA abundance) for different carbon substrates, as a function of the corresponding growth rate (data in Supp. Files S5 and S6). For most glycolytic carbon sources, 3 to 5 ATP molecules are regenerated for each amino acid synthesized. Instead, the net energy balance is close to zero, or negative (net consumption), in cells growing on gluconeogenic carbon sources such as pyruvate, fumarate, succinate and, in particular, acetate.

Each flux component, Eq. (2), describes the pathway used by the cell to synthesize either ATP or biomass building blocks, including their associated ATP production or consumption, including contributions from both the synthesis of biosynthetic precursors and due to the flux coupling (Fig. 1BC and Supp. Note 2). Using the flux components, we computed carbon and energy costs for each biomass component and, in particular, for amino acids and nucleotides. These are shown in Fig. 4BC in either costs per molecule (Fig. 4B) or per amount of cellular biomass (Fig. 4C) in glucose minimal medium, and reported in Supp. File S7. We observed that the overall ATP balance is positive for many amino acids, especially for the most represented in the biomass composition (glycine, leucine). On the other hand, nucleotides biosynthesis is expensive in terms of both energy and carbon consumption.

We validated these results in two ways. Firstly, the energetic costs associated with the biosynthetic pathways of each amino acid are well characterized [29, p.138-140]. These costs are computed for each pathway starting from carbon precursors, rather than from the external carbon source. In order to compare to these costs, we removed from the flux components all reactions belonging to the central carbon pathways, and computed the energetic balance for the biosynthetic reactions only. Reassuringly, the resulting energy cost were positive and matched well those reported in [29, p.138-140] (Supp. Fig. S5A). As a second test, we compared our carbon and ATP costs to those obtained in a previous analysis from Kaleta et al. [30]. In this study, the costs were computed either by manual counting or through optimization on a simplified *E. coli* network. The comparison displayed a general agreement for most amino acids, see Supp. Fig. S5BC. However, our approach led to much smaller ATP costs for the biosynthesis of a few amino acids compared to Ref. [30]. As illustrated in Supp. Fig. S5D, this was traced back to the different assumptions involved in the methods: while NADPH requirements were simply converted to energetic costs in Ref. [30], our methods enforces the complete flux balance of NADPH, leading to significant differences for some amino acids.

By using experimentally determined acetate production fluxes from Ref. [5], as well as the biomass composition for C-limited cells (Fig. 2A), we were able to repeat the analysis for cells grown aerobically on a variety of carbon sources. We found that the efficiency of the energetic pathways and that of biosynthetic pathways depended strongly on the substrate, as seen in Fig. 4D. The energy yields of the respiration pathways (Fig. 4D) are *∼*4 ATP per C atom for glycolytic carbon sources (open circles), 2.5 to 3 ATP/C atom for gluconeogenic carbon sources such as acetate, succinate, fumarate and pyruvate (filled symbols). The energetic efficiency of the fermentation pathway is typically 2 ATP/C atom for glycolytic carbon sources, while gluconeogenic carbon sources yields between 1 ATP/C atom (pyruvate) to *∼*2.6 ATP/C atom (glycerol).

The net energy yield associated to amino acid biosynthesis is also strongly affected by the carbon substrate on which the cells are grown Fig. 4E). Amino acid biosynthesis is on average much more expensive for cells grown on gluconeogenic carbon sources, compared to glycolytic substrates. For example, the synthesis of an amino acid on mannose produces about 3 ATP, while it consumes 7 ATP when cells are grown on acetate. (See Supp. Fig. S5E-L for a breakdown of the costs for individual amino acids and nucleotides.) Interestingly, no particular differences among glycolytic substrates is seen in either the energetic efficiencies (Fig. 4D, filled circles) nor the average ATP produced consumption per synthesized amino acid (Fig. 4E), in contrast to the wide range of growth rates achieved by cells reared on these substrates (shown as x-axis in Fig. 4E). For example, owing to their similar chemical composition, mannose and glucose have very similar energetic yields, but the growth rates for cells grown on the two substrates differ by a factor 3. Therefore, the growth rates achieved with different glycolytic substrates do not appear to be determined by differences in the energetic parameters of each carbon source. Rather, the differences in growth rates likely stem from the expression levels of the catabolic proteins [7, 41]. Similarly, the glucose intake flux for wild-type *E. coli* in anaerobic conditions is about 2 times the flux in aerobic conditions, and most of the glucose is used for producing energy (compare Supp. Fig. S3J to Fig. 2F). However, the growth rate is only reduced by less than 20%, suggesting that neither carbon nor energy are the main limiting factors for aerobic growth in glucose minimal media. Gluconeogenic carbon sources (empty symbols) tend to yield less energy and allow for slower growth rates compared to the best glycolytic carbon sources (compare e.g. succinate vs glucose), or similar to the worst glycolytic sources (e.g. acetate and mannose, both yielding growth rates close to 0.3/h).

### Functional decomposition of the proteome

The functional decomposition described above defined functional shares for each flux-carrying metabolic reaction in the cell. In turn, these functional shares can be used to generate a functional decomposition for the corresponding metabolic proteins. To do so, we made use of highly accurate experimental protein abundances [7, 39] obtained for cells grown in conditions matching those explored above, namely carbon and translational limitation. Protein abundances (in units of protein mass fractions) were obtained for a total of 2017 out of 4312 protein-coding genes in *E. coli* (Fig. 5A). After computing the functional decomposition in each growth condition, we made use of the gene-protein-reaction matrix of the iML1515 model to associate the reactions to expressed proteins. Overall, 412 proteins were both associated to flux-carrying reactions and detected in reference condition; the numbers are similar for the other growth conditions. For these reactions, the joint use of the experimental protein abundances and of the functional decomposition allowed us to quantify the contribution of each enzyme to the various metabolic functions, as illustrated in Fig. 5B for the enzyme enolase. For enzymes catalyzing multiple reactions (including various promiscuous biosynthetic enzymes such as ArgD, AspC and Ndk), we took the average functional shares of each reaction, weighted by the flux magnitudes (see Supp. Note 5 for details).

**Figure 5.**
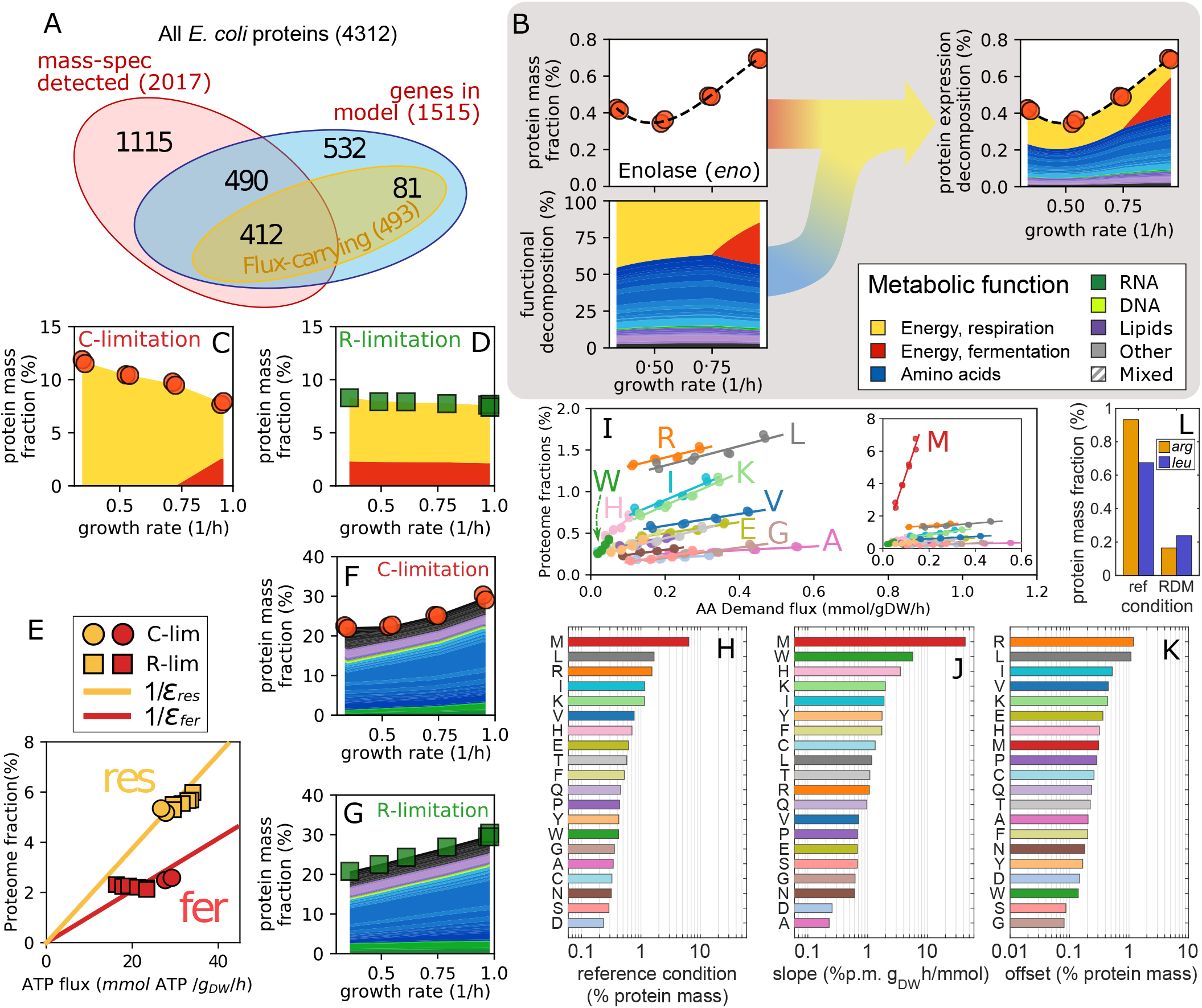
Cellular allocation of metabolic proteins across conditions. **(A)** Overlap between metabolic proteins (included in the iML1515 metabolic model), proteins associated to active reactions in reference condition, and proteins detected in the same condition [39]. **(B)** Experimental protein abundances can be matched with the reaction functional decomposition to obtain a functional decomposition for metabolic proteins. In this example we combine protein mass fractions for the Enolase enzyme in glucose-limited cells (red dots, from Ref. [39]) with the functional decomposition in the same conditions, allowing us to define function-specific protein shares (colored bands; see legend on the right). **(C)** Total abundance of energy-associated proteome *φ*_*E*_ in C-limited growth, obtained by summing over the energy-associated protein shares of all metabolic reactions. Colors indicate respiration (*φ*_E,r_, yellow) and fermentation (*φ*_E,f_, red). **(D)** Same as panel (C), but for R-limited conditions (protein data from Ref. [7]). **(E)** Protein cost of the respiratory (yellow) and fermentative (red) pathways. Solid lines are best fits lines passing through the origin; the slopes indicate the inverse of the protein efficiencies of the two pathways. We obtained *ε*_*res*_ = 5.33 *±* 0.15 and *ε*_*fer*_ = 9.62 *±* 0.37, in units of ATP flux (mmol/g_DW_h) per percent of allocated proteome. **(F)** Total proteome mass fraction associated to the biosynthesis of biomass building blocks in C-limited conditions, obtained in the same way the energy-associated proteome fraction was computed in panel (C). Colors indicate individual building blocks. **(G)** Same as panel (F), but for R-limited conditions. **(H)** Protein mass fractions associated to each amino acid in reference condition. **(I)** As the growth rate is varied in C-limited conditions, the protein shares allocated to the biosynthesis of each amino acid (symbols) are well described by linear function of the demand flux of the amino acid (solid lines indicate best fits). The inset shows the protein mass fraction associated to methionine biosynthesis, which is dominated by MetE. Letters indicate a few amino acids; colors match those used in the bars in panel (H). Similar linear relations are also observed in R-limitation (Supp. Fig. S8C); the values of slopes and offsets (y-intercepts) for both limitation series are reported in Supp. File S7. **(J-K)** Slopes and offsets (y-intercepts) of the linear relations shown in panel (I). **(L)** Protein mass fractions in reference condition and in rich defined medium (from Ref. [39]) of the biosynthetic enzymes for arginine and leucine (*arg* and *leu* genes, respectively). Levels in rich media are much lower than in glucose minimal media, as opposed to the lack of change observed in C-limitation for these two amino acids (see panel I).

### Protein costs associated with energy production

The protein fraction associated with energy production in carbon-limited growth is similar to previous estimates [5, 21], increasing from about 8% to 12% of the total proteome (Fig. 5C) and switching from a mix of respiration and aerobic fermentation at fast growth, to respiration only at slow growth. Given the decreased energetic flux in slow, carbon-limited conditions (Fig. 3AB and Supp. Fig. S4A), the overall energetic efficiency of the cell (ATP flux per unit of invested proteins) decreases at slow growth (Supp. Fig. S6A, red circles). The observed patterns are quite different in R-limited cells: the proteome shares allocated to energy production via either respiration or fermentation are mostly independent on the growth rate (Fig. 5D), mirroring the lack of change observed for the energy flux (Fig. 3B). As a consequence, the efficiency of the energetic pathways is mostly constant across growth rates (Supp. Fig. S6A, green squares). The protein efficiency for the respiration and fermentation pathways can be obtained by comparing the associated protein shares to the corresponding ATP fluxes. Respiration pathway requires about twice the proteins associated to aerobic fermentation (Fig. 5E), consistently with previous analysis [5, 21].

### Protein costs associated to biomass production

Globally, the protein fraction associated to biosynthetic activities (Fig. 5FG) ranges between 20 and 30%, and decreases as growth is slowed down. The protein cost is dominated by amino acid synthesis (shades of blue). The biosynthesis of the other components (RNA, DNA, lipids, cofactors) only makes use of a small fraction of the total proteome, less than 10% in total. In reference condition, synthesis of methionine was by far the most expensive process, requiring more than 6% of the total proteome mass (red bar in Fig. 5H). Most of this cost is due to the highly inefficient enzyme homocysteine methyltransferase (MetE). Instead, the relatively small production costs of proline, glutamine and glutamate were the only ones dominated by the shared central carbon pathways (Supp. Fig. S7A). We looked at the relationships between enzyme abundance and demand flux for individual amino acids. Across conditions, the protein mass fractions associated to each amino acid and nucleotide scale remarkably linearly with the corresponding demand flux, with more proteins being allocated in presence of higher fluxes (Fig. 5I, Supp. Fig. S8A-D). These results are consistent with linear relations observed for transcriptional reporters [3] or for protein “sectors” [4, 7] in *E. coli*, and more recently in yeast [42]. The slopes and y-intercepts (“offsets”) of the linear relations (Supp. File S7), are important parameters that reflect the overall efficiency of the pathway [4, 25] and the capacity of the pathway to rapidly change flux in dynamic conditions [39, 43], respectively. The slopes and offsets are summarized in Figure 5JK. Overall, the protein offsets were similar (at most a 25-fold difference, with 16 out of 20 in the range 0.15% to 0.35%) for most amino acids, while slopes varied over a broader range (more than two orders of magnitude). Methionine (M) biosynthesis (inset in Fig. 5I) is by far the most expensive process across all conditions, and had the steepest slope among all amino acids (Fig. 5J). Other expensive processes are the biosynthesis of tryptophan (W) and histidine (H), as indicated by their large slopes (Fig. 5J). On the other hand, leucine (L) and arginine (R) stand out as the amino acids whose associated proteome had the largest offsets (Fig. 5K) and small slope across conditions in C-limitation. In contrast, the biosynthetic enzymes are expressed at very low levels in rich media (Fig. 5L). This suggests that glucose limitation specifically impacts the expression of biosynthetic enzymes for arginine and leucine, possibly indicating that their biosynthesis becomes growth-limiting in slow, carbon-limited growth.

Overall, results for R-limited growth were similar to those obtained for C-limitation (Supp. Fig. S8A-D), with the notable exception of methionine which presented a much larger protein offset (Supp. Fig. S8EF). Protein costs for the synthesis of purines was 3 to 4 times that of pyrimidine (Supp. Fig. S8BD), and larger than most amino acids other than methionine, although the overall allocation towards nucleotide synthesis is small compared to the amino acid (Fig. 5FG, compare green to blue), due to the much smaller number of nucleotides compared to amino acids. We observed that for most amino acids, the slopes of the protein-flux relationships were slightly more steep in R-limitation compared to C-limitation, while the opposite was true for nucleotides. Such small systematic patterns were mostly due to changes in the flux through the central carbon pathways (glycolysis and TCA cycle), which influence how the corresponding proteins are allocated (Supp. Fig. S7B through D). Slopes were tightly correlated to the ratio of allocated protein and demand flux in reference condition, related to the so-called “effective” turnover rate of the pathway, and presented weaker correlations with other quantities such as the carbon costs and demand flux of each amino acid (Supp. Fig. S9).

### Coarse graining of the proteome according to metabolic function

The functional decomposition described above defined functional shares for each flux-carrying metabolic reaction in the cell. As summarized in Fig. 6A, this corresponds to about 40% of the proteome in reference condition, of which 7% is associated to energy production (“energy” sector) and 33% to the biosynthesis of biomass precursor (“biomass” protein sector). About half of the proteome is not associated to the metabolic model, and an additional 10% is associated, but the corresponding reactions carry no flux. We associated these proteins to specific cellular functions via an iterative GO-terms based categorization. By sequentially selecting the GO-terms associated with the largest protein shares, we were able to categorize *>*90% of the proteome mass using *∼* 50 GO-terms, Fig. 6B. In order to simplify the description of the proteome, we grouped all translation-associated GO-terms into a “translation” sector; similarly, GO-terms associated to nutrient transport and catabolism, as well as motility, were grouped into a single “foraging” protein sector; the remainder of the GO-terms were grouped into a “housekeeping” sector. Together with the metabolic “energy” and “biomass” sectors, these define a five-way function-based classification of the proteome.

**Figure 6.**
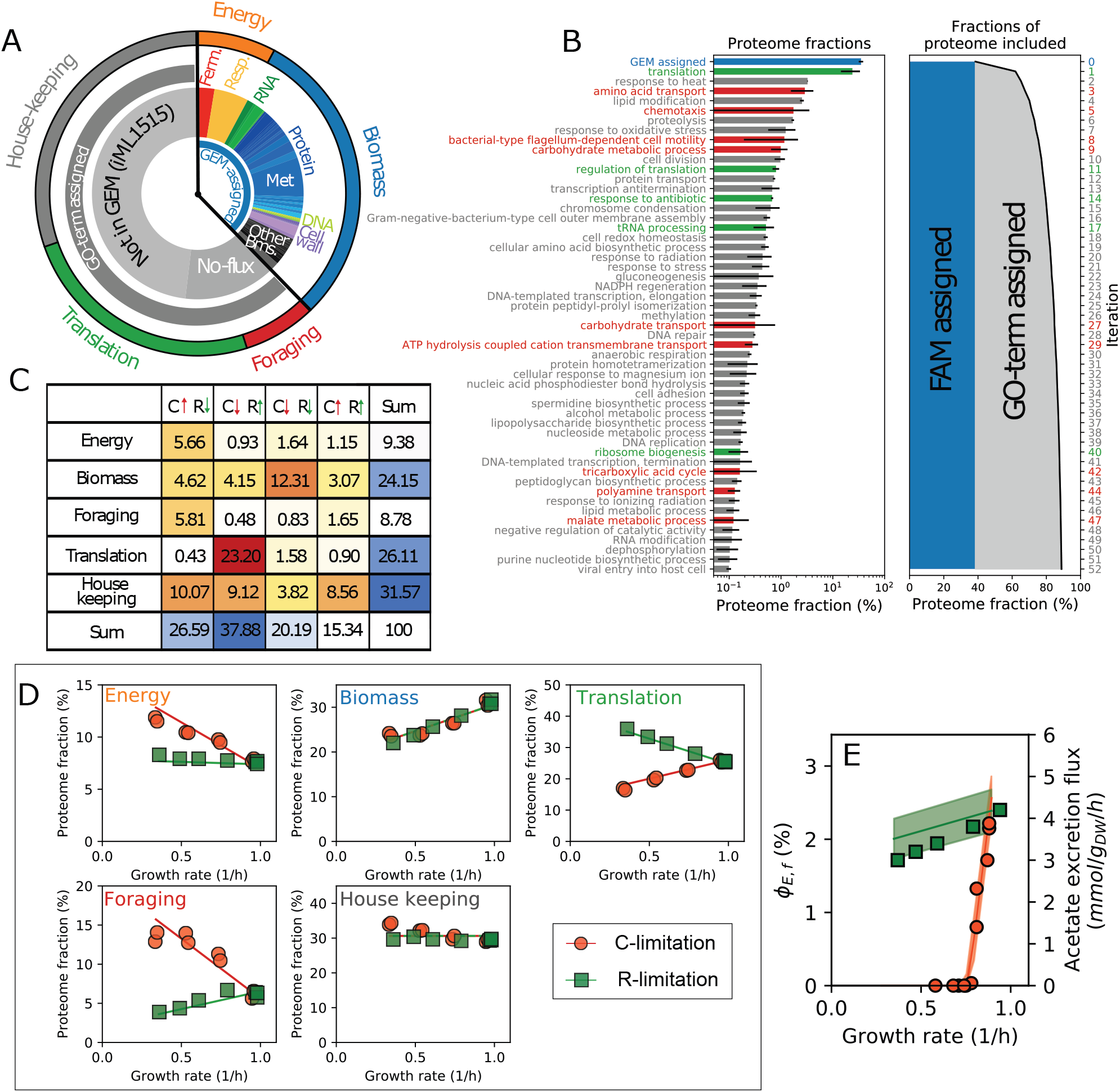
Function-based global classification and coarse-graining of the expressed proteome. **(A)** Summary of the functional decomposition of *E. coli* proteome in glucose minimal medium. The fraction of the proteome contributing to a defined metabolic function (GEM-assigned) is about 40%, with proteins involved in the biosynthesis of cellular building blocks (“biomass” sector) about 32%, and energy-producing proteins (“energy” sector) close to 8%. The remainder of the proteome is divided between non-metabolic proteins (e.g. ribosomal or motility genes), and proteins associated to zero-flux reactions (e.g. nutrient transporters). Using a GO-term enrichment analysis (see next panel) these were assigned to three sectors associated to “translation”, “foraging” (motility and nutrient assimilation) and “house-keeping”. **(B)** An iterative GO-term assignment procedure (see Methods) was used to assign specific biological functions to non-metabolic proteins. At each steps, the GO-term associated to the largest proteome share is selected, and the corresponding genes are assigned exclusively to that GO-term. Bars indicate the average proteome fractions of each GO-term across conditions, with error bars representing their variability; colors indicate the three coarse-grained sectors indicated in the previous panel. **(C)** For each protein in the five functional sectors, we summarized changes in protein levels with a binary classification [4] describing whether the protein is up- or down-regulated in C- or R-limitation, as indicated by the arrows. For all sectors except the “housekeeping”, most of the proteome (here corresponding to reference condition) is associated to only one of the four possible combinations, indicating that most proteins are consistently regulated across conditions. **(D)** Protein mass fraction associated to the five functional sectors across growth rates for C-limited (red circles) and R-limited (green squares) growth. Solid lines indicate the best fit for a coarse-grained model of protein allocation. **(E)** Using the calculated energy production fluxes and protein efficiencies for the respiration and fermentation pathways, as well as the modeled total protein abundance, the coarse grained model predicts the partitioning between respiration and fermentation-associated proteins (see Supp. Note 6). Here we show the predicted allocation towards fermentation proteins and the associated acetate excretion flux (solid lines; bands represent error propagated from the estimated errors for the efficiencies), as well the experimentally determined acetate excretion fluxes (same symbols as in the previous panel), for C-limited and R-limited growth.

We first assessed whether proteins within the same functional sector are similarly regulated across conditions. Each protein was previously classified depending on their change in protein abundance upon carbon starvation or translation limitation [7]. Hence, we were able to break down each functional sector into four different components, corresponding to the four possible combinations of up- and down-regulation in the two growth limitations (Fig. 6C; see also Supp. Fig. S10A-C). For most functional groups, proteins appear to be consistently regulated across conditions, with the vast majority of the associated proteome belonging to only one of the four regulatory groups. This is most evident for the “translation” functional group, where the vast majority of proteins are similarly upregulated in translation-limited conditions and downregulated in carbon-limiting conditions. Most proteins in the energy and foraging groups show the opposite behavior, i.e. are upregulated in carbon-limited conditions and downregulated in translation-limited conditions. Proteins in the “biomass” sector are downregulated in both growth limitations. Finally, housekeeping proteins are regulated more hereogeneously. This is also expected given the wide variety of GO-terms associated to the proteins in this functional group. Similar results are obtained by considering a more detailed binary classification based on three (rather than two) different growth limitations (Supp. Fig. S10D through G). We then computed the protein abundance of the functional sectors as a whole, which can be seen plotted against the growth rate in Fig. 6D. The growth-dependence of the sectors recapitulates what was seen at the level of individual proteins in Fig. 6C: The energy and foraging sectors increase strongly in carbon-limited conditions, while the translation sector is upregulated in R-limited conditions. Allocation towards the “biomass” sector is proportional to the growth rate, while the housekeeping sector changes little across conditions.

The trends observed for the protein sectors, as well as the observed switch between respiration and fermentation, can be captured quantitatively by a single phenomenological model of protein allocation. The protein mass of each sector changes according to two parameters, the quality of the carbon source *ν*_*C*_ and the translational capacity of the cell *ν*_*R*_. These two parameters determine a “phase space” of cellular growth [2]: changes in either parameters leads to simultaneous changes in the cellular growth rate *µ* and in the size of the protein sectors. The expression of the “foraging” (including a variety of transporters and chemotactic proteins) and “translation” sectors respond specifically to changes in the carbon quality and in the translation capacity, respectively. Expression of the “biomass” sector is linearly related to growth rate, while the “housekeeping” sector is constant. Finally, we assumed a regulatory constraint setting the energy sector in response to carbon starvation and translational limitation of the following form:

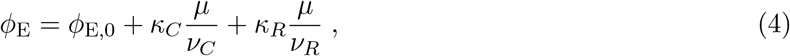

where *κ*_*C*_ and *κ*_*R*_ determine change in protein allocation in response to the specific growth limitation. As detailed in Supp. Note 6.3, these two term arise from the reduced efficiency of the energetic pathways in carbon-limited conditions (Supp. Fig. 6A), and from increased energetic demands per unit of biomass in translational-limited conditions (Supp. Fig. S6B). The fitted model (solid lines in Fig. 6D) correctly recapitulates the experimental data. Because the model explicitly accounts for the protein share associated to energy production, it can also account for the impact of protein allocation on acetate overflow. Assuming that the respiration and fermentation-associated components of *φ*_*E*_ generate an ATP flux in proportion to their proteome share (using the efficiencies determined in Fig. 5G). Without changing any of the parameters leading to the fit in Fig. 6D, the model predicts a sharp decrease in the protein share associated to fermentation *φ*_*E,f*_ in carbon-limited growth, and sustained fermentation in R-limited growth (Fig. 6E, solid lines), in agreement with the experimental acetate excretion fluxes (Fig. 6E, symbols). Thus, the model is able to simultaneously capture the observed protein allocation patterns and the metabolic switch between respiration and fermentation.

## Discussion

In this work, we presented a computational method for studying cellular metabolism and protein allocation based on the decomposition of metabolic fluxes into distinct functional components. This framework, termed *Functional Decomposition of Metabolism* (FDM), integrates a wide variety of physiological, flux and protein data using genome-scale metabolic models, and hence provides a general avenue to the analysis of complex multi-omics datasets (as summarized in Supp. Fig. S11). FDM allowed us to comprehensively evaluate metabolic costs and protein burdens associated to each metabolic functionality, a feat impossible to achieve from the analysis of single reactions or protein abundances because of the deeply interconnected nature of metabolic networks. The functional components and the costs, reported in Supp. Files S3 and S7, will be a valuable resources for model-building and for understanding the physiology of *E. coli*.

The flux decomposition on which FDM is based upon is fundamentally based on mathematical properties of FBA solutions. Furthermore, it does not rely on parameters such as kinetic constants, nor requires simplifying hypothesis on the structure of the metabolic network: FDM can be generally applied to any network, as long as the application of FBA (or other optimization approaches) estimates correctly the intracellular fluxes. At the core of FDM is a quantitative definition of system-level *metabolic functions* for each reaction and protein. Each of these metabolic functions can be associated to multiple activities, e.g. synthesis of amino acid and ATP production, when these are tightly coupled. Furthermore, flux constraints are often applied to the network in order to improve the agreement with experimentally determined fluxes; these constraints are fully accounted for by the flux decomposition, at the cost of introducing associated metabolic functions. This can seen in Fig. 1BC, where the prescribed acetate excretion flux is associated to the aerobic fermentation flux.

A long-standing puzzle in bacterial energetics is the fact that bacteria do not always utilize energy efficiently, but instead often “waste” it through various energy-spilling processes [44, 45]. The determination of carbon and energetic fluxes associated to the biosynthesis of cellular components showed that for aerobic growth on glucose, the ATP flux produced as by-product of biosynthesis is already sufficient to supply for the need for amino acid polymerization, the single biggest ATP expenditure for growing cells (Fig. 3A-C). Furthermore, specific ATP and carbon costs for biosynthesis are generally uncorrelated with the rate of aerobic growth (Fig. 4DE), suggesting that the energy-production pathways are not growth-limiting in aerobic conditions. Together, our results raise fundamental questions on the metabolic purposes of energy biogenesis by respiration and fermentation, which comprise 30% of the total carbon flux during aerobic growth on glucose and even larger fraction for slower growth, and are apparently not needed.

Indeed, the opposite changes observed in the levels of the two NADH dehydrogenases NDH-I and NDH-II across conditions (Supp. Fig. S4) suggest that *E. coli* is able to decouple electron transport from ATP production, thus reducing the production of ATP and bringing it closer (but not equal) to the estimated consumption. In anaerobic conditions, when the electron transport chain is not used, the predicted ATP production matches the costs (Fig. 3D). These results suggest that the energetic pathways in fast-growing *E. coli* cells in aerobic conditions operate with and efficiency far from the maximum allowed by the biochemical constraints. For cells grown in R-limited conditions, we observed high, constant energy production fluxes for both the respiration and fermentation pathways, despite a predicted reduction in ATP demand at slow growth. Such constant flux is accompanied by a constant share of proteome allocated to energy production, irrespective of the growth rate. This might suggest the presence of additional energy-consuming processes in slow, R-limited growth, e.g. additional ribosome turnover due to biogenesis defects [46]. Alternatively, the flux of carbon substrate towards energy production might be set by the carbon availability (which is constant in R-limited conditions), while being independent on the actual energetic demand; the latter could be matched instead by modulating the ATP yield of the energetic pathways, in agreement with the analysis discussed above.

Combining FDM with proteomics data allowed us to calculate the proteome costs associated with the *de novo* biosynthesis of each cellular component, including not just the contributions from the curated pathways, but also the prorated cost of carbon/nitrogen uptake and energy biogenesis needed for biosynthesis. For the biosynthesis of amino acids and nucleotides, we found the total abundance of the allocated proteins to be linearly increasing functions of the growth rate under carbon catabolic limitation. In addition to the well-known large cost of methionine biosynthesis, we found large protein reserves at slow growth for the production of several amino acids, particularly leucine and arginine. These large protein reserves might indicate that the synthesis of these two amino acids becomes growth-limiting in poor carbon conditions.

The whole-proteome, function-based model of protein allocation enabled by FDM is a step forward in the quantitative modelling of bacterial protein allocation. Our work allowed us reconcile two distinct classes of protein allocation models. The first class includes models based on regulation-based protein sectors [4, 7]. In these models, protein sectors are defined based on protein expression patterns, but they do not always correspond to unique biological functions. Models in the second class are formulated with function-based sectors [5, 21], and have a narrower scope (e.g. focusing on energetic metabolism). The design of a systematic procedure to functionally classify all of the expressed proteome across conditions (Fig. 6A-B), and the finding that most proteins within each condition-dependent protein sector were consistently regulated across conditions, enabled us to build a quantitative model of protein allocation bridging the two model classes.

Still, the regulation of the energy sector in poor carbon sources is only accounted for by an effective constraint, and linking its share to the underlying regulatory processes is an open problem. The increase in the protein share assigned to energy production might be a regulatory strategy to more efficiently divert flux from biosynthetic activities, or to prepare for a switch to gluconeogenic substrates [47]. In either case, fully explaining the observed patterns likely requires including information on the concentrations of metabolites and the kinetics of the respiration pathway [48, 49].

Ultimately, the wide range of analysis enabled by FDM makes it a general and powerful tool for the bioengineering and systems biology communities. The systematic evaluation of yields and costs for the production of individual metabolites enabled by FDM has natural applications to the study of microbial cell factories in which the production of a metabolite of interest is maximized. Furthermore, FDM allowed to uncover a hierarchy of functional modules in cellular metabolism without supervised knowledge on curated metabolic pathways. Thus, FDM could facilitate the rational design of heterologous metabolic pathways with varying degrees of coupling to other metabolic activities [50]. Consistently with the modular structure of bacterial metabolic networks [51, 52], the functional patterns exhibited by the reactions displayed a rich structure (Fig. 3A), which provides the quantitative counterpart of known biochemical pathways. Thus, the definition of functional shares for each reaction represents a simple alternative to other system-level approaches to the analysis of metabolic function, e.g. based on the exhaustive enumeration of extreme pathways or elementary modes [53].

## Methods

### Experimental methods

#### Bacterial strains

All strains used in this work are derived from *E. coli* K-12 NCM3722 [54–56] and listed in Supp. Table S1.

#### Growth media

The growth media for aerobic growth is the MOPS-buffered minimal medium from Ref. [5]. The phosphate-based growth media used for anaerobic growth and for other control samples in aerobic conditions contained 10 mM glucose, 80 mM K_2_HPO_4_, 20 mM KH_2_PO_4_, 10 mM NaCl, 10 mM NH_4_Cl, 0.5 mM Na_2_SO_4_, a phosphate buffer and a 1000x micronutrient solution. The 1000x micronutrient solution contained 20 mM FeSO_4_, 500 mM MgCl_2_, 1 mM MnCl_2_ ·4 H_2_O, 1 mM CoCl_2_ ·6 H_2_O, 1 mM ZnSO_4_ ·7 H_2_O, 1 mM H_24_Mo_7_N_6_O_24_ ·4 H_2_O, 1 mM NiSO_4_ ·6 H_2_O, 1 mM CuSO_4_ ·5 H_2_O, 1 mM SeO_2_, 1 mM H_3_BO_4_, 1 mM CaCl_2_, and 1 mM MgCl_2_ dissolved in a 0.1 M HCl solution. Carbon limitation was implemented by titrating 3-methyl-benzylalcohol (3MBA) concentration in strains NQ1243, NQ1448, and NQ1554 as well as by growing strain NQ1261 (*ptsG* deletion) in glucose. Translational limitation was obtained by adding sublethal doses of chloramphenicol. Concentrations of nutrients, 3MBA and chloramphenicol in each experiment are reported in Supp. File S2.

#### Growth measurements

Growth measurements for aerobic culture were performed as in Ref. [3]. Briefly, exponential cell growth was performed in a 37 ^*°*^C water bath shaker at 240 rpm. Cultures were grown in the following three steps: seed culture, pre-culture, and experimental culture. Cells were first grown as seed cultures in LB broth for several hours, then as pre-cultures overnight in an identical medium to the experimental culture. Experimental cultures were started by diluting the exponentially growing pre-culture to an optical density at wavelength 600 nm (OD_600_) of *∼*0.01–0.02. Growth rates were calculated from at least seven OD_600_ points within a range of OD_600_ of *∼*0.04–0.4.

Anaerobic growth was performed similarly to aerobic growth with a few exceptions. All transfers were performed with disposable syringes to avoid oxygen contamination. Aerobic seed cultures were diluted into Hungate tubes for preculture. After overnight growth, the precultures were diluted into fresh Hungate tubes for experimental culture. To avoid atmospheric exposure from removing samples, OD measurements were performed with a Thermo Genesys 20 modified to hold Hungate tubes in place of cuvettes. The culture temperature was kept stable during OD measurements by removing and replacing the Hungate tubes from the water bath shaker within 30 seconds. The OD_600_ measured through the Hungate tubes was equivalent to the OD_600_ measured through a cuvette for the range of 0.04-0.5.

#### Metabolite measurements

Metabolites were prepared and quantified as in Ref. [57]. Four samples of 200 *µ*L were pipetted from culture tubes at regularly spaced ODs during exponential growth. For anoxically grown cultures, samples were removed with tuberculin syringes inserted into the rubber stopper. Samples were then transferred to 0.22 *µ*m nylon filter centrifuge tubes (Corning Costar Spin-X Centrifuge Tubes) and quickly filtered by centrifugation. Samples were then stored at -20 ^*°*^C until HPLC analysis, which was performed using the Rezex RoA (H+) organic acid column with 10 mM H_2_SO_4_ as the mobile phase.

### Computational and numerical methods

#### Calculation of FBA solution and numerical derivatives

The formulation of the FBA and FDM optimization problems is described in detail in Supp. Note 1. The FBA calculations were performed in Python using CVXPY [58, 59] and the GUROBI solver. In order to estimate the numerical derivatives reliably, we set the following GUROBI parameters: maximum iteration to 1000, “BarConvTol” and “Bar-QCPConvTol” to 10^*−*12^, “FeasibilityTol” and “OptimalityTol” to 10^*−*9^. Details on the implementation of FBA and of the functional decomposition are provided in Supp. Notes 1 and 2, respectively. Additional analysis were performed in Matlab.

#### Hierarchical clustering

The hierarchical clustering in Fig. 4A was performed on reactions whose fluxes were larger than 10^*−*4^ mmol/g_DW_h and only considering the metabolic functions associated to energy production, biosynthesis of amino acid and nucleotides, plus the “mixed” functional component; clusters were determined based on the cityblock distance among the functional shares 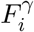. Full results are reported in Supp. File S4.

#### GO term-based decomposition

We used the “biological process” terms to define the biological functions of *E. coli* proteins which were not categorized using the functional decomposition (Fig. 6B), as follows. We considered the average protein mass fractions between reference condition and extreme C/R-limitations. The GO-term associated with the largest protein mass fractions was identified, and the genes associated to that GO-term were assigned to the corresponding biological function. This process was iterated on the remainder of the genes until the largest protein mass fraction for each GO-term was less than 0.1%.

#### Fit procedures and confidence bands

Best fit parameters for the ATP maintenance flux were obtained by minimizing the squared residuals between the experimental and modeled glucose intake fluxes, 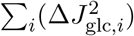 where *i* indicates the samples. The best fit parameters for the coarse grained model, including the values of *ν*_*C*_ and *ν*_*R*_ in each growth condition, were obtained by minimizing the sum of two terms. The first term is the sum of squared residuals between the protein mass fractions of the five protein sectors *α* and the modeled mass fractions 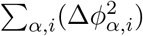; the second term are the squared residuals between modeled and experimental growth rates for each condition 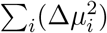. The overall function to be optimized takes the form 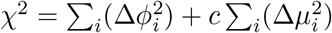, where the scale factor *c* is needed to compare the two terms since they have different physical units. We chose *c* = 0.1h^2^ so that the residuals of the growth rates are close to the typical experimental uncertainty on the growth rates (0.02/h or less).

## Data availability

The Python code necessary to run the functional decomposition on a given metabolic model is available on Github (https://github.com/ahoiching/FDM). Supplementary Files S1 to S7 are not included in this pre-print, and are available upon reasonable request.

## Author contribution

Conceptualization: MM and TH. Methodology: MM. Software: CC and MM. Validation: CC and MM. Formal analysis: MM. Investigation: BT and HO (growth and flux measurement), CC and MM (data analysis). Data curation: CC and MM. Writing - original draft: MM. Writing - review and editing: MM and TH. Visualization: CC and MM. Supervision: MM and TH. Project administration: MM. Funding acquisition: TH. All authors reviewed the results and approved the final version of the manuscript.

## Acknowledgements and funding

MM thanks Andrea De Martino, Enzo Marinari, Cong Trinh and Martin Lercher for discussions. The work was supported by NSF grant no. MCB-1818384 and NIH grant no. R01GM095903 through TH.

