## Supplementary Information for "Functional Decomposition of Metabolism allows a system-level quantification of fluxes and protein allocation towards specific metabolic functions"

#### Supplementary Material

Matteo Mori *†[1], Chuankai Cheng *[2], Brian Taylor[1], Hiroyuki Okano[1], and Terence Hwa[1]

[1]Department of Physics, University of California San Diego, 9500 Gilman Dr. La Jolla, CA 92093, USA

[2]Department of Biological Sciences, University of Southern California, Los Angeles, California, USA

---

* These authors contributed equally to this work.

**Supplementary Figures** **47**

**Supplementary Tables** **62**

**Supplementary Files** **64**

**Supplementary References** **65**

#### Supplementary Note 1: Constraint-based modelling of *E. coli* metabolism

In this note, we describe our procedure to model the metabolic fluxes of *E. coli* in glucose minimal media which constitute the basis of the flux and functional decomposition procedure (described in Supp. Note 2). This note is divided in three sections. First we will describe the definition of the FBA optimization problem, including additional flux constraints. Second, we will describe the definition of a condition-dependent biomass composition for *E. coli* in the growth conditions considered in this work. Third, we will describe the numerical implementation of the FBA optimization problem.

##### 1.1 Definition of the FBA optimization problem

Flux balance analysis (FBA) is the most widely used framework within the family of constraint-based modeling (CBM) approaches. In short, FBA allows to estimate metabolic fluxes by finding the optimal flux patterns maximizing a given objective function (a linear combination of the fluxes), subject to a set of linear constraints. In this work, we computed the set of metabolic fluxes $\mathbf{v}$ which minimize the carbon intake flux $v_{\text{glc}}$ (i.e. maximize the growth yield with respect to the carbon source) under the constraints set by mass-balance, reaction irreversibility, plus a set of three condition-dependent constraints: (i) the acetate production flux $J_{\text{ac}}$, fixed by directly measuring acetate excretion; (ii) the ATP maintenance (ATPM) flux $J_{\text{ATPM}}$, fixed by the observed cellular growth yield; (iii) the biomass sink fluxes $J_\kappa^{\text{BM}}$, set by the cellular biomass composition. The optimization problem reads:

$$\min_{\mathbf{v}} \; v_{\text{glc}} \quad \text{s.t.} \quad S\mathbf{v} = \mathbf{0} \qquad \text{(mass balance constraints)} \tag{N1.1}$$

$$v_i \geq 0 \qquad \text{(if reaction } i \text{ is irreversible)} \tag{N1.2}$$

$$v_{\text{ac}} = J_{\text{ac}} \qquad \text{(acetate excretion, aerobic cond. only)} \tag{N1.3}$$

$$v_{\text{succ}} = J_{\text{succ}} \qquad \text{(succinate excretion, anaerobic cond. only)} \tag{N1.4}$$

$$v_{\text{ATPM}} = J_{\text{ATPM}} \qquad \text{(maintenance energy)} \tag{N1.5}$$

$$v_\kappa^{\text{sink}} = J_\kappa^{\text{BM}} \qquad \text{(for biomass components } \kappa\text{)} \tag{N1.6}$$

The FBA formulation shown above is slightly different than usual. The growth rate $\mu$ is typically an element of the vector $\mathbf{v}$ and corresponds to the rate of a "biomass accumulation" reaction. Instead, the growth rate does not appear explicitly in this formulation, and it is replaced by (condition-dependent) demand fluxes $J_{\text{ac}}$, $J_{\text{succ}}$, $J_{\text{E}}$ and $J_\kappa^{\text{BM}}$. Each of the constraints in Eq. (N1.1)–(N1.6) is described in detail below.

##### 1.2 Mass balance and irreversibility constraints

The basic constraints used in constraint-based modelling are the mass-balance constraints (N1.1), arising from the assumption of stationarity for intracellular metabolites. By setting to zero the time derivative of their concentrations, one obtains a set of linear constraints among the fluxes of the reactions producing or consuming the metabolite. These constraints are written in compact form as a linear system $S\mathbf{v} = \mathbf{0}$, with $S$ being the stoichiometric matrix and $\mathbf{0}$ being the null vector. The stoichiometric matrix encodes the structure of the metabolic network, including exchanges of nutrients iwth the environment. In this work, we used the most recent metabolic network reconstruction *iML1515* [1], which includes 2719 reactions and their association to 1515 genes.

The second class of constraints, Eq. (N1.2), are irreversibility bounds for reactions which are known to be effectively irreversible in physiological conditions[1]. In this work, we use the default mass balance and thermodynamic constraints for the iML1515 metabolic model, with the exception of two reactions involved in the interconversion of acetate and acetyl-CoA. Acetate can be converted into acetyl-CoA via either the acetyl-CoA synthetase (ACS) or via the PTA-ACK pathway. The fluxes through these pathways not only depends on the overall production or consumption of acetate, but also on its external concentration [2]. We set the ACS and PTAr reactions, by default reversible in iML1515, to be irreversible. This avoids the shift to negative values of the PTAr flux at slow growth, where instead the Acs protein is highly expressed [3]. Instead, the PTA-ACK pathway is used at fast growth when acetate is excreted.

##### 1.3 Constraints on exchange fluxes

Overflow metabolism is the prototypical example of apparent "suboptimality" of cellular metabolism, since the growth yield (biomass per carbon consumed) is reduced as a fraction of the carbon flux is excreted in form of acetate. Therefore, optimizing the metabolic fluxes of carbon-limited cells in aerobic conditions predicts vanishing acetate excretion fluxes. We accounted for the overflow metabolism by introducing the constraint (N1.3) which sets the growth-rate dependent acetate excretion flux $v_{\mathrm{ac}}$ to the experimental values modeled as $J_{\mathrm{ac}}$. Similarly, succinate excretion in anaerobic, carbon-limited conditions is not predicted by FBA without additional constraints. This was accounted for by the constraint (N1.4)

[1]These are often called "thermodynamic" constraints. However, this term is misleading as constraint-based models of metabolism can be extended to explicitly include thermodynamic effects (via chemical potentials and reaction free energies). We will thus stick to the term "irreversibility" for this class of constraints.

setting $v_{\text{succ}}$ to the value $J_{\text{succ}}$.

##### 1.4 Energy demand

The so-called ATP maintenance energy flux accounts for the energy (i.e. ATP) consumed by the cell for homeostasis and growth. This flux is traditionally separated in two components: a "non-growth associated maintenance" (NGAM) and a "growth-associated maintenance" (GAM) flux. The first components is modeled as a cellular "ATP maintenance" (ATPM) reaction, describing the hydrolysis of ATP to ADP. Its flux $v_{\text{ATPM}}$ is fixed to a specific value, e.g. 6.86 mmol/$g_{\text{DW}}$h in the default iML1515 model. The second component, GAM, is assumed to be proportional to the growth rate. Given the proportionality to the growth rate, it is usually convenient to encapsulate the GAM in the coefficients of the biomass production reaction. (Since the reaction is mass-balanced, it does not contribute to the biomass accumulation.) Here, we found useful to separate the GAM from the biomass accumulation, and to use the ATPM reaction to account for both GAM and NGAM. This is done in Eq. (N1.5) by setting the ATPM flux $v_{\text{ATPM}}$ to the value $J_{\text{ATPM}} = \sigma_0 + \sigma\mu$, with $\sigma_0$ and $\sigma$ specifying NGAM and GAM, respectively.

We set the parameters of the energy maintenance constraint, $J_{\text{ATPM}}(\mu) = \sigma_0 + \sigma\mu$ by matching the predicted glucose uptake rate $v_{\text{glc}}$ with the experimental data, as described below.

##### 1.5 Definition of the condition-dependent biomass composition

Building blocks (amino acids, nucleotides, and so on) are consumed and converted into biomass (dry weight) as the cells grow. Metabolic models usually have a "biomass" reaction with flux equal to the growth (dilution) rate, $\mu$, which mimics cellular growth: each biomass component $\gamma$ is removed from the system with a flux given by $J_\kappa^{\text{BM}} = s_\kappa^{\text{BM}} \times \mu$. Here, the "stoichiometric" coefficients $s_\kappa^{\text{BM}}$ have units of mmol/$g_{\text{DW}}$, and specify the amount of the metabolite requested form a unit of dry biomass. Since the biomass composition of the cell changes in different growth conditions (see below), the coefficients $s_\kappa^{\text{BM}}$ are also growth rate-dependent. For convenience, we separated the original iML1515 "biomass" reaction into a set of "sink" fluxes $v_\kappa^{\text{sink}}$, one for each biomass component; the sink fluxes are then constrained so as to match the biomass demand $J_\kappa^{\text{BM}}$ (N1.6).

Following Mori et al. [4], we defined a growth-rate dependent biomass composition by adapting the stoichiometry coefficients $s_\kappa^{\text{BM}}$ to each growth conditions by varying the cellular macromolecular composition, while fixing the ratios of individual components: building blocks of the same class (e.g. amino acid, nucleotides, deoxynucleotides) were grouped together, and their coefficients $s_\kappa^{\text{BM}}$ were changed in proportion. The coefficients were then calculated from the macromolecular abundances determined experimentally across conditions, as described below for each macromolecular group.

**1.5.1 Protein, RNA and DNA.** The total abundances of proteins, RNA and DNA per unit of dry mass were obtained using data from Basan et al. [5] (see Supp. Fig. S1A-C). (For anaerobic carbon-limited growth, we used the same biomass composition as for the aerobic case at the same growth rate.) The stoichiometric coefficients for the individual building blocks (amino acids, nucleotides and deoxynucleotides) were determined as follows. For RNA and DNA, we used the reported ratio among nucleotides and deoxynucleotides used in the iML1515 model. For proteins, we made use of ratios of protein abundances obtained from recent mass spectrometry experiments in order to estimate the frequency of the individual amino acids in the expressed proteome, see Supp. Fig. S1K.

It is important to note that the "biomass" reaction also accounts for the polymerization of the building blocks into macromolecules. In particular, a water molecule is produced for each peptidic bond, and a pyrophosphate molecule is produced during the polymerization of either RNA and DNA. Therefore, the sink reactions must take the following form:

$$\begin{aligned} \text{Amino acid sink:} &\quad \text{AA} \longrightarrow \text{H}_2\text{O} \\ \text{Nucleotide sink:} &\quad \text{NTP} \longrightarrow \text{PPi} \\ \text{Deoxynucleotide sink:} &\quad \text{dNTP} \longrightarrow \text{PPi} \end{aligned}$$

Since the experimental biomass composition is expressed as as a fraction of the cellular dry mass, one has to take into account that not all the mass of the building blocks (amino acids, nucleotides and deoxynucleotides, respectively) is incorporated into the biomass, i.e. the mass of water and pyrophosphate (for NTP and dNTP) molecules has to be removed. Failing to account for this can greatly affect the estimated fluxes of the biosynthetic pathways, especially for DNA and RNA given the large mass of pyrophosphate compared to the (deoxy)nucleotide triphosphate.

**1.5.2 Cell wall components.** Given the scarcity of available data, estimating the other cellular components is more complicated and requires additional assumptions.

The mass fraction of lipids for *E. coli* cells in glucose minimal media has been estimated by Gonzalez et al. [6] to be 6%, and by Beck et al. [7] to be 6.7%. In our analysis, we approximated the lipid mass fraction as 6.5% under reference condition (glucose minimal media, growth rate 0.98/h). Furthermore, the mass ratio between the three main cell membrane components, lipid, lipopolysaccharide and peptidoglycan was reported to be 9.1:3.4:2.5 in Neidhardt et al. [8]. We assumed that these ratios are maintained across all conditions. Therefore, the total cell membrane takes roughly 11% of the total dry mass under glucose minimal media condition.

The relative abundances of phospholipids were estimated based on data from Neidhardt et al. [8]: phosphatidylethanolamine (pe) at 76.5%, phosphatidylglycerol (pg) at 18.4%, and cardiolipin (clpn) at 5.1%. We also refered to the data provided in Ref.[6] when classifying the lipids with their numbers of carbon atoms and their levels of saturation. For example, the palmitic lipid (16:0) is reported to have 33.7% of the total cellular lipid mass, we can then have the mass fraction of palmitic phosphatidlyethanolamine ("pe160_p" in the iML1515 model) over the total cellular lipid as $33.7\% \times 76.5\% = 25.8\%$.

**1.5.3 Cellular surface area.** To estimate how the demand of cell membrane precursors varied across different growth conditions, we assumed that the cell membrane material is proportional to the cellular surface. This assumption is equivalent to the proportionality between the mass fraction of cell membrane biomass and the ratio of the cellular surface area and its dry mass (Supp. Fig. S1D).

As initial estimate for the cell surface area, we considered the average cellular widths ($d$) and lengths ($l$) reported in Ref. [5]. For capsule-shaped cells, the volume $V$ and surface $S$ can be computed as:

$$V = \frac{\pi d^2}{4}(l + d/6) \qquad \text{(capsule volume)} \tag{N1.7}$$

$$S = \pi d(l + d/2) \qquad \text{(capsule surface area)} \tag{N1.8}$$

In particular, Eq. (N1.8), divided by the cellular dry mass reported in the same manuscript, provided our first estimate of the desired surface area to dry mass ratio.

To verify our estimation of cell volume and surface area, we calculated the buoyant density with our estimation of cell volume, cell dry mass and assuming that the dry mass accounts for $\sim 30\%$ of the cellular mass [9]. This lead to a buoyant density around $\sim 0.6\text{g/cm}^3$, significantly lower than values $\sim 1.1\text{g/cm}^3$ reported in literature [10]. This suggested that the cell dimensions (cell width and/or length) reported in [5] might be overestimated. We compared the data with the measurements using similar methods in Ref.[11], and found that the cell widths are roughly twice as large in Ref.[5], while the cell lengths are consistent. This variability might be due to different image segmentation protocols, which would affect primarily the cell width (since it is smaller than the length).

Given the observed variability in reported cell width, we reduced the cell width reported in Basan et al. [5] by a constant factor across all conditions, so as to bring the calculated buoyant density up to $1.1g/cm^3$. This new set of cell widths was then used to calculate the surface area according to Eq. (N1.8).

**1.5.4 Other cellular components.** Given the lack of available data, cellular abundances of other cellular component and small molecules were assumed to be constant. In most cases, we used the values reported in the iML1515 model, with a few exceptions for particularly abundant components. Firstly, the mass fraction of glycogen was estimated at 2.5% in Neidhardt et al. [8]. Secondly, the concentrations of glutamate and glutathione were reported in Park et al. [12] to be 96 mM and 16.6 mM, respectively. These concentrations amount to 3.23% and 1.17% of the cellular dry mass, respectively. For the remainder of the small molecules (cofactors, ions), we assumed that their total mass is 1%, with ratios prescribes as in the iML1515 model.

**1.5.5 Final normalization.** Symbolically, the constraint on the total cellular dry mass can be expressed as follows: if $\Delta m_\kappa$ is the specific mass of the $\kappa$-th building block, minus the polymerization byproduct, then:

$$\sum_\kappa \Delta m_\kappa s_\kappa^{\mathrm{BM}} = 1 \text{ g/g}_{\mathrm{DW}} \ . \tag{N1.9}$$

This constraint is crucial, as it determines the overall normalization of the biosynthetic fluxes. When adding up the contribution from all the biomass components, we found that the total mass slightly exceeded 100% (Supp. Fig. S1. Therefore, in each growth conditions, we normalized the sum of the estimated mass fractions of all biomass components so that the constraint (N1.9) was satisfied. The resulting mass fractions were finally used to compute for each condition the demand fluxes for each biomass precursor.

#### 1.6 Flux optimization

The FBA problem described by Eq. (N1.1)-(N1.6) is a linear programming (LP) problem, for which a number of efficient solving algorithms exist. Importantly, this optimization problem has typically multiple solutions, due to the presence of "flux loops", i.e. flux vectors $\boldsymbol{\ell}$ satisfying $S\boldsymbol{\ell} = \mathbf{0}$. We removed this degeneracy my minimizing the $L_2$ norm of the fluxes [13].

Most FBA results refer to either carbon-limited (C-lim) or translationally-inhibited (R-lim) growth condition in MOPS media. Additionally, carbon-limited growth was studied in anaerobic conditions (Supp. Fig. S3). In each of the three growth limitations considered, we defined a growth-dependent biomass composition, and obtained both the glucose intake flux and the acetate excretion fluxes across a range of growth rates, approximately 0.3/h to 1/h. The growth-dependent biomass composition was taken to be the same in the carbon limited growth conditions in presence or absence of oxygen.

For aerobic growth, we first fitted the observed acetate flux as a linear function of the growth rate, and used this linear relation as a constraint in subsequent FBA calculations. Then, we computed the maximum yield solution (minimum glucose uptake) at the growth rates at which the glucose intake rates were obtained. By minimizing the difference between the measured and the modeled glucose fluxes, we determined the parameters $\sigma_0$ and $\sigma$, (NGAM and GAM) determining the ATP maintenance flux across growth rates; the best fit values are reported in Table S2. Using these parameters, we estimated the intracellular fluxes across growth rates by minimizing the glucose intake flux as described in Eq. (N1.1)–(N1.6), followed by the minimization of the $L_2$ norm of the fluxes at fixed (optimal) glucose intake flux. The procedure for anaerobic growth is analogous, except that the constraint Eq. (N1.3) was replaced by the constraint on succinate excretion, Eq. (N1.4).

### Supplementary Note 2: Flux and functional decompositions of *E. coli*'s metabolic network

The main focus of this work is on the metabolism of bacterial cells, studied from the point of view of genome-scale models of metabolism. Typically, such models include nutrient uptake, catabolism and anabolism, but exclude the details of the synthesis of macromolecules (e.g. proteins and RNA). In growing cells, the network transforms nutrients into biomass building blocks (e.g. individual amino acids and NTPs), and provide the energy (hydrolizable ATP) to sustain a variety of cellular activities, including the polymerization of the building blocks into macromolecules. Each reaction contributes to varying extent to the production of biomolecules and energy. Furthermore, since the cellular biomass composition and the energetic demands depend strongly on the state of the cell, the activity of metabolic reactions has to be regulated depending on the environment.

In this work we aimed to define the natural definitions of biological *functions* for the metabolic network of *E. coli*, and to define quantitatively the contribution of individual metabolic reactions to each metabolic function. In biology, the word "function" is usually understood as a description of the activity of some biological component (from cell types in multicellular organisms, to organelles, to individual genes) that emerges as a result of adaptation, or confers some fitness advantage [14, 15]. As we circumscribe ourselves to the study of the metabolic reactions, it is natural to consider the production of biomass precursors and of energy to be the set of functions that can be attributed to each reaction: each reaction directly or indirectly contributes to the production of precursors and energy, whose production is obviously useful for cellular fitness. A quantitative, network-level definition of metabolic functions can potentially allow to connect the activity of individual reactions or enzymes on one hand, and physiology and the environment on the other, thus providing insights on regulation of gene expression at the genome-scale level. For example, reactions with different functions are likely to be differently regulated in response to intracellular or environmental cues, while reactions with similar functions are instead more likely to be co-regulated. This is reflected, for example, in the shared operon structures of many genes encoding for different steps of the same biosynthetic pathways.

#### 2.1 Flux decomposition

The functional decomposition of the *E. coli* metabolic network described in this work is based on a decomposition of metabolic fluxes whose mathematical basis have been described previously [16], and applied to the study of energy and biomass production within CAFBA solutions [4]. Here, we considerably extend its range of application, allowing to study realistic FBA solutions with growth-dependent biomass coefficients and experimental constraints (such as acetate overflow). We use the flux decomposition prescribed by the optimization problem at hand to define a consistent *functional* decomposition, and we discuss in detail how external constraints (e.g. enforcing a givens acetate production), and negative flux components can be accounted for in this framework.

Given a constraint-based model of metabolism, the flux decomposition can only be applied to an *optimal* flux solution satisfying the mass-balance, thermodynamic and, possibly, additional constraints on the fluxes. While this might seem a strong limitation, the maximum yield solution provides generally good estimates of the intracellular fluxes, especially for the central carbon pathways. In aerobic conditions, overflow metabolism can be captured by adding a growth rate-dependent constraint, leading to good agreement with experimental $^{13}$C fluxes [17].

The optimal solution $\mathbf{v}$ to the optimization problem (N1.1)-(N1.6) depends implicitly on the values of the constrained demand fluxes $J_\kappa^{\mathrm{BM}}$, $J_{\mathrm{ATPM}}$ and $J_{\mathrm{ac}}/J_{\mathrm{succ}}$ (depending on the availability of oxygen). We now want to express $\mathbf{v}$ as a function of these constraints. Because of dimensional analysis and the linearity of the system, it is possible to write the solution as a linear combination of different components:

$$\mathbf{v} = \sum_\kappa \boldsymbol{\beta}^{(\kappa)} J_\kappa^{\mathrm{BM}} + \boldsymbol{\eta} J_{\mathrm{ATPM}} + \boldsymbol{\alpha} J_{\mathrm{ac}} \;, \tag{N2.1}$$

where the sum runs over all biomass building blocks $\kappa$, and $\boldsymbol{\beta}^{(\kappa)}$, $\boldsymbol{\eta}$ and $\boldsymbol{\alpha}$ are constant vectors, called *flux modes*. In the case of anaerobic growth, Eq. (N2.1) is replaced by the analogous expression

$$\mathbf{v} = \sum_\kappa \boldsymbol{\beta}^{(\kappa)} J_\kappa^{\mathrm{BM}} + \boldsymbol{\eta} J_{\mathrm{ATPM}} + \boldsymbol{\alpha} J_{\mathrm{succ}} \;. \tag{N2.2}$$

The relations Eq. (N2.1) and (N2.2) can both be written in a more compact form:

$$\mathbf{v} = \sum_\gamma \boldsymbol{\xi}^{(\gamma)} J_\gamma \;, \tag{N2.3}$$

where here $\gamma$ labels any of the terms appearing in Eq. (N2.1) or (N2.2). These expressions describe a *parameterization* of the optimal flux $\mathbf{v}$ as a function of the constraints, $\mathbf{v} = \mathbf{v}(\{J_\gamma\})$. The parameterization is only valid locally: the global solution to the optimization problem is a complicated piecewise linear function of the constraints $J_\gamma$, with the flux modes $\boldsymbol{\xi}^{(\gamma)}$ "jumping" between different vectors as the demand fluxes $J_\gamma$ are varied.

##### 2.1.1 Computation of the coefficients of the decomposition.

The vectors $\boldsymbol{\xi}^{(\gamma)}$ in Eq. (N2.3) are effectively the partial derivative of the optimal flux vector with respect to the constraints:

$$\xi_i^{(\gamma)} = \frac{\partial v_i}{\partial J_\gamma} \tag{N2.4}$$

This suggests that the flux modes $\boldsymbol{\xi}^{(\gamma)}$ can be computed numerically by perturbing the optimal solution by slightly changing the demand fluxes. Consider a network with $R$ reactions and $N$ demand fluxes $J_\gamma$. We determine $R \times N$ matrix elements $\xi_i^{(\gamma)}$ by re-computing the optimal flux $\mathbf{v}$ using a slightly different demand flux $J_\gamma \to J_\gamma(1+\epsilon)$, leading to a set of perturbed fluxes $\mathbf{v} \to \mathbf{v} + \delta\mathbf{v}^{(\gamma)}$. In practice, we found that a value $\epsilon = 10^{-4}$ allowed for accurate calculations without generating artifacts due to the finite numerical precision of the system. Then, for each reaction $i$, we solve the system:

$$\begin{pmatrix} J_1 & J_2 & \cdots & J_{N-1} & J_N \\ J_1 \cdot (1+\epsilon) & J_2 & \cdots & J_{N-1} & J_N \\ J_1 & J_2 \cdot (1+\epsilon) & \cdots & J_{N-1} & J_N \\ \vdots & \vdots & \ddots & \vdots & \vdots \\ J_1 & J_2 & \cdots & J_{N-1} \cdot (1+\epsilon) & J_N \end{pmatrix} \begin{pmatrix} \xi_i^{(1)} \\ \xi_i^{(2)} \\ \vdots \\ \xi_i^{(N-1)} \\ \xi_i^{(N)} \end{pmatrix} = \begin{pmatrix} v_i \\ v_i + \delta v_i^{(1)} \\ v_i + \delta v_i^{(2)} \\ \vdots \\ v_i + \delta v_i^{(N-1)} \end{pmatrix} \tag{N2.5}$$

Note that the original flux vector $\mathbf{v}$ can be used as a constraint, so only $N-1$ perturbations are necessary to solve for the coefficients $\xi_i^{(\gamma)}$; for instance, this allows not to take the derivative against the "other" biomass components. Optionally, taking the last derivative against $J_N$ allows to check whether the flux decomposition satisfies Eq. (N2.3) is satisfied up to numerical precision.

Since the flux decomposition is defined in terms of derivatives with respect to the constraints, it is clear that the analysis presented is only valid if the demand fluxes are *independent* of each other. For example, it is possible that, in a given network, the production of two metabolites A and B is constrained to happen in a fixed ratio. In this case, the derivatives $\partial\mathbf{v}/\partial J_\mathrm{A}$ and $\partial\mathbf{v}/\partial J_\mathrm{B}$ are not defined, since it is not possible to perturb the two demand fluxes separately.

Incidentally, note that without any flux norm minimization, or even with the minimization of the $L_1$ norm, many irreversibility constraints are marginal, and the value of many fluxes is assigned randomly during the optimization due to the presence of loops. This variability leads to numerical problems when computing the perturbations. Instead, using the $L_2$ norm minimization does not lead to such degeneracies. Furthermore, using this norm tends to increase the number of active reactions in the cell compared to the $L_1$ norm, allowing us to perform the flux decomposition on more reactions; this property is desirable in the light of the functional decomposition of the proteins described in Supp. Note 5, since it maximizes the amount of proteins whose metabolic function can be analyzed with this method.

**2.1.2 Coarse-graining the functional decomposition.** An important property of the flux decomposition is that a simplified version of Eq. (N2.1) can be obtained if some demand fluxes are proportional to each other. This happens, for instance, when biomass components are produced in fixed stoichiometric ratios.

Consider for example two biomass components $\alpha$ and $\beta$, and their corresponding flux modes $\boldsymbol{\xi}^{(\alpha)}$ and $\boldsymbol{\xi}^{(\beta)}$, as well as the demand fluxes $J_\alpha$ and $J_\beta$. If these two components are present in fixed proportion in the cellular biomass across conditions, then the demand fluxes are proportional to each other, e.g. $J_\beta = qJ_\alpha$ for some $q > 0$. In this case, it is possible to define a single flux mode that accounts for the production of both biomass precursors. This can be done in multiple ways depending on the normalization of the flux components, e.g.:

$$\boldsymbol{\xi}^{(\alpha)} J_\alpha + \boldsymbol{\xi}^{(\beta)} J_\beta = [\boldsymbol{\xi}^{(\alpha)} + q\boldsymbol{\xi}^{(\beta)}] J_\alpha \tag{N2.6}$$

$$= [\boldsymbol{\xi}^{(\alpha)}/q + \boldsymbol{\xi}^{(\beta)}] J_\beta \tag{N2.7}$$

$$= \left[\frac{\boldsymbol{\xi}^{(\alpha)} + q\boldsymbol{\xi}^{(\beta)}}{1+q}\right] (J_\alpha + J_\beta) \ . \tag{N2.8}$$

In all cases, the term in the square brackets represents a flux mode associated to the synthesis of the two components $\alpha$ and $\beta$. This property is useful if one is not interested in the details of how fluxes are associated to the production of individual precursors, but only in a coarser description of the network. This also allows to reduce the number of optimizations needed to compute the flux modes. (The linear properties of the flux decomposition will also be crucial in the application of the flux decomposition to realistic networks, as shown in Supp. Note 2.2.3.)

In particular, if the biomass composition of the metabolic model is considered to be fixed across conditions, all biomass-associated demand fluxes are proportional to each other, and they can be collected into a single flux mode associated to biomass production. The growth rate itself can be conveniently used as "demand flux". Taking the demand fluxes to be $J_\kappa = s_\kappa^{\mathrm{BM}} \mu$, one obtains:

$$\sum_\kappa \boldsymbol{\xi}^{(\kappa)} J_\kappa = \left( \sum_\kappa s_\kappa^{\mathrm{BM}} \boldsymbol{\xi}^{(\kappa)} \right) \mu \equiv \boldsymbol{\xi}^{(BM)} \mu \ , \tag{N2.9}$$

with the flux mode $\boldsymbol{\xi}^{(BM)}$ describing the accumulation of biomass in the cell. In the extreme case in which all demand fluxes are proportional to the growth rate (e.g. neglecting the growth-independent ATP maintenance flux $\sigma_0$, which is often a reasonable approximation for fast growing cells), then the cellular fluxes $\mathbf{v}$ can be locally parametrized using a single growth-associated flux mode $\boldsymbol{\xi}$:

$$\mathbf{v} = \boldsymbol{\xi} \mu \ . \tag{N2.10}$$

Here, $\boldsymbol{\xi}$ summarizes the "metabolic state" of the cell [16], since it captures the overall metabolic patterns and growth yields associated to the network.

#### 2.2 Functional decomposition

The flux decomposition can be used to define a *functional* decomposition of the reaction fluxes (and later of the proteome, as described in Supp. Note 4). Specifically, our aim is to define a set of metabolic functions, and to quantify how much does each reaction contribute to each function.

**2.2.1 Naïve functional decomposition.** We first rewrite the flux decomposition (N2.3) in terms of *flux components* $\mathbf{v}^{(\gamma)}$ as:

$$\mathbf{v} = \sum_\gamma \mathbf{v}^{(\gamma)} \quad \text{where} \quad \mathbf{v}^{(\gamma)} = \boldsymbol{\xi}^{(\gamma)} J_\gamma \tag{N2.11}$$

In this form, the flux $\mathbf{v}$ is expressed as the sum of several components $\mathbf{v}^{(\gamma)}$, where each components is associated to a single demand flux $J_\gamma$. When interpreting the demand fluxes as *metabolic functions*, Eq. (N2.3) intuitively provides as the basis of a functional decomposition.

As a simplified, but concrete example, consider the case of cells growing aerobically on minimal carbon medium, with a fixed (condition-independent) biomass composition and without fermentative metabolism (obtained e.g. by growing cells in a chemostat with a low dilution rate). In this case, it is possible to express the optimal solution as the linear combination of two components, one associated to energy production $J_E$ and one associated to biomass production $J_B$:

$$\mathbf{v} = \boldsymbol{\xi}^{(E)} J_E + \boldsymbol{\xi}^{(B)} J_B = \mathbf{v}^{(E)} + \mathbf{v}^{(B)} \tag{N2.12}$$

This expression partitions the flux of each reaction in two terms, one associated to energy production, and the other to biosynthesis. For each reaction $i$, $F_i^{\mathrm{E}} \equiv v_i^{\mathrm{E}}/v_i$ represents the fraction of flux associated to energy, and the remainder $F_i^{\mathrm{B}} = 1 - F_i^{\mathrm{E}} = v_i^{\mathrm{B}}/v$ the fraction associated to the production of biomass components. These two quantities, $F_i^{\mathrm{E}}$ and $F_i^{\mathrm{B}}$, describe the contribution of the $i$-th reaction to energy and biomass generation, and therefore describe the *function* of the reaction.

More in general, a possible definition for $F_i^{\gamma}$ is the ratio between the flux component $v_i^{(\gamma)}$ and the overall flux $v_i$:

$$F_i^{\gamma} = \frac{v_i^{(\gamma)}}{v_i} = \frac{v_i^{(\gamma)}}{\sum_{\alpha} v_i^{(\alpha)}} \tag{N2.13}$$

Clearly, this definition can only be applied to reactions with non-zero fluxes. If a reaction flux is zero, then all the flux derivatives are also zero, and the decomposition Eq. (N2.13) cannot be applied. Even focusing on flux-carrying reactions only, two main issues prevent the straightforward application of Eq. (N2.13) in the concrete case of genome-scale models. These are briefly described in the Main Text; here we provide and extended discussion.

- **Sign-mismatch.** If some flux components have opposite sign, then at least one of the fractions $F_i^{\gamma}$ will be negative. In this case, the interpretation of the flux decomposition is not straightforward. What is the meaning of a *negative* contribution to a biological function? However, note that if *all* flux components are negative, then all fractions $F_i^{\gamma}$ are positive. The conventional direction of the reaction does not matter, but the relative sign among the flux components does.

- **Interpretation of constraints.** The demand fluxes $J_{\gamma}$ cannot always be straightforwardly interpreted as metabolic function. In particular, the interpretation of the additional terms arising in Eq. (N2.1) due to the presence of phenomenological constraints must emerge from biological considerations, and have to be treated on a case-by-case basis. For instance, the production of signalling molecules, antibiotics, extracellular polymeric substances from cells in biofilms, might all be considered as proper metabolic functions; but the meaning of the functional decomposition in presence of the constraint on acetate excretion is less clear. We discuss this latter case in Section 2.3.

**2.2.2 Interpretation of negative components.** To describe the problem posed by the negative flux components, it is worth discussing in detail a simple example, analogous to that shown in Main Text Figure 1D. The system shown in Fig. N2.1a includes three reactions, with fluxes $v_1$, $v_2$ and $v_3$. These

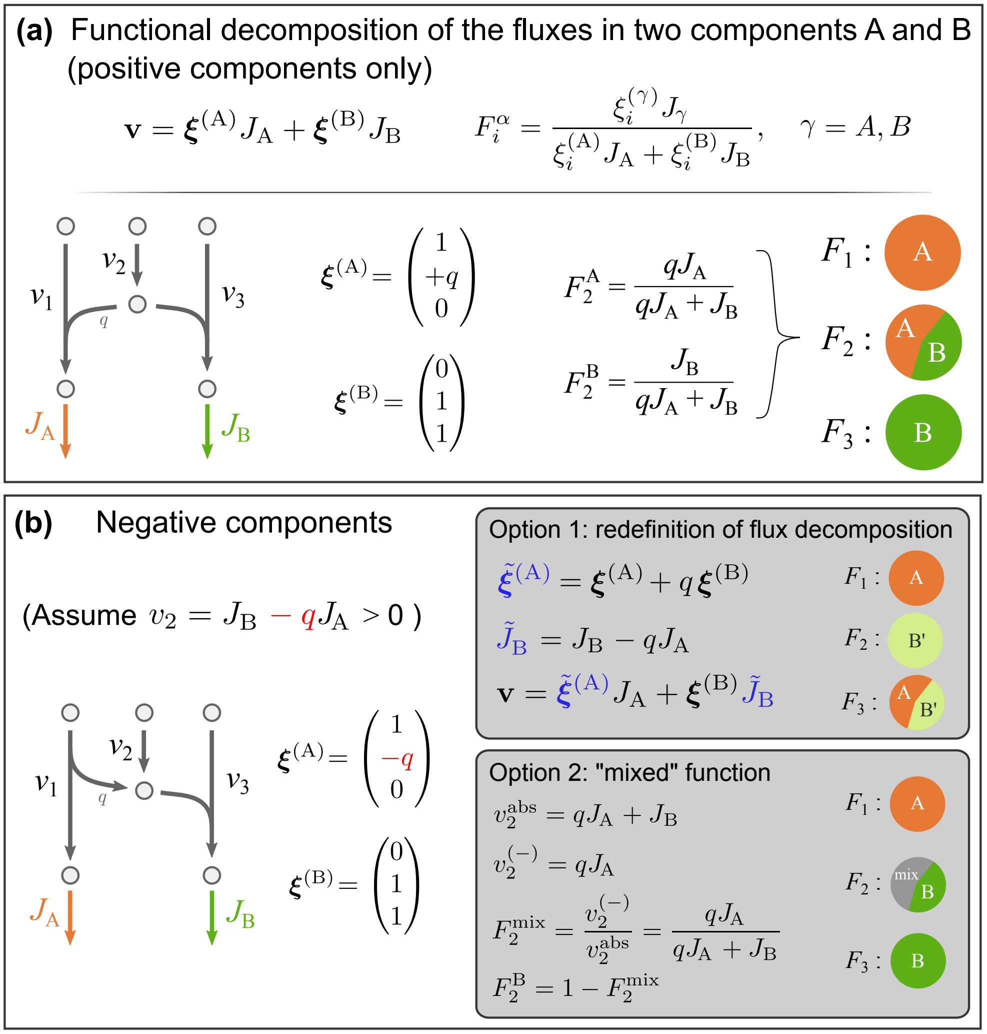

**Figure N2.1.** **(a)** Functional decomposition when all flux components have the same sign. **(b)** Functional decomposition when some flux components have mixed signs. In this example, the flux $v_2$ is written as the difference of $J_B$ and $qJ_A$. A functional decomposition can be performed consistently in two ways: (1) Redefining the decomposition by taking linear combinations of the metabolic modes $\boldsymbol{\xi}^{(\gamma)}$ and the demand fluxes $J_\gamma$ in order to remove the negative entries; (2) Absorbing the negative flux components into a "mixed" functional component associated to neither A or B. See text for details.

reactions convert three metabolites (X, Y and Z) into two other metabolites A and B, which are drained by the two demand fluxes $J_A$ and $J_B$; the $v_2$ flux supplies an intermediate metabolite C that is requested by both reaction 1 (with a stoichiometric coefficient $q$) and 3:

$$\text{Reaction 1}: \quad X + qC \to A \tag{N2.14}$$

$$\text{Reaction 2}: \quad Y + C \to B \tag{N2.15}$$

$$\text{Reaction 3}: \quad Z \to C \tag{N2.16}$$

The steady state equations are simple enough to be solved analytically. We find that the flux vector can be written as

$$\mathbf{v} = \boldsymbol{\xi}^{(\mathrm{A})} J_{\mathrm{A}} + \boldsymbol{\xi}^{(\mathrm{B})} J_{\mathrm{B}} \tag{N2.17}$$

where the components of the two vectors $\boldsymbol{\xi}^{(\mathrm{A})} = (1\ q\ 0)^{\mathrm{T}}$ and $\boldsymbol{\xi}^{(\mathrm{B})} = (0\ 1\ 1)^{\mathrm{T}}$ are non-negative. Since $v_1 = J_{\mathrm{A}}$, reaction 1 contributes completely to function A; similarly, reaction 3 contributes only to function B. The flux $v_2$ is split in two components, $v_2 = qJ_{\mathrm{A}} + J_{\mathrm{B}}$. Therefore, we assign a fraction $F_1^A = qJ_A/(qJ_A + J_B)$ to function A, and the rest to function B.

Let us now consider the case shown in Fig. N2.1b. The reaction network is similar to the previous case, except now the intermediate metabolite is also produced by reaction 1 (with a stoichiometric coefficient $q$):

Reaction 1 : $$\mathrm{X} \to \mathrm{A} + q\mathrm{C} \tag{N2.18}$$

Reaction 2 : $$\mathrm{Y} + \mathrm{C} \to \mathrm{B} \tag{N2.19}$$

Reaction 3 : $$\mathrm{Z} \to \mathrm{C} \tag{N2.20}$$

The flux decomposition is the same as in the previous case, except for the substitution $q \to -q$. This leads to $\boldsymbol{\xi}^{(\mathrm{A})} = (1\ \ -q\ \ 0)^{\mathrm{T}}$, $\boldsymbol{\xi}^{(\mathrm{B})} = (0\ 1\ 1)^{\mathrm{T}}$ and, in particular, $v_2 = J_B - qJ_A$. In the following, we assume for simplicity $v_2 > 0$, i.e. $J_B > qJ_A$. In this situation, reaction 2 compensates for the mismatch between the production of metabolite C from reaction 1 and its the demand from reaction 3. A naïve application of Eq. (N2.13) does not lead to a sensible functional decomposition, as we would obtain a negative component $F_2^{\mathrm{A}} < 0$. There are at least two ways to address this issue, as discussed below.

**2.2.3 Coupling demand fluxes - example.** The first option relies on the fact that the flux decomposition (N2.17) is not unique: it is possible to consider linear combinations of the flux modes $\boldsymbol{\xi}^{(\gamma)}$ and demand fluxes $J_\gamma$. Setting aside for the moment the biological interpretation of the decomposition, we seek a redefinition of the vector $\boldsymbol{\xi}^{(\mathrm{A})}$ so that its components are non-negatives. This can be done by adding and subtracting a term proportional to $\boldsymbol{\xi}^{(B)}$ in the flux decomposition:

$$
\begin{aligned}
\mathbf{v} &= \boldsymbol{\xi}^{(\mathrm{A})} J_{\mathrm{A}} + \boldsymbol{\xi}^{(\mathrm{B})} J_{\mathrm{B}} && \text{(N2.21)}\\
&= \boldsymbol{\xi}^{(\mathrm{A})} J_{\mathrm{A}} + \boldsymbol{\xi}^{(\mathrm{B})} J_{\mathrm{B}} + \left[\boldsymbol{\xi}^{(\mathrm{B})} \cdot (q J_{\mathrm{A}}) - \boldsymbol{\xi}^{(\mathrm{B})} \cdot (q J_{\mathrm{A}})\right] && \text{(N2.22)}\\
&= \left[\boldsymbol{\xi}^{(\mathrm{A})} + q\boldsymbol{\xi}^{(\mathrm{B})}\right] J_{\mathrm{A}} + \boldsymbol{\xi}^{(\mathrm{B})} \left[J_{\mathrm{B}} - q J_{\mathrm{A}}\right] && \text{(N2.23)}\\
&= \tilde{\boldsymbol{\xi}}^{(\mathrm{A})} J_{\mathrm{A}} + \boldsymbol{\xi}^{(\mathrm{B})} \tilde{J}_{\mathrm{B}} && \text{(N2.24)}
\end{aligned}
$$

We see that the new flux mode $\tilde{\boldsymbol{\xi}}^{(\mathrm{A})} = \boldsymbol{\xi}^{(\mathrm{A})} + q\boldsymbol{\xi}^{(\mathrm{B})} = \begin{pmatrix} 1 & 0 & q \end{pmatrix}^{\mathrm{T}}$ no longer has negative entries. However, the redefinition of $\tilde{\boldsymbol{\xi}}^{(\mathrm{A})}$ goes hand in hand with a redefinition of the demand flux $J_{\mathrm{B}}$ as $\tilde{J}_{\mathrm{B}} = J_{\mathrm{B}} - q \cdot J_{\mathrm{A}}$. Now that all flux components are positive, we can apply Eq. (N2.13) to obtain the functional decomposition of the reaction network. First, $v_1 = J_{\mathrm{A}}$, so its function cannot be anything other than the synthesis of metabolite A. The flux of the second reaction equals the redefined demand flux, $v_2 = \tilde{J}_{\mathrm{B}}$, and therefore the reaction is completely dedicated to the synthesis of metabolite B. Finally, $v_3 = q \cdot J_{\mathrm{A}} + \tilde{J}_{\mathrm{B}}$, hence reaction 3 contributes both to the production of metabolite B (directly) and of metabolite A (through removal of the byproduct C).

Biologically, the flux $q \cdot J_{\mathrm{A}}$ represent the amount of metabolite B synthesized as a byproduct of the synthesis of metabolite A. Once the latter fraction is subtracted, $\tilde{J}_{\mathrm{B}}$ represents the amount of metabolite B which is synthesized by the cell without considering the activity of other pathways. We have thus found that in order to "fix" the negative entries, we had to *couple* the metabolic functions (and corresponding demand fluxes) so that we could *decouple* the flux modes. A generic linear combination of the two demand fluxes $J_{\mathrm{A}}$ and $J_{\mathrm{B}}$ does not have any intrinsic biological meaning; however, it should not be taken for granted that the initial choice makes sense, either. For instance, when two metabolites are co-produced by the same pathway, it can be more reasonable to define the simultaneous production of both metabolites as a unique metabolic function.

**2.2.4 Introduction of mixed functions.** Instead of modifying the definition of the demand fluxes and of the flux modes, an alternative option is to keep the same flux decomposition $v_i^{(\gamma)}$, but to change the definition of the functional decomposition $F_i^{\gamma}$, Eq. (N2.13), so as to account for the negative flux components. In our example, this would change the functional decomposition of reaction 2, without changing the decomposition of reactions 1 and 3.

As we discussed earlier, the flux through reaction 2 is $v_2 = J_{\mathrm{B}} - q \cdot J_{\mathrm{A}}$. Consider the case in which

$v_2$ is small compared to the absolute value of the two components. In this case, the flux through the reaction is strongly affected by any variation in the fluxes $J_\mathrm{A}$ and $J_\mathrm{B}$, and is adjusted so as to keep the concentration of metabolite C constant. This is the role played by anaplerotic reactions, which allow to keep a constant concentration of intermediate metabolites within pathways. In a sense, the metabolic function of $v_2$ is associated to the synthesis of neither A or B, but it is required to keep the concentration of the intermediate metabolite. This peculiar role is not captured by any biological function associated with a demand flux; instead, a new biological function has to be introduced. We therefore aimed to extend Eq. (N2.13) by defining a "mixed function", whose share $F^\mathrm{mix}$ measures the extent by which a reaction balances internal metabolites, as opposed to matching the demand fluxes. In our example, the net flux $v_2 = J_B - qJ_A$ is less than the sum of the absolute values of each flux component, $v_2^\mathrm{abs} = J_B + qJ_A$. We can then assign the fraction of "useful" flux to function B, and the rest to a "mixed" function:

$$F_2^B = \frac{|v_2|}{\sum_\gamma |v_2^{(\gamma)}|} = \frac{J_B - qJ_A}{J_B + qJ_A} \, , \tag{N2.25}$$

$$F_2^\mathrm{mix} = 1 - F_2^B = \frac{2qJ_A}{J_B + qJ_A} \, . \tag{N2.26}$$

We will show in a following section how this example can be generalized to multiple flux components.

**2.2.5 General case.** In this section, we generalize the lessons illustrated in the previous example, Fig. N2.1, and provide a general framework that can be applied to genome-wide models of metabolism.

**Coupling demand fluxes.** The definition of the flux decomposition, Eq. (N2.1), shows that the flux vectors $\mathbf{v}$ can be *locally* considered as elements of a vector space, with the flux modes $\boldsymbol{\xi}^{(\gamma)}$ providing a basis, and the demand fluxes being the coordinates of the flux with respect to this basis. However, infinite choices exist for such coordinate system, and each choice leads to a different set of biological functions used in the functional decomposition.

The most general linear transformation that can be applied to the demand fluxes is described by an invertible "coupling" matrix $C$, combining the demand fluxes $J$ into "coupled" fluxes $\tilde{J}_\delta = \sum_\gamma C_{\gamma\delta} J_\delta$.

The flux modes $\boldsymbol{\xi}$ will be transformed into new components $\tilde{\boldsymbol{\xi}}$ with the opposite transformation $C^{-1}$:

$$\mathbf{v} = \sum_{\gamma} \boldsymbol{\xi}^{(\gamma)} J_{\gamma} \tag{N2.27}$$

$$= \sum_{\gamma,\delta,\lambda} \boldsymbol{\xi}^{(\gamma)} C^{-1}_{\gamma\delta} C_{\delta\lambda} J_{\lambda} \tag{N2.28}$$

$$= \sum_{\delta} \left[ \sum_{\gamma} \boldsymbol{\xi}^{(\gamma)} C^{-1}_{\gamma\delta} \right] \left[ \sum_{\lambda} C_{\delta\lambda} J_{\lambda} \right] \tag{N2.29}$$

$$\equiv \sum_{\delta} \tilde{\boldsymbol{\xi}}^{(\delta)} \tilde{J}_{\delta} \ . \tag{N2.30}$$

If the matrix $C$ is the identity matrix, then the flux decomposition is unchanged. Diagonal terms different from 1 change the overall scale of the demand fluxes, while off-diagonal entries combine different demand fluxes. If the matrix $C$ is chosen carefully, then the new flux modes can be substantially "decoupled", i.e. the new flux components $\tilde{\mathbf{v}}^{(\gamma)} = \tilde{\boldsymbol{\xi}}^{(\gamma)} \tilde{J}_{\gamma}$ are much less affected by the sign-mismatch problem. In fact, we argue that, generally, *the most biologically meaningful choice for a set* $\{\gamma\}$ *of metabolic functions is the one that minimizes the negative entries in the flux decomposition* $F_i^{\gamma}$. This guiding principle is extremely useful in the application of the flux and functional decompositions to generic genome-scale model of metabolism. It should also be possible to take this principle literally, and numerically optimize the off-diagonal entries of the coupling matrix $C$ so that the cases of sign-mismatch are minimized. In this work, however, we use a more conservative approach in the analysis of *E. coli* metabolism, which is described in Supplementary Note 2.3.

**Mixed functions.** In order to define the functional share $F_i^{\mathrm{mix}}$ of the mixed function, we note that $F_i^{\mathrm{mix}}$ should be close to 1 in presence of large flux components with almost exact cancellations, and equal to zero if all flux components have the same sign. A definition of $F_i^{\mathrm{mix}}$ that satisfies these requirements is the following:

$$F_i^{\mathrm{mix}} = 1 - \frac{|v_i|}{\sum_{\gamma} \left| v_i^{(\gamma)} \right|} \ . \tag{N2.31}$$

This quantity is also useful to quantify the presence of sign-mismatch in the flux components, and to assess the performance of different sets of biological functions associated to the flux decomposition. This is done in Fig. S2A-F, where we show the reduction in negative flux components associated to coupling different metabolic functions.

The remaining functional share, $1 - F_i^{\mathrm{mix}}$, has to be allocated to the biological functions associated with the positive flux components. To do so, we follow the procedure below. (We suppress here the reaction index $i$ to reduce clutter.)

- If the flux $v$ is negative, we first change its sign, along with the sign of all flux components.
- We compute $F^{\text{mix}}$ using (N2.31).
- We define new auxiliary flux components $\mathbf{w}$ with the requirement that $\sum_\gamma w^\gamma = v$ and $w^\gamma = 0$ if $v^\gamma \leq 0$. We do so by iteratively distributing the sum of the negative components $v^{(-)}$ on the nonzero positive components as follows:
  1. Start with $w^\gamma = v^{(\gamma)}$ if $v^\gamma > 0$, or $w^\gamma = 0$ otherwise.
  2. Compute the sum of the negative components, $v^{(-)} = \sum_{\gamma:\, v^{(\gamma)}<0} v^{(\gamma)}$.
  3. Call $N$ the number of strictly positive components $w^\gamma$ and $w^{\text{min}} = \min_{\gamma: w^\gamma > 0} w^\gamma$ their minimum value.
  4. If $v^{(-)} > N \cdot w^{\text{min}}$, remove $w^{\text{min}}$ from all positive entries of $\mathbf{w}$, reduce $v^{(-)}$ by $N \cdot w^{\text{min}}$, and go back to the previous step.
  5. If instead $v^{(-)} \leq N \cdot w^{\text{min}}$, remove $v^{(-)}/N$ from all positive entries of $\mathbf{w}$, and exit.
- The functional decomposition is then computed as:

$$F^\gamma = (1 - F^{\text{mix}}) \cdot \frac{w^\gamma}{\sum_\gamma w^\gamma} = \frac{w^\gamma}{\sum_\gamma |v^\gamma|} \tag{N2.32}$$

Note that all fractions sum up to one: $F^{\text{mix}} + \sum_\gamma F^\gamma = 1$.

##### 2.3 Application to *E. coli* metabolism in aerobic conditions

In this section we apply the previous definitions and results to define a consistent functional decomposition of *E. coli* reaction network. Given that the cases of aerobic and anaerobic growth differ mainly in how the exchange fluxes are constrained (with constraints on either acetate or succinate excretion), we will describe in detail the functional decomposition for the aerobic case only. In this case, the flux decomposition is given by Eq. (N2.1), which we report here for convenience of the reader:

$$\mathbf{v} = \boldsymbol{\eta} J_{\text{ATPM}} + \boldsymbol{\alpha} J_{\text{ac}} + \sum_\kappa \boldsymbol{\beta}^{(\kappa)} J_\kappa^{\text{BM}} \,.$$

We will redefine the demand fluxes in two steps; first, combining combining the maintenance energy ($\boldsymbol{\eta} J_{\mathrm{ATPM}}$) and acetate components ($\boldsymbol{\alpha} J_{\mathrm{ac}}$), leading to the definition of "respiration" and "fermentation" fluxes; second, by combining the energy and biomass ($\boldsymbol{\beta}^{(\kappa)} J_{\kappa}^{\mathrm{BM}}$) terms in order to properly account for the energy produced during biosynthetic activities.

**2.3.1 Acetate production.** The flux $J_{\mathrm{ac}}$ used to constraint the acetate exchange flux represents an additional unit-bearing quantity that can be used to describe the flux vector $\mathbf{v}$, leading to an extra term $\boldsymbol{\alpha} J_{\mathrm{ac}}$ in Eq. (N2.1). While the vector $\boldsymbol{\alpha}$ can be computed by taking perturbing the optimal solution by varying $J_{\mathrm{ac}}$, the interpretation of this term is not straightforward. Should the production of acetate be considered a "biological function" on par with "energy production" or "biomass production"?

To answer this question, we first note that acetate production in *E. coli* is eminently associated to aerobic fermentation, i.e. the conversion of glucose into acetate and carbon dioxide:

$$\mathrm{glc} \rightarrow 2 \cdot \mathrm{ac} + 2 \cdot \mathrm{CO_2} + \mathrm{energy}\ . \tag{N2.33}$$

However, the flux component $\mathbf{v}^{(\mathrm{ac})} = \boldsymbol{\alpha} J_{\mathrm{ac}}$ is mass balanced, implying that it cannot lead to any net production of energy (conversion of ADP into ATP). In fact, the vector $\boldsymbol{\alpha}$ transforms glucose into acetate and CO2, without a net production of energy; furthermore, the stoichiometric ratios between $\alpha_{ac}$, $\alpha_{glc}$ and $\alpha_{co_2}$ are not the ones expected from Eq. N2.33.

Second, we note that the entries of the vector $\boldsymbol{\eta}$ associated to energy (ATP) production correspond to the well known aerobic respiration pathways (glycolysis, TCA cycle, oxidative phosphorylation), both below and above the growth rate $\mu_{\mathrm{ac}}$ at which *E. coli* cells start excreting acetate. On the other hand, the entries of $\boldsymbol{\alpha}$ corresponding to the TCA cycle are negative, suggesting that $\boldsymbol{\alpha}$ represents the difference between a component representing aerobic fermentation and a term proportional to aerobic respiration, proportional to $\boldsymbol{\eta}$ (also written as $\boldsymbol{\eta}^{(\mathrm{r})}$ to indicate aerobic respiration). To disentangle the two components we proceed to remove the negative entries from $\boldsymbol{\alpha}$ by adding and subtracting a term proportional to $\boldsymbol{\eta} \cdot J_{\mathrm{ac}}$ to the flux decomposition, as described in Section 2.2.3. Focusing on the two terms proportional to the maintenance flux $J_{\mathrm{ATPM}}$ and acetate flux $J_{\mathrm{ac}}$, we can combine the fluxes as follows:

$$
\begin{aligned}
\mathbf{v} &= \boldsymbol{\eta} J_{\mathrm{ATPM}} + \boldsymbol{\alpha} J_{\mathrm{ac}} + (\dots) \\
&= \boldsymbol{\eta} J_{\mathrm{ATPM}} + \boldsymbol{\alpha} J_{\mathrm{ac}} + \boldsymbol{\eta} \cdot (y_{\mathrm{ac}}^{\mathrm{E}} J_{\mathrm{ac}}) - \boldsymbol{\eta} \cdot (y_{\mathrm{ac}}^{\mathrm{E}} J_{\mathrm{ac}}) + (\dots) \\
&= \boldsymbol{\eta} \left[ J_{\mathrm{ATPM}} - y_{\mathrm{ac}}^{\mathrm{E}} \cdot J_{\mathrm{ac}} \right] + \left[ \boldsymbol{\alpha} / y_{\mathrm{ac}}^{\mathrm{E}} + \boldsymbol{\eta} \right] \cdot (y_{\mathrm{ac}}^{\mathrm{E}} J_{\mathrm{ac}}) + (\dots) \\
&\equiv \boldsymbol{\eta} J_{\mathrm{E,r}} + \boldsymbol{\eta}^{(\mathrm{f})} J_{\mathrm{E,f}} + (\dots)
\end{aligned}
\tag{N2.34}
$$

Such flux decomposition depends on the value of a parameter $y_{\mathrm{ac}}^{\mathrm{E}}$. We found that with the value $y_{\mathrm{ac}}^{\mathrm{E}} = 5.5$, the combination $\boldsymbol{\eta}^{(\mathrm{f})} \equiv \boldsymbol{\alpha}/y_{\mathrm{ac}}^{\mathrm{E}} + \boldsymbol{\eta}$ no longer contains the negative entries of TCA reactions that were present in $\boldsymbol{\alpha}$. Upon inspection of the vector components, we found that $\boldsymbol{\eta}^{(\mathrm{f})}$ satisfies the reaction stoichiometry of the aerobic fermentation pathway, (N2.33), up to a proportionality constant. The value $y_{\mathrm{ac}}^{\mathrm{E}} = J_{\mathrm{E,f}}/J_{\mathrm{ac}} = 5.5$ implies that acetate is coupled to the production of 5.5 units of ATP.

In the decomposition (N2.34), we multiplied the acetate demand flux $J_{\mathrm{ac}}$ by $y_{\mathrm{ac}}^{\mathrm{E}}$: this is convenient because this quantity corresponds to the generated ATP, rather than the acetate. With this normalization, the flux mode $\boldsymbol{\eta}^{(\mathrm{f})}$ produces 1 mmol ATP/$\mathrm{g_{DW}h}$ . We thus interpret $J_{\mathrm{E,f}} \equiv y_{\mathrm{ac}}^{\mathrm{E}} J_{\mathrm{ac}}$ as the energy produced by the fermentation pathway, and $J_{\mathrm{E,r}} = J_{\mathrm{ATPM}} - J_{\mathrm{E,f}}$ as the energy produced by the respiration pathway.

**2.3.2 Coupling energy with biosynthetic pathways.** After modifying the flux mode $\boldsymbol{\alpha}$, associated to acetate excretion, into the flux mode $\boldsymbol{\eta}^{(\mathrm{f})}$, associated to energy production via aerobic fermentation, we analyzed the flux components associated to biosynthesis of biomass building blocks. We found that these flux components contain both positive and negative components, often for TCA cycle reactions. In this case, the sign-mismatch presumably originates from a combination of three terms: the contribution associated to the production of energy via respiration, $J_{\mathrm{E,r}}$, the flux associated to the generation of carbon skeletons for biomass precursors, and the flux associated to the compensation of energy production or consumption associated to the biosynthesis of biomass components.

In order to disentangle these terms, we first identified AKGDH as a reaction purely associated to energy production, consistently with previous work [4, 18]. We confirmed this by observing that the AKGDH flux is zero in FBA simulations of anaerobic growth on glucose, indicating that it is not coupled to the biosynthesis of biomass precursors. Hence, we linearly combined the demand fluxes associated to respiration, $J_{\mathrm{E,r}}$, and to the biosynthesis of biomass precursors, $J_{\kappa}^{\mathrm{BM}}$, in order to remove the dependence of the AKGDH flux components on the biosynthetic demand fluxes. Following the same steps as in the previous case, we rewrite Eq. (N2.1) as:

$$\mathbf{v} = \boldsymbol{\eta}^{(\mathrm{f})} J_{\mathrm{E,f}} + \boldsymbol{\eta} J_{\mathrm{E,r}} + \sum_{\kappa} \boldsymbol{\beta}^{(\kappa)} J_{\kappa}^{\mathrm{BM}} \tag{N2.35}$$

$$= \boldsymbol{\eta}^{(\mathrm{f})} J_{\mathrm{E,f}} + \boldsymbol{\eta}\Big[J_{\mathrm{E,r}} - \sum_{\kappa} y_{\kappa}^{\mathrm{E}} \cdot J_{\kappa}^{\mathrm{BM}}\Big] + \sum_{\kappa}\Big[\boldsymbol{\beta}^{(\kappa)} + y_{\kappa}^{\mathrm{E}} \cdot \boldsymbol{\eta}\Big] J_{\kappa}^{\mathrm{BM}} \tag{N2.36}$$

$$= \boldsymbol{\eta}^{(\mathrm{f})} J_{\mathrm{E,f}} + \boldsymbol{\eta} J'_{\mathrm{E,r}} + \sum_{\kappa} \left(\boldsymbol{\beta}^{(\kappa)}\right)' J_{\kappa}^{\mathrm{BM}} \tag{N2.37}$$

For each biomass component $\kappa$, we set the value of $y_{\kappa}^{\mathrm{E}}$ such that the AKGDH flux component of $\left(\boldsymbol{\beta}^{(\kappa)}\right)'$ is zero. The vectors $\left(\boldsymbol{\beta}^{(\kappa)}\right)'$ will now produce or consume energy in association to the production of the biomass component. The term $y_{\kappa}^{\mathrm{E}} \cdot J_{\kappa}^{\mathrm{BM}}$ represents the energy flux produced or consumed in association to the synthesis of biomass precursor $\kappa$, which is removed from the "respiration" energy flux, $J_{\mathrm{E,r}}$. Therefore, the new respiration flux $J'_{\mathrm{E,r}} = J_{\mathrm{E,r}} - \sum_{\kappa} y_{\kappa}^{\mathrm{E}} \cdot J_{\kappa}^{\mathrm{BM}}$ no longer includes the contribution from the biosynthetic pathways, and genuinely reflects the activity of the energy-producing respiration pathway. This flux is referred to simply as $J_{\mathrm{E,r}}$ in the rest of the note and in the Main Text.

In sum, after the flux coupling process, the ATPM flux $J_{\mathrm{ATPM}}$ can be expressed as the sum of three terms, corresponding to the energy flux from respiration, ($J_{\mathrm{E,r}}$), the energy flux from aerobic fermentation ($J_{\mathrm{E,f}}$), and the total energy contribution from the biosynthetic pathways, ($J_{\mathrm{BM}} = \sum_{\kappa} J_{\kappa}^{\mathrm{BM}}$). These three terms are shown in Supp. Fig. S4A in both carbon and translation limitation.

**2.3.3 Functional coupling and negative components.** The results above can be summarized in a general form using the coupling matrix $C$ introduced before. The starting flux decomposition makes use of the demand fluxes ($J_{\mathrm{E}}, J_{\mathrm{ac}}, J_1^{\mathrm{BM}}, \ldots J_N^{\mathrm{BM}}$) associated to the flux modes $\boldsymbol{\eta}, \boldsymbol{\alpha}, \boldsymbol{\beta}_1, \ldots \boldsymbol{\beta}_N$). The redefined demand fluxes and the coupling matrix take the form:

$$\begin{pmatrix} J_{\mathrm{E,r}} \\ J_{\mathrm{E,f}} \\ J_1^{\mathrm{BM}} \\ \vdots \\ J_N^{\mathrm{BM}} \end{pmatrix}_{\mathrm{new}} = \begin{pmatrix} 1 & -y_{\mathrm{ac}}^{\mathrm{E}} & -y_1^{\mathrm{E}} & \ldots & -y_N^{\mathrm{E}} \\ 0 & y_{\mathrm{ac}}^{\mathrm{E}} & 0 & \ldots & 0 \\ 0 & 0 & 1 & & 0 \\ \vdots & \vdots & & \ddots & \vdots \\ 0 & 0 & 0 & \ldots & 1 \end{pmatrix} \begin{pmatrix} J_{\mathrm{E}} \\ J_{\mathrm{ac}} \\ J_1^{\mathrm{BM}} \\ \vdots \\ J_N^{\mathrm{BM}} \end{pmatrix}_{\mathrm{old}} \tag{N2.38}$$

Instead, the flux modes (organized in a row vector) transformed with the inverse matrix $C^{-1}$ as follows:

$$\begin{pmatrix}\boldsymbol{\eta} & \boldsymbol{\eta}^{(\mathrm{f})} & \boldsymbol{\beta}^{(1)} & \dots & \boldsymbol{\beta}^{(N)}\end{pmatrix}_{\mathrm{new}} = \begin{pmatrix}\boldsymbol{\eta} & \boldsymbol{\alpha} & \boldsymbol{\beta}^{(1)} & \dots & \boldsymbol{\beta}^{(N)}\end{pmatrix}_{\mathrm{old}} \begin{pmatrix} 1 & 1 & y_1^{\mathrm{E}} & \dots & y_N^{\mathrm{E}} \\ 0 & 1/y_{\mathrm{ac}}^{\mathrm{E}} & 0 & \dots & 0 \\ 0 & 0 & 1 & & 0 \\ \vdots & \vdots & & \ddots & \vdots \\ 0 & 0 & 0 & \dots & 1 \end{pmatrix} \quad \text{(N2.39)}$$

Here, the coefficients $y_{\mathrm{ac}}^{\mathrm{E}}$ and $y_{\kappa}^{\mathrm{E}}$ ($\kappa = 1, \dots, N$) are set so that the AKGDH reaction is uniquely assigned to aerobic respiration, as described above. Note that the flux mode corresponding to energy production, $\boldsymbol{\eta}$, has not changed. However, the corresponding demand flux has, together with its interpretation: the redefined flux component $\mathbf{v}^{(\mathrm{E,r})} \equiv \boldsymbol{\eta} J_{\mathrm{E,r}}$ now only corresponds to energy production associated to the TCA flux, excluding explicitly the energy flux associated to aerobic fermentation or coupled to biosynthesis. Instead, all other flux modes have been affected by the coupling of acetate or biomass production and energy.

With this redefinition of the flux components, we observed a marked reduction in sign-mismatch, as quantified by the values of the mixed functional fraction $F_i^{\mathrm{mix}}$, Eq. (N2.31) across all active reactions. These values are plotted in Supp. Fig. S2 against the magnitude of the corresponding fluxes, $|v_i|$, to highlight large fluxes with large negative entries (indicated by large $F_i^{\mathrm{mix}}$ values). Panels A through C show the original state. We notice large values of $F_i^{\mathrm{mix}}$ for the majority of the reactions carrying large ($\gtrsim$ 5mmol/$\mathrm{g_{DW}}$h) fluxes, including reactions belonging to the electron transport chain (ETC) pathway, ATP synthase (ATPS), TCA cycle, glycolysis, penthose phosphate pathway (PPP) and major transport fluxes (water, oxygen, carbon dioxide). The mixed components $F_i^{\mathrm{mix}}$ drop substantially after decoupling the respiration and fermentation pathways, as well as decoupling energy and biomass production (Supp. Fig. S2D-F). While the negative components for energy-producing pathways (ETC, TCA) are mostly cancelled by this coupling process, the mixed fractions of some reactions stay at large levels. These reactions include reactions in glycolysis (e.g. DHAPT) and in the penthose phosphate pathway (e.g. G6PDH2h, GND). This reflects what is also seen in the example shown in Fig. N2.1b: since the reactions $v_1$ and $v_3$ are set by the energy and biomass demand, respectively, the flux through $v_2$ has to match the difference between consumption and production of the intermediate metabolite.

##### 2.4 Application to anaerobically grown cells

In the case of anaerobic growth, we followed the same procedures described above, with a few differences. Similarly to what done in the case of aerobic growth, the flux components associated to the ATP maintenance flux $J_{\mathrm{ATPM}}$ and the succinate excretion flux $J_{\mathrm{succ}}$ were combined into two functional modes associated to energy production. In this case, the resulting functional components were associated to two modes of mixed acid fermentation, illustrated in Supp. Fig. S3G. For simplicity, instead of considering individual functional modes for the synthesis of each biomass component, we considered the synthesis of biomass precursors in aggregate. This led to a decomposition of the metabolic in three functional components (the two fermentation modes and the biosynthesis of all biomass precursors). In order to minimize the mixed functional components across the network, we subtracted the flux component describing mixed acid fermentation via ethanol production (associated originally to the ATP maintenance constraint) from the biomass-associated flux components, so that none of the ethanol excretion was associated to biomass biosynthesis. This led to a drastic reduction of mixed functional components across the network, as shown in Supp. Fig. S3H-I.

### Supplementary Note 3: Metabolite balance and mixed functions

Insights into the origin of sign-mismatch among the flux components can be obtained by studying the metabolic fluxes from a metabolite-centric perspective, and analyzing how the total production and consumption fluxes of each metabolite are associated to individual flux components. This analysis resembles the well-known thermodynamic relations between forward and backward flux components [19], which we summarize below.

#### 3.1 Thermodynamics flux decomposition

Consider the mass balance equation for a metabolite $m$:

$$\frac{\mathrm{d}[m]}{\mathrm{d}t} = \sum_i S_{m,i} v_i \ . \tag{N3.1}$$

The metabolic fluxes $v_i$ can be split into the forward ($v_i^{(+)} \geq 0$) and backward ($v_i^{(-)} \leq 0$) components as $v_i = v^{(+)} + v^{(-)}$; the ratio between the forward and backward fluxes is set by the variation of free energy of the reaction, $|v_i^{(+)}/v_i^{(-)}| = \exp\bigl(-\Delta_r G/RT\bigr)$. By labeling the two components with a single index, $\kappa = \{+, -\}$, we can rewrite the mass balance constraint as:

$$\frac{\mathrm{d}[m]}{\mathrm{d}t} = \sum_{i\kappa} S_{m,i} v_i^{(\kappa)} \ . \tag{N3.2}$$

The characteristic turnover rate of the metabolite $m$ is determined by the total magnitude of production and consumption fluxes. We define the total production flux as the sum of all positive terms in the r.h.s. of Eq. (N3.2)

$$P_m = \sum_{i\kappa : S_{m,i} v_i^{(\kappa)} > 0} S_{m,i} v_i^{(\kappa)} \tag{N3.3}$$

The turnover rate of the metabolite is then given by $k_m = P_m/[m]$. At steady state, production and consumption fluxes need to match, leading to:

$$P_m = \frac{1}{2} \sum_{i\kappa} \left| S_{m,i} v_i^{(\kappa)} \right| \tag{N3.4}$$

This expression can be compared to the analogous quantity computed using only the net fluxes:

$$P_m^{\text{net}} = \frac{1}{2} \sum_i |S_{m,i} v_i| \tag{N3.5}$$

With a few lines of algebra, it is possible to express the relationship between $P_m^{\text{net}}$ and $P_m$ as follows:

$$P_m - P_m^{\text{net}} = \frac{1}{2} \sum_i F_i^{\text{therm}} \sum_\kappa |S_{m,i} v_i^\kappa| \geq 0 \ , \tag{N3.6}$$

where $F_i^{\text{therm}} \equiv 1 - |v_i| / \sum_\kappa |v_i^{(\kappa)}|$ is between 0 and 1, and it is set by the thermodynamic driving force of the reaction:

$$F_i^{\text{therm}} = \frac{2e^{|\Delta_r G|/RT}}{1 + e^{|\Delta_r G|/RT}} \tag{N3.7}$$

Hence, if a metabolite is produced or consumed by reactions far from equilibrium, $F_i^{\text{therm}} \sim 0$ and $P^{\text{net}} \sim P_m$; instead, for reactions close to equilibrium, $F_i^{\text{therm}} \sim 1$ and $P^{\text{net}}$ is much lower than the true production flux $P_m$.

##### 3.2 Functional flux decomposition

The function-based flux decomposition presented in this work present clear analogies with the the decomposition of the fluxes into forward and backward components. Each functional component satisfies the mass-balance constraints, similarly to the thermodynamic components in Eq. (N3.2). Three main differences should be noted, though. First, the total flux can be split into only two components, forward and backward, as opposed to a multitude of possible functional components; second, forward and backward flux components are related by thermodynamics, while no similar relation exists between the functional flux components. On the other hand, the forward and backward flux components $\mathbf{v}^{(+)}$ and $\mathbf{v}^{(-)}$ do not satisfy the mass balance constraints (only their sum does), while the mass balance constraints are satisfied by each individual functional flux component, i.e. $\sum_m S_{m,i} v_i^{(\kappa)} = 0$ for every metabolite $m$ and function $\kappa$. Third, the total production flux $P_m$ can be associated to a physical timescale in the thermodynamic case, while no intuitive meaning exists in the flux decomposition case. Still, $P_m$ and $P_m^{\text{net}}$ represent useful quantities for the functional characterization of the metabolic network from the perspective of metabolite balance.

In the context of a functional flux decomposition, the expression Eq. (N3.6) shows that the difference between $P^{\text{net}}$ and $P_m$, which we term $P_m^{\text{mix}}$, is determined by the magnitude of fluxes of reactions with large mixed flux components, as determined by the mixed functional component $F_i^{\text{mix}} = 1 - |v_i| / \sum_\kappa |v_i^{(\kappa)}|$:

$$P_m^{\text{mix}} \equiv P_m - P_m^{\text{net}} = \frac{1}{2} \sum_i F_i^{\text{mix}} \sum_\kappa |S_{m,i} v_i^\kappa| \ , \tag{N3.8}$$

In fact, this difference is simply related to the magnitude of the negative flux components. It is easy to check that

$$|v_i| = \sum_\kappa |v_i^{(\kappa)}| - \sum_{\kappa: v_i^{(\kappa)} \cdot v_i < 0} 2|v_i^{(\kappa)}| \ . \tag{N3.9}$$

By multiplying both sides by $|S_{m,i}|$, and summing over all reactions $i$, we obtain:

$$\sum_{i,\kappa: v_i^{(\kappa)} \cdot v_i < 0} |S_{m,i} v_i^{(\kappa)}| = P_m - P_m^{\text{net}} = P_m^{\text{mix}} \ . \tag{N3.10}$$

This relation provides a convenient framework to study the presence of sign-mismatch reactions across the whole network. Metabolites produced or consumed by reactions with sign-mismatched flux components are characterized by a non-vanishing $P_m^{\text{mix}}$. Supplementary Figure S2G shows a scatter plot of $P_m^{\text{mix}}$ versus $P_m$ for several metabolites, and illustrates how the majority of metabolites in the upper glycolytic pathway and the penthose phosphate pathway are associated to reactions with mixed functional fractions. Further insights can be obtained by focusing on different flux topologies around the metabolite of interest. To do so, we define the functional overlap of two reactions $r$ and $s$ as:

$$q_{ij} = 1 - \frac{1}{2} \sum_\kappa \left| F_i^{(\kappa)} - F_j^{(\kappa)} \right| \tag{N3.11}$$

In this expression, mixed flux components are included, and the other functional components are determined as described in Section 2.2.5. Another useful quantity is a binary indicator that matches the relative sign of the production or consumption of the metabolite $m$ by two reactions $r$ and $t$:

$$b_m^{ij} = \text{sign}\left( (S_{m,i} v_i) \cdot (S_{m,i} v_j) \right) \tag{N3.12}$$

Using these quantities, it is possible to provide a partial functional classification of the possible flux topologies of metabolite balance, illustrated in Supplementary Figure S2H-I. Consider the set of reactions either producing or consuming a metabolite $m$ for which $P_m^{\text{mix}} > 0$. We can identify two key topologies:

- **Class I topology.** If two reactions $i$ and $j$ have both mixed functional components, large overlap

$q_{ij} \sim 1$ and opposite production sign, $b_m^{ij} = -1$, then these reactions two form a linear pathway, with the metabolite $m$ being an intermediate metabolite in the pathway.

- **Class II topology.** If only one reaction $i$ has a mixed component, $F_i^{\mathrm{mix}} > 0$, then its flux is adjusted to keep the concentration of $m$ steady as it is being produced and consumed by reactions associated to different functional components. Similarly, if two reactions $i$ and $j$ have both mixed functional components, large overlap $q_{ij} \sim 1$ and the same production sign, $b_m^{ij} = +1$, then they effectively behave as a unique reaction.

In particular, class II topologies provide the key insight into the emergence of mixed functions in the metabolic network. In these cases, anaplerotic reactions are needed to makes sure that the concentration of currency/intermediate metabolites is stable upon different demand fluxes. Since the flux components are mass-balanced, the presence of sign-mismatch in these reactions causes a cascade of flux-mismatches across other reactions (e.g. via linear pathways, i.e. class I topologies).

The topologies in Supplementary Figure S2H-I were classified as follows. First, for each metabolite, all associated reactions ($S_{m,i} v_i \neq 0$) were ranked by $F_i^{\mathrm{mix}}$, so that $F_1^{\mathrm{mix}} \geq F_2^{\mathrm{mix}} \geq \ldots$. Then:

- If $F_1^{\mathrm{mix}} \geq 0.005$, $F_2^{\mathrm{mix}} \geq 0.005$, $q_{12} > 0.85$ and $b_m^{12} = -1$, then the metabolite is associated with a class I topology.

- If $F_1^{\mathrm{mix}} \geq 0.1$ and $F_1^{\mathrm{mix}} \leq 0.05$ (i.e. there is only one predominant reaction with mixed flux components), then the metabolite is associated with a class II topology.

- If $F_1^{\mathrm{mix}} \geq 0.005$, $F_2^{\mathrm{mix}} \geq 0.005$, $q_{12} > 0.85$ and $b_m^{12} = +1$, then the metabolite is associated with a class II topology.

### Supplementary Note 4: Energy and carbon costs of biosynthetic pathways

This section describes the calculation of the carbon costs and yields shown in Main Text Fig. 4 and reported in Supp. File S6, as well as several details on the interconversion between electron acceptors, protons and ATP.

#### 4.1 Calculation of specific costs and yields for biomass components

We first consider the naïve flux decomposition obtained from the perturbations to the FBA solution, before performing the coupling described in the previous section. In this case, each flux mode $\boldsymbol{\xi}^{(\kappa)}$ represent genome-scale pathways uniquely associated to a single demand flux $J_\kappa$. The demand flux can either correspond to the production of a given metabolite (a biomass component, or extracellular acetate when constraining its production) or the flux of a constrained reaction (e.g. ATP "maintenance" hydrolysis flux).

For a single reaction $i$, the value $\xi_i^{(\kappa)}$ represent the reaction flux associated with function $\kappa$, expressed in units of $J_\kappa$. The flux modes associated to exchange fluxes are particularly interesting. Consider for example the important case of the glucose uptake flux, with flux mode $\xi_{\text{glc}}^{(\kappa)}$. This coefficient indicates how many molecules of glucose are needed per each molecule of the biomass component $\kappa$, i.e. represent the glucose cost of biomass precursor $\kappa$. Its inverse, $1/\xi_i^{(\kappa)} = J_\kappa/v_i^{(\kappa)}$ represents the yield of biomass component $\kappa$ per unit of intaken glucose, and thus represents the carbon efficiency of the pathway.

**4.1.1 Coupling biomass production to energy.** We now consider how the flux coupling procedure impacts the costs. Coupling the demand fluxes $J_\gamma \to \sum_\delta C_{\gamma\delta} J_\delta$ leads to a redefinition of the flux modes, $\boldsymbol{\xi}^{(\gamma)} \to \sum_\delta \boldsymbol{\xi}^{(\delta)} C_{\delta\gamma}^{-1}$. In particular, with our choice of the coupling matrix, the biosynthetic and the acetate-producing flux components are coupled with energy production as $\boldsymbol{\xi}^{(\gamma)} \to \boldsymbol{\xi}^{(\gamma)} + y_\gamma^E \boldsymbol{\eta}$. The flux-coupling allows to minimize the impact of sign-mismatches among the flux components, and therefore to improve the interpretability of the pathways described by the flux modes. However, the combination of different demand fluxes also means that each flux mode is no longer uniquely associated to the production of the biomass component of interest. After coupling the biosynthesis of the $\kappa$-th biomass precursor to energy, the flux mode will represent the fluxes associated to the production of 1 mmol/$\text{g}_{\text{DW}}$h of biomass precursor, and $y_\kappa^E$ mmol/$\text{g}_{\text{DW}}$h of ATP. Thus, the parameters $y_\kappa^E$ represent the energy cost (or production) per unit of biomass precursor produced.

Similarly, the glucose specific cost $\xi^{(\kappa)}_{\text{glc}}$ not only reflects the amount of carbon atoms present in the product of the pathway, but also takes into account the carbon flux associated to energy production/consumption. Therefore, the glucose cost is not proportional to the number of carbon atoms in each molecule (Fig. N4.1A). Instead, the flux modes associated to nitrogen (ammonium, $\xi^{(k)}_{\text{nh4}}$), phosphorus (phosphate, $\xi^{(k)}_{\text{pi}}$) and sulfur uptake (sulfate, $\xi^{(k)}_{\text{so4}}$) match precisely the elemental composition of each molecule (Fig. N4.1B-D).

**4.1.2 Combined costs of multiple functions.** If one is only interested in the overall costs of groups of biomass precursors, e.g. all amino acids, or the entire set of biomass precursors, it is possible to compute the average costs and yields in a straightforward manner. Consider the set $A$ of biomass precursors of which we want to calculate the average yields. A combined flux $J_A$ associated to the production of all metabolites has to be defined as $J_\kappa = c_\kappa J_A$; the parameters $c_\kappa$ depend on the relative magnitude of the demand fluxes $J_\kappa$ and on the overall normalization of $J_A$. Then, the flux decomposition can be rewritten as:

$$\mathbf{v} = \sum_\kappa \boldsymbol{\xi}^{(\kappa)} J_\kappa \tag{N4.1}$$

$$= \sum_{\kappa \in A} \boldsymbol{\xi}^{(\kappa)} J_\kappa + \sum_{\kappa \notin A} \boldsymbol{\xi}^{(\kappa)} J_\kappa \tag{N4.2}$$

$$= \left( \sum_{\kappa \in A} \boldsymbol{\xi}^{(\kappa)} c_\kappa \right) J_A + \sum_{\kappa \notin A} \boldsymbol{\xi}^{(\kappa)} J_\kappa \tag{N4.3}$$

$$\equiv \boldsymbol{\xi}^{(A)} J_A + \sum_{\kappa \notin A} \boldsymbol{\xi}^{(\kappa)} J_\kappa \tag{N4.4}$$

The newly defined flux mode $\boldsymbol{\xi}^{(A)}$ accounts for the production of the biomass components in the proportions sets by the coefficients $c_\kappa$, and can thus be used for the computation of specific fluxes (in units of $J_A$) and yields.

#### 4.2 Calculation of ATP equivalents for electron carriers

To simplify the metabolic network, a common approximation is to assume the equivalence between the production of electron carriers, NAD(P)H and FADH, and ATP. Other common metabolites, such as water and protons, are often neglected. We computed the effective conversion coefficients within the flux decomposition by introducing a small "dissipation" flux, and evaluating the additional ATP flux necessary

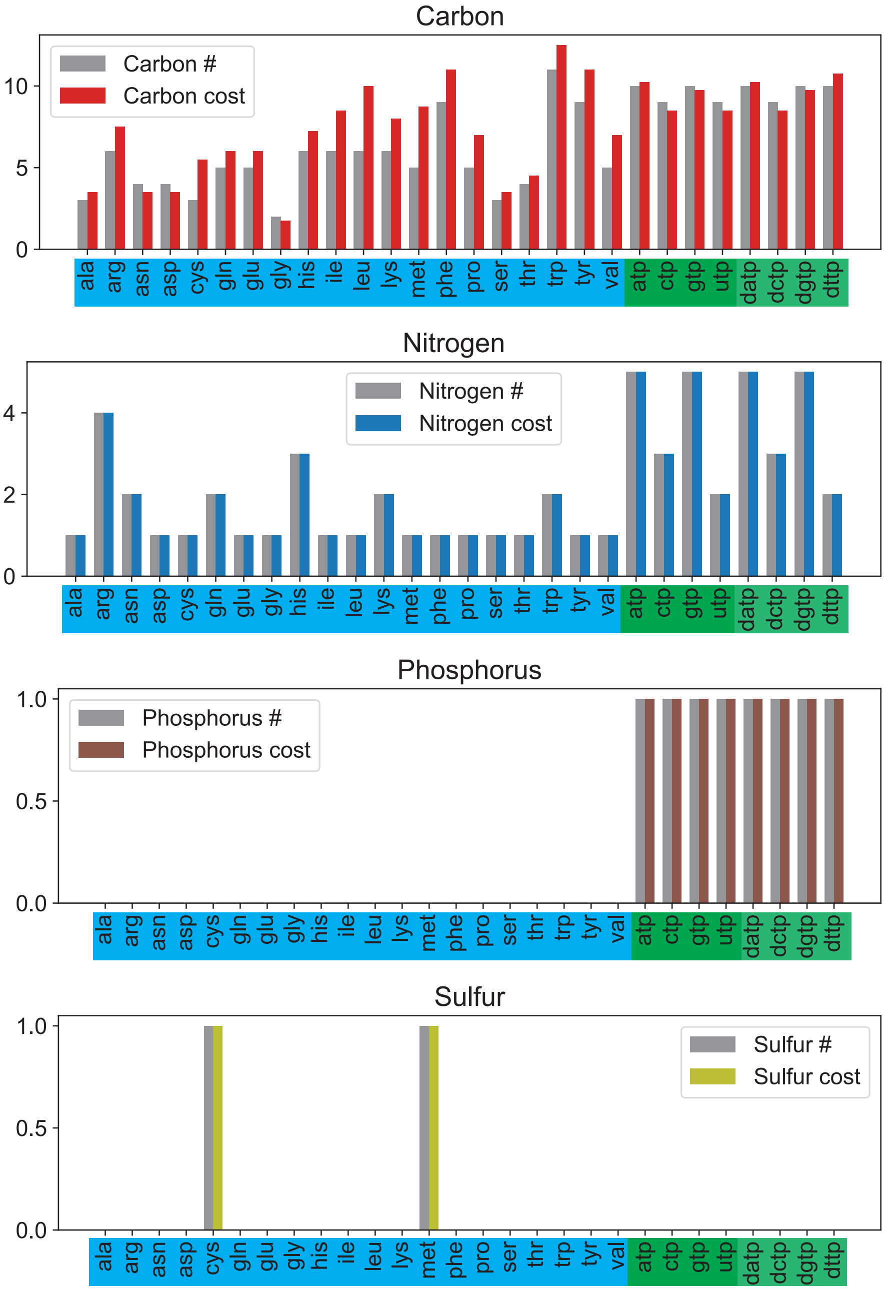

**Figure N4.1.** Specific cost (colored bars) of amino acids and nucleic acids in glucose minimal media. (A) Specific glucose intake in units of carbon atoms, $6 \times \xi_{\text{glc}}^{(\kappa)}$. (B) Specific ammonia uptake, $\xi_{\text{nh4}}^{(\kappa)}$. (C) Specific phosphate uptake, $\xi_{\text{pi}}^{(\kappa)}$. (D) Specific sulfate uptake $\xi_{\text{so4}}^{(\kappa)}$. In all panels, grey bars indicate the elemental composition of the same biomass building block, i.e. the number of C/N/P/S atoms per molecule. The cost of N/P/S always matches the elemental composition of the biomass component, while significant differences are seen when comparing the C cost to the C content of each molecule. This mismatch is mostly due to the $CO_2$ exchanged alongside the production of the metabolite, e.g. because of (de)carboxylase reactions in energy production (e.g. TCA) and anaplerotic (e.g. PPC) pathways.

to compensate the fluxes by computing the derivative of the optimal fluxes with respect to the dissipation flux. Importantly, this reaction needs to be both mass and charge balanced in order to yield consistent results. For this reason, oxygen is explicitly added as electron acceptor in the reactions. The dissipation reactions and the associated energy costs are summarized in Table S3. Specifically, when calculating these conversion factors, we first run flux balance analysis (FBA) using the canonical constraints and objective function: minimizing the carbon (glucose) intake flux in a fixed growth rate, with the dissipation flux set to zero. Next, we constraint the glucose intake reaction to the minimized flux from the previous FBA, set the dissipation flux to a small value (the derivative), and rerun FBA by maximizing the ATPM reaction. We took the difference between the two set of FBA solutions, and divided this by the derivative, which gave us the cost of each dissipation reaction. The costs show a good agreement with the conversion coefficients generally assumed for NAD(P)H (1 NAD(P)H = 2 ATP). However, our calculation of the conversion coefficients between FADH and ATP (1.75:1) is higher than what was generally assumed (1:1). We also found that the leakage of protons from the cytoplasm to the periplasm is associated to an additional cost of 0.25 ATP. Failing to account for the proton balance will hence affect the evaluation of the energetic costs for the production of biosynthetic precursors. See Supp. File S6 for the reactions that are involved in the balancing between those electron carriers and ATP.

#### Supplementary Note 5: Functional decomposition of the proteome

The aim of this section is to describe how to connect the experimental protein abundances to the functional decomposition described in Supplementary Note 2, and how to the use the proteome functional decomposition to coarse grain the cellular proteome. To do so, we adopted the protein abundance data from Mori et al. [3] and Wu et al. [20]. In particular, we used protein mass fractions $\phi_p^{\text{exp}}$ (mass of the $p$-th protein per total protein mass), as opposed to protein number fractions (proportional to concentrations), for three reasons. (1) The total protein mass (per optical density unit, or per cell dry weight) can be measured relatively straightforwardly, e.g. with biochemical assays (Biuret); instead, the total number of proteins cannot be measured directly. (2) Since most cellular ribosomes are elongating except at very slow growth [21], the concentration of ribosomes limits the total number of residues polymerized per unit of time, i.e. the total synthesis flux of protein mass. Therefore, protein mass is a better measure of the protein burden associated to ribosome utilization compared to protein copy number (or concentration). (3) Mass spectrometry-based proteomics typically have a lower detection limit set by the protein mass, not the protein number [3].

##### 5.1 Reaction-associated protein mass fractions

The relationship between reactions and proteins is not always straightforward. Different proteins can catalyze the same reaction and, in turn, single enzymes can catalyze different reactions. We define $S_i$ as the set of proteins associated to the $i$-th reaction; the associations can be easily generated using the gene-protein-reaction (GPR) matrix contained in the iML1515 *E. coli* model (including all genes associated to the reactions). It is also useful to introduce the indicator function $\chi_{ip}$ that summarizes the association between proteins and reactions as follows:

$$\chi_{ip} = \begin{cases} 1, & p \in S_i \\ 0, & p \notin S_i \end{cases} \tag{N5.1}$$

In order to associate a protein mass fraction $\phi_i$ to a single reaction $i$ based on the experimental protein abundances $\phi_p^{\text{exp}}$ we will adhere to the following guidelines: (1) If multiple proteins are associated to the same reaction, then we want to associate their protein abundances to the same reaction. (2) If a protein is associated to multiple reactions, we need to partition it abundance over them. In principle, we would like to allocate the protein abundance in proportion to the activity of the enzyme in each reaction. In the context of Michaelis-Menten kinetics, the concentration of enzyme-substrate complexes, which equals the ratio of reaction flux and turnover number of the reaction, would be the most direct proxy for the enzyme activity. Since *in vivo* turnover numbers are notoriously hard to estimate, we will consider for simplicity that the amount of enzymes associated to each reaction is proportional to the reaction flux, assume a simplest approximation of enzyme bound being proportional to the flux of the reaction.

The simplest way to partition protein abundances over the reactions satisfying these two requirements is to introduce a *protein distribution* matrix $Q_{ip}$ as follows:

$$\phi_i = \sum_p Q_{ip}\phi_p^{\text{exp}} \qquad Q_{ip} = \begin{cases} \dfrac{\chi_{ip}|x_i|}{\sum_j \chi_{jp}|x_j|} & \text{if } x_i \neq 0\ , \\ \\ 0 & \text{if } x_i = 0\ . \end{cases} \tag{N5.2}$$

With this choice, it is straightforward to check that the requirements described above are met. For example, if two proteins with mass fractions $\phi_1^{\text{exp}}$ and $\phi_2^{\text{exp}}$ both catalyze the same reaction with a flux $x_3$, then $Q_{31} = Q_{32} = 1$ and $\phi_3 = \phi_1^{\text{exp}} + \phi_2^{\text{exp}}$. If instead a protein with mass fraction $\phi_1^{\text{exp}}$ catalyzes two reactions with fluxes $x_1 = x_2 = 1$ mmol/$\text{g}_{\text{DW}}$h , then $\phi_1 = \phi_2 = \frac{1}{2}\phi_1^{\text{exp}}$. Note that the factor $\frac{1}{2}$ makes sure that the protein abundances are not over-counted when summing over reactions. It is easy to see that $\chi_p^{\text{MET}} \equiv \sum_i Q_{ip}$ is the indicator function for metabolism-associated proteins, i.e. it is 1 if the protein is associated to at least one metabolic reaction with nonzero flux, and zero otherwise. The total fraction of proteome associated to metabolism, $\phi_{\text{MET}}$, is obtained by summing over all metabolic reactions:

$$\phi_{\text{MET}} \equiv \sum_i \phi_i = \sum_{i,p} Q_{ip}\phi_p^{\text{exp}} = \sum_p \chi_p^{\text{MET}}\phi_p^{\text{exp}}\ . \tag{N5.3}$$

#### 5.2 Functional decomposition for the proteome

The functional decomposition described in the previous Supplementary Note can be used to partition protein abundances into components associated to each metabolic functions. For any given reaction $i$ and a given set of metabolic functions $\{\gamma\}$ (including among them the mixed component with magnitude $F_i^{\text{mix}}$), one can compute a partitioning $\sum_\gamma F_i^\gamma = 1$, with $F_i^\gamma$ specifying the contribution of reaction $i$ to the function $\gamma$. By using the functional decomposition and the relationship between reaction-associated protein mass fractions and experimental protein abundances, Eq. (N5.2), we can write:

$$\phi_i = \left(\sum_\gamma F_i^\gamma\right)\phi_i = \sum_\gamma \left(F_i^\gamma \phi_i\right) \equiv \sum_\gamma \phi_i^\gamma \ , \tag{N5.4}$$

where $\phi_i^\gamma \equiv F_i^\gamma \phi_i$ naturally defines the amount of proteins associated to reaction $r$ and metabolic function $\gamma$. By summing both sides of the equation over all metabolic reactions, we obtain a functional decomposition describing the allocation of metabolism-associated proteins over the metabolic functions:

$$\phi_{\text{MET}} = \sum_i \phi_i \equiv \sum_\gamma \phi_{\text{MET}}^\gamma \ , \tag{N5.5}$$

where $\phi_{\text{MET}}^\gamma = \sum_i \phi_i^\gamma$ represent "functional sectors" describing succinctly how the proteome is allocated by the cell to each metabolic function $\gamma$. Explicitly, in terms of the experimental proteins fractions $\phi_p^{\text{exp}}$:

$$\phi_{\text{MET}}^\gamma = \sum_{i,p} F_i^\gamma Q_{ip} \phi_p^{\text{exp}} = \sum_p \left[\sum_i F_i^\gamma Q_{ip}\right] \phi_p^{\text{exp}} \ . \tag{N5.6}$$

The term in square brackets, mapping experimental protein abundances to each function-specific protein sector, is obtained as the matrix product of the functional decomposition matrix, $F_i^\gamma$, and the protein distribution matrix $Q_{ip}$. It is important to note that in this approach a single protein generally belongs to multiple functional sectors: the only case in which it is associated to a single functional sector is when the protein only catalyzes one reaction, and the reaction has a unique function. The mass fractions $\phi_{\text{MET}}^\gamma$ describe succinctly how the proteome is allocated by the cell to different metabolic functions.

#### Supplementary Note 6: Models of protein allocation

The proteome composition of bacterial cells depends strongly on the growth conditions. However, several classes of proteins show striking linear relations with the growth rates of the cell [22]. Examples include the ribosomal proteins, which are linearly correlated with the growth rate, and catabolic genes, which are anticorrelated with the growth rate in carbon-limited growth. These linear relationships have inspired the formulation of several protein allocation models, of which the one presented in Scott *et al.* [23] is the prototypical example. This model was introduced to explain the striking linear relationships observed for the expression of ribosomes (proportional to the total RNA/protein mass ratio of the cells) in either nutrient- or translation-limited conditions.

The model described in Scott *et al.* can be entirely constructed based on the empirical relations displayed by the data. We will denote the ribosomal protein mass fraction as $\phi_{\mathrm{Rb}}$. Across nutrient limiting conditions, in absence of antibiotics, $\phi_{\mathrm{Rb}}$ satisfies a linear relation with the growth rate,

$$\phi_{\mathrm{Rb}} = \phi_{\mathrm{Rb},0} + \mu/\nu_R \ . \tag{N6.1}$$

The inverse of the slope $\nu_R$ is closely related to the maximum elongation rate of the ribosomes, and is therefore termed "translational capacity" of the cell. A mechanistic interpretation of Eq. (N6.1) is that $\phi_{\mathrm{Rb},0}$ represents an "inactive" fraction of ribosomes, and the rest an "active" part working at a rate set by $\nu_R$. (The interpretation of these terms has later evolved to include the effect of varying translation elongation rate [21] and ribosome-inhibiting proteins [20].) In conditions with increasingly higher concentrations of antibiotics, $\phi_{\mathrm{Rb}}$ satisfies a different linear relation with a negative slope,

$$\phi_{\mathrm{Rb}} = \phi_{\mathrm{Rb,max}} - \mu/\nu_C \ . \tag{N6.2}$$

The value of $\nu_C$ correlates with the cellular growth rates obtained supplying different nutrients to the cell, and is therefore termed "nutritional capacity" of the cell. Crucially, the value of $\phi_{\mathrm{Rb,max}}$ depends only weakly on the nutrient condition, and represents the maximal protein mass fraction which the cell can allocate to the ribosomes. By assuming that $\phi_{\mathrm{Rb,max}}$ is condition-independent[2], it is possible to express both the growth rate and the size of the ribosome-associated protein sector as a function of the

[2]This assumption breaks down at slow growth, $\mu \lesssim 0.5/h$, where $\phi_{\mathrm{Rb,max}}$ is reduced [21].

two parameters $\nu_C$ and $\nu_R$ as follows:

$$\mu(\nu_C, \nu_R) = \phi_{\max} \frac{\nu_C \nu_R}{\nu_C + \nu_R} \tag{N6.3}$$

$$\phi_{\mathrm{Rb}}(\nu_C, \nu_R) = \phi_{\mathrm{Rb},0} + \phi_{\max} \frac{\nu_R}{\nu_C + \nu_R} \tag{N6.4}$$

In these relations, $\phi_{\max} \equiv \phi_{\mathrm{Rb,max}} - \phi_{\mathrm{Rb},0}$ represents the maximum change in allocation allowed to the ribosomes. Overexpression of useless proteins was found to reduce $\phi_{\max}$ and affect both the growth rate and the ribosomal protein abundance according to the predictions of Eq. (N6.3) and (N6.4).

#### 6.1 Multiple protein sectors and regulation-based sectors

A notable consequence of Eq. (N6.3) and Eq. (N6.4) is that the changes in ribosomal protein mass fractions across growth rates constrain the expression of the rest of the proteome. Since the sum of all protein fraction is 1, the remainder of the proteome must obey relations specular to those satisfied by the ribosomal proteins. Indeed, constitutively expressed proteins were found to obey relations opposite to those of the ribosomal ones [23], suggesting the presence of global constraints on gene expression [24]. Furthermore, the interplay between the regulation of ribosomal proteins and other global regulatory systems was explored in a subsequent study [22], where the introduction of additional protein sectors accounted for the regulation on catabolic and anabolic genes, as determined by promoter reporters.

In order to explore the regulation of the proteome at the genome-scale, Hui et al. [25] defined empirically a partitioning of the proteome based on the response of each protein to three different perturbations. The first two perturbations are the C-limitation and R-limitation; these are the growth perturbations studied in this manuscript. A third "anabolic" (A-) limitation was imposed by titrating the cellular nitrogen uptake flux. Genes were assigned to protein sectors based on a binary classification of their regulations, either up- or downregulated in each of the three growth limitations. For example, proteins upregulated in carbon starvation and downregulated in the other two growth limitations (the behavior exhibited by many catabolic proteins) were assigned to the so-called C-sector. The three growth limitations give rise to $2^3 = 8$ possible proteins sectors, as summarized in Supp. Fig. S10-D.

Similarly to [22], the data was captured by a coarse-grained protein allocation model in which the expression of each protein sector was set by three phenomenological parameters related to the growth conditions of the cells: $\nu_C$, correlated to the carbon quality of the medium or the cellular uptake capacity; $\nu_A$, correlated to the cellular capacity of assimilating nitrogen; and $\nu_R$, the translational capacity of the cell. Each of these parameters sets the growth rate dependence of the corresponding protein sector ($\phi_C$, $\phi_A$ and $\phi_R$) in the growth limitations that do not specifically induce them (e.g. C-sector in non-C limitations).

##### 6.2 Energy protein allocation and the respiration-fermentation switch

A very different model was introduced in Basan et al. [18]. The model described the shift between respiration and aerobic fermentation as a consequence of protein allocation constraints. The model is based on three main equations, namely the balance of carbon and energy production and the sum rule on the protein mass fractions:

$$J_{C,r} + J_{C,f} = J_{C,in} - \beta\mu \tag{N6.5}$$

$$J_{E,r} + J_{E,f} = \sigma\mu \tag{N6.6}$$

$$\phi_{E,r} + \phi_{E,f} = \phi_{E,max} - b\mu \tag{N6.7}$$

Here, $J_{C,in}$ is the total carbon intake. The two parameters $\sigma$ and $\beta$ determine the carbon and energetic demand of the cell due to biosynthetic activities, and the total proteins allocated towards energy production is constrained to decrease at fast growth as dictated by the parameters $\phi_{E,max}$ and $b$. These three constraints are supplemented by constitutive relations set by pathway-specific parameters: the ATP production per unit of carbon ($e_r$ and $e_f$), and the ATP production per unit of proteome allocated ($\varepsilon_r$, $\varepsilon_f$). Using these relationships, the model takes the following form:

$$J_{E,r}/e_r + J_{E,f}/e_f = J_{C,in} - \beta\mu \tag{N6.8}$$

$$J_{E,r} + J_{E,f} = \sigma\mu \tag{N6.9}$$

$$J_{E,r}/\varepsilon_r + J_{E,f}/\varepsilon_f = \phi_{E,max} - b\mu \tag{N6.10}$$

Among these equations, the last two are the ones determining the fluxes through the individual pathways as a function of the growth rate. For example, solving for $J_{E,f}$ leads to:

$$J_{E,f} = \frac{(\varepsilon_r\sigma + b)\mu - \phi_{E,max}}{\varepsilon_f - \varepsilon_r} \tag{N6.11}$$

Given that the fermentation pathway produces more ATP per unit of invested proteome compared to the respiration one, $\varepsilon_f > \varepsilon_r$, leading to a positive correlation between the fermentation flux and the growth rate, as long as the growth rate is larger than the "acetate onset" value $\mu_{\rm ac}$ and lower than the growth rate at which the respiration flux goes to zero, $\mu_{\rm max}$:

$$\mu_{\rm ac} = \frac{\phi_{E,max}}{\varepsilon_r \sigma + b}\ , \quad \mu_{\rm max} = \frac{\phi_{E,max}}{\varepsilon_f \sigma + b} \tag{N6.12}$$

Note that the carbon efficiency does not play any role in the calculation above; instead, Eq. (N6.5) and Eq. (N6.8) define the growth yield (biomass accumulated per unit of intaken carbon) as a function of the energetic fluxes, and thus to calculate the growth rate as a function of the intaken carbon flux.

While this model is able to generate a variety of predictions on how the "acetate line" is affected by a variety of perturbations (including additional energy expenditure, or wasteful protein allocation), it is manifestly incompatible with the coarse-grained models formulated in You et al. [22] and Hui et al. [25], because these models do not include an energy-associated protein sector. Instead, in Hui et al. [25], proteins associated to the central carbon pathways are classified in a variety of ways depending on their regulation: many TCA proteins are upregulated in carbon-limited conditions, and are therefore part of the C- and S-sectors; several glycolytic proteins are instead upregulated in anabolic limitation, and belong to the A-sector. This problem reflects the fact that different schemes for the generation of protein sectors from proteomics data generally lead to incompatible results: proteins with similar responses to growth perturbations can have different biological functions, and vice versa proteins with the same biological functions can be differentially regulated.

#### 6.3 A complete function-based protein allocation model

The functional classification introduced in this work allows us to introduce a coarse-grained protein allocation model for protein sectors with definite functions, as opposed to regulation; additionally, the model includes that from Basan et al. [18], and is thus able to predict the "acetate shift". We will first describe the mathematical formulation of the model, and discuss its biological interpretation in the next section.

As described in the Main Text, the model includes five protein sectors, each associated with specific biological functions. Metabolic proteins, whose function was determined using the functional decomposition introduced in this work, were minimally divided in two groups: an "energy" sector of protein mass fraction $\phi_E$, and a biosynthetic sector, including enzymes producing precursors of cellular biomass, with mass fraction $\phi_{\rm bm}$. The rest of the proteome was classified using two condition-dependent sectors defined in terms of GO-terms ("translation" and "foraging", with mass fractions $\phi_{\text{tsl}}$ and $\phi_{\text{for}}$), and a fixed protein sectors with "housekeeping" functions with fraction $\phi_{\text{hk}}$. The mathematical model is based on the following constraints on the protein mass fractions:

$$\phi_E = \phi_{E,0} + \kappa_C \mu/\nu_C + \kappa_R \mu/\nu_R \tag{N6.13}$$

$$\phi_{\text{bm}} = \phi_{\text{bm},0} + \mu/\nu_B \tag{N6.14}$$

$$\phi_{\text{tsl}} = \phi_{\text{tsl},0} + \mu/\nu_R \tag{N6.15}$$

$$\phi_{\text{for}} = \phi_{\text{for},0} + \mu/\nu_C \tag{N6.16}$$

Here, the protein fractions $\phi_{E,0}$, $\phi_{\text{bm},0}$, $\phi_{\text{tsl},0}$, $\phi_{\text{for},0}$, as well as the share of housekeeping proteins $\phi_{\text{hk}}$, are considered fixed. Several parameters determine how each protein sector changes with the growth condition. The constant $\nu_B$ determines the slope of the "biomass" sector against the growth rate; the parameter $\nu_C$ measured the quality of the carbon source, while the parameter $\nu_R$ measured the translational capacity of the cell. In particular, $\nu_C$ and $\nu_R$ are the "environmental" parameters that reflect the growth condition of the cell. Finally, the parameters $\kappa_C$ and $\kappa_R$ determine the response of the E-sector to changes in either the carbon quality or the translational capacity of the cell.

The simplest interpretation of Eq. (N6.14) – (N6.16), consistent with the linear dependencies on the growth rate observed for these sector, is that the offsets $\phi_{i,0}$ are inactive, and the reaction fluxes are proportional to the remainder of the proteome $\Delta\phi_i = \phi_i - \phi_{i,0}$ via a rate $k_i$; for the translation sector, this rate matches well that of ribosome elongation. Mechanistically, such offsets can be caused by either the inactivation of the enzymes, e.g. the end product inhibition acting in several biosynthetic pathways, or a reduced thermodynamic driving force for reactions close to equilibrium, often due to low substrate levels [26]. The constraint on the energy protein sector, Eq. (N6.13), is more involved, and will be discussed later below in Section 6.3.1. By definition, the sum of all protein fractions is equal to 1:

$$\phi_{\text{for}} + \phi_{\text{tsl}} + \phi_{\text{bm}} + \phi_E + \phi_{\text{hk}} = 1 \; . \tag{N6.17}$$

Substituting Eq. (N6.13)–(N6.16) into the sum rule above, and rearranging terms, we obtain the following relation linking the growth rate $\mu$ to the parameters $\nu_C$ and $\nu_R$:

$$(1+\kappa_C)\frac{\mu}{\nu_C} + (1+\kappa_R)\frac{\mu}{\nu_R} + \frac{\mu}{\nu_B} = 1 - \phi_{\text{for},0} - \phi_{\text{tsl},0} - \phi_{\text{bm},0} - \phi_{\text{hk}} \equiv \Delta\phi_{\max} \; , \tag{N6.18}$$

where we defined $\Delta\phi_{\mathrm{max}}$ as the sum of all constant protein mass fractions. Solving for the growth rate:

$$\mu(\nu_C, \nu_R) = \frac{\Delta\phi_{\mathrm{max}}}{(1+\kappa_C)/\nu_C + (1+\kappa_R)/\nu_R + 1/\nu_B} \ . \tag{N6.19}$$

This relation can then be substituted into Eqs. (N6.13)–(N6.16) to yield the protein shares of the other sectors in terms of $\nu_C$ and $\nu_R$. The fit is performed as described in the Methods, and the best-fit values of all parameters are reported in Supp. Table S4.

**6.3.1 Interpretation of the energy sector regulation constraint.** The expression relating the proteome share of the energy sector to the parameters $\nu_C$ and $\nu_R$, Eq. (N6.13), is a simple phenomenological expression that minimally captures the observed dependence of the energy sector on the growth rate. Such functional form can be considered an approximation of a more complete relation, that depends on the protein efficiency of the energy sector $\varepsilon$ (ATP produced per unit of inversted proteome) and the energetic demand of the cell $J_E$ across conditions:

$$\phi_E = \frac{J_E}{\varepsilon} \ . \tag{N6.20}$$

The energy flux is typically modeled as a linear function of the growth rate, $J_E = J_{E,0} + s_E \mu$. As seen in Main Text Fig. 3AB, the cellular energy requirements are significantly different in carbon-limited and translationally-inhibited growth: for a given growth rate, both the inferred ATP maintenance flux and the flux from the energetic pathways is larger in R-limited growth. Such increase can be accounted for by letting $s_E$ to depend on the translational capacity of the cell, $\nu_R$. A possible form for $J_E$ compatible with the increase is the following:

$$J_E = J_{E,0} + \left( s_{bm} + \frac{\kappa_R}{\nu_R} \right) \mu \ . \tag{N6.21}$$

In carbon-limited growth, $\nu_R$ is fixed, and therefore the relation prescribes a linear function of $\mu$ with intercept $J_{E,0}$ and slope $s_{bm} + c_R/\nu_R$. However, in R-limitation, growth rate is changed by changing the translational quality $\nu_R$. This modifies the linear relation between $J_E$ and $\mu$, leading to larger y-axis intercept and a smaller slope.

The protein efficiency of the energy sector depends on several factors, some of which are hard to quantify. Not only it depends on the energetic pathway (respiration vs fermentation) used by the cell, but also by the efficiency of the enzymes in the central carbon pathways, the ATP synthase enzyme and the electron transport chain. In particular, substrate levels in slow growth conditions are likely to be reduced [17]. By comparing the share of energy-associated proteins to the energy flux, we found that the efficiency $\varepsilon$ drops significantly in carbon-limited conditions, while it remains approximately constant in R-limitation (Supp. Fig. S6A). This suggests a positive correlation between $\varepsilon$ and the carbon quality $\nu_C$. On the other hand, the maximal efficiency of the energetic sector is that of the fermentation pathway, $\varepsilon_f$. A minimal form consistent with these observations is a Michaelis-Menten function of $\nu_C$:

$$\varepsilon = \varepsilon_f \frac{\nu_C}{\nu_C + \kappa_C} \ . \tag{N6.22}$$

Substituting Eq. (N6.21) and (N6.22) into Eq. (N6.20) one obtains the explicit dependence of $\phi_E$ on the growth rate, $\nu_C$ and $\nu_R$:

$$\phi_E = \frac{1 + \kappa_C/\nu_C}{\varepsilon_f} \cdot \left[ J_{E,0} + \left( s_{bm} + \frac{\kappa_R}{\nu_R} \right) \mu \right] \ . \tag{N6.23}$$

This expression includes all terms appearing in Eq. (N6.13), as well as other terms. In particular, by expanding Eq. (N6.23), we obtain a constant term; a $k_C\mu/\nu_C$ term, originating from the Michaelis-Menten dependence of the protein efficiency, and a $k_R\mu/\nu_R$, descending from the increased energetic demand in R-limitation. Given the lack of a mechanistic support for Eq. (N6.22), we leave the expression (N6.23) for another study and we only present the results obtained with the simplified expression Eq. (N6.13).

**6.3.2 Reconciliation with the model from Basan *et al.*** The resulting model provide a description of $\phi_E$ in terms of the growth rate and of the quality of the carbon source $\nu_C$, Eq. (N6.13). Furthermore, our modelling allowed to infer the total ATP flux of the energetic pathways, $J_E$, from the difference between ATP maintenance fluxes and the ATP produced in conjunction with the biomass components, Supp. Fig. S4A. Comparing to the protein allocation model from Basan et al. [18], we see that these quantities are the r.h.s. of Eqs. (N6.9)–(N6.10). These relations can be recast as:

$$J_{E,r} + J_{E,f} = J_E \tag{N6.24}$$

$$J_{E,r}/\varepsilon_r + J_{E,f}/\varepsilon_f = \phi_E \ , \tag{N6.25}$$

or, in terms of protein mass fractions:

$$\phi_{E,r} \cdot \varepsilon_r + \phi_{E,f} \cdot \varepsilon_f = J_E \ , \tag{N6.26}$$

$$\phi_{E,r} + \phi_{E,f} = \phi_E \ . \tag{N6.27}$$

These equations fully determine the activity of the respiration and fermentation pathways as a function of $\phi_E$ and $J_E$ (as long as the efficiencies $\varepsilon_r$ and $\varepsilon_f$ are constant). In turn, $\phi_E$ and $J_E$ can be expressed in terms of the model parameters: $\phi_E$ via Eq. (N6.13), and $J_E$ using a similar functional form[3]. Solving the system above for $\phi_{E,f}$ yields the model prediction for the protein mass fraction associated to aerobic fermentation (shown in Fig. 6E):

$$\phi_{E,f} = \frac{J_E - \phi_E \varepsilon_r}{\varepsilon_f - \varepsilon_r} \ . \tag{N6.28}$$

The results in Main Text Fig. 6E are obtained using the expression above, substituting for $\phi_E$ the protein mass fractions estimated via the functional decomposition and for $J_E$ the energy fluxes shown in Main Text Fig. 3. In carbon limitation, the energy flux $J_E$ is positively correlated with the growth rate, while $\phi_E$ is negatively correlated, thus leading to a positive correlation between $\phi_{E,f}$ and the growth rate. Instead, in R-limitation, $\phi_E$ is approximately growth-independent, while $J_E$ has a milder correlation with the growth rate. Therefore, the flux through the fermentation pathway has a much weaker dependence on the growth rate, leading to acetate excretion at slow growth.

[3]The energy flux $J_E$ can be approximately characterized by the value in reference condition, $J_E^{ref}$, and by the slopes in C- and R-limitation, $s_E^C$ and $s_E^R$. These three parameters can be mapped to three parameters in a "growth law" constraint, such as $J_E = c_1 + c_2\mu + c_3\mu/\nu_R$, where $\mu$ is expressed as a function of $\nu_C$ and $\nu_R$ via Eq. (N6.19).

### Supplementary Figures

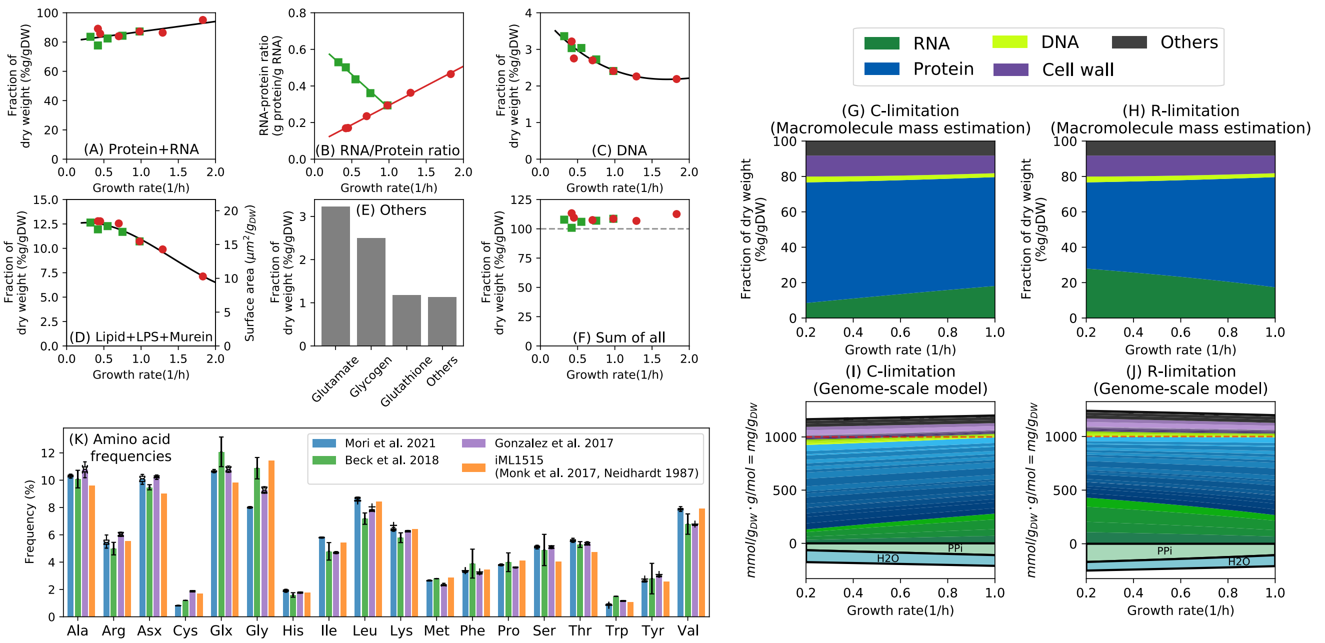

**Figure S1. Biomass composition of *E. coli* cells in carbon-limited conditions or subject to translational inhibition. (A-F)** Experimental constraints used to define a growth-dependent biomass composition for *E. coli* K-12 strain NCM3722 in carbon-limited conditions (red circles) and translationally inhibited growth using sublethal doses of chloramphenicol (green squares), with solid lines indicating a fitted analytical relation used to constrain the metabolic model (see also Supplementary Note 1): **(A)** Sum of protein and RNA mass per dry weight (data from Ref. [5]); **(B)** ratio of protein and RNA mass [5]; **(C)** DNA mass per dry weight [5]; **(D)** Due to the paucity of experimental data, we assumed that the abundance of cell wall/membrane components varied proportionally to the surface area of the cell. The surface area was computed based on the data from Ref. [5], corrected so that the cellular density is consistent with what reported in Ref. [10]. The conversion from surface area to cell membrane mass, as well as the abundance of individual cell membrane components were determined based on Ref. [6–8], see Supplementary Note 1.5.2–3. **(E)** The abundance of other abundant cellular components (glutamate, glycogen, glutathione) was fixed based on available data [8, 12], while the concentration of the other components (coenzyme and ions) were set proportionally to the values reported in the iML1515 metabolic model. **(F)** The sum of all components is slightly above 100%, which likely results from overestimating some of the biomass components. In order to define a condition-dependent biomass composition, we scaled the fitted relationships (solid lines in the previous panels, plus the values for the small molecules) so that their sum is 100%. **(G)** Modeled macromolecular biomass composition in carbon-limited growth. **(H)** Modeled macromolecular composition in translational limitation. **(I)** Summary of the biomass reaction stoichiometry in carbon-limited growth (C-limitation). The biomass reaction includes sink fluxes individual building blocks, such as amino acids (shades of blue) or nucleotides (shades of green), as well as byproduct of their polymerization: water from the formation of peptide bonds, and pyrophosphate from the synthesis of RNA. **(J)** Same as the previous panel, but for translational-limited growth (R-limitation). **(K)** Amino acid frequencies obtained from available proteomics studies [3, 6, 7] or reported in the iML1515 model [1]. Error bars reflect the variability across conditions explored in each study. Since the proteome composition is mostly constant across conditions, we used average abundances obtained from Ref. [3]. This lead in noticeable changes in the demand fluxes for cysteine and glycine compared to the values used in iML1515.

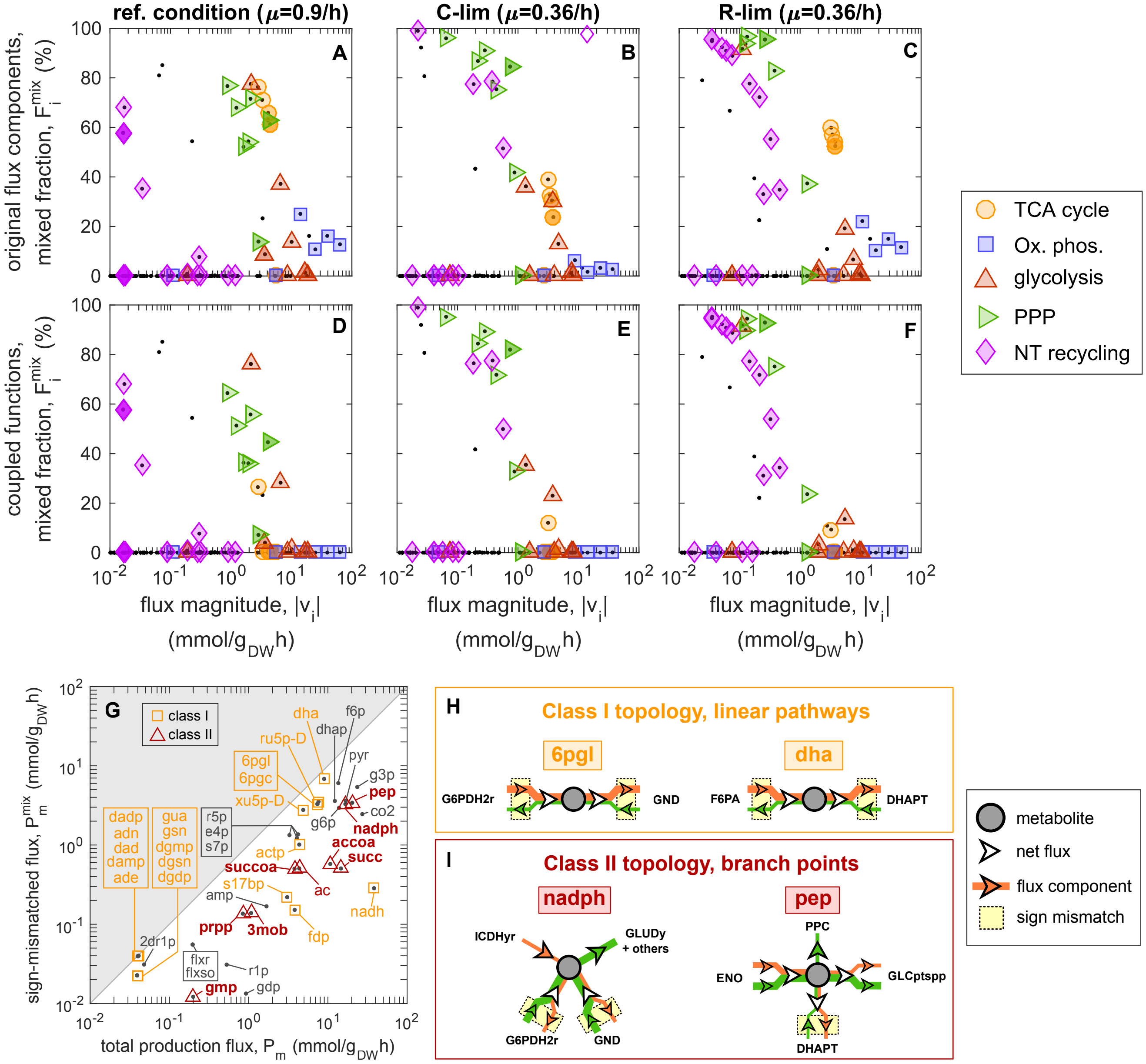

**Figure S2. Global analysis of mixed functional fractions and their relation to network topology.** *(See caption in the next page.)*

**Global analysis of mixed functional fractions and their relation to network topology.** (*See figure in the previous page.*) **(A-C)** For each reaction $i$ with flux components $v_i^{(\gamma)}$ and total flux $v_i$, we calculated the "mixed" functional fractions, $F_i^{\mathrm{mix}} = 1 - |v_i| / \sum_\gamma |v_i^{(\gamma)}|$, which quantify the degree of sign-mismatch among the flux components. The resulting values are here shown here against the flux magnitude, $|v_i$, for aerobic growth in reference condition and in slow carbon-limited or translation-inhibited growth. Large mixed fractions are observed for several reactions, including those associated to TCA cycle (orange circles), oxidative phosphorylation (blue squares), some glycolytic protein (red triangles), pentose phosphate pathway (PPP, green triangles) and several reactions involved in nucleotide metabolism and recycling (purple diamonds). **(D-F)** The mixed fractions were recalculated after linearly combining energetic, acetate-associated and biosynthetic flux components, as described in Fig. 1BC and Supplementary Note 2.3. This approach removed most sign-mismatches for the energetic pathways (TCA and oxidative phosphorylation), but did not remove those associated to PPP and NT recycling. **(G)** To further investigate the emergence of mixed fractions, we computed for each metabolite $m$ its total production flux $P_m$ (by summing over all reaction fluxes producing the metabolite), as well as the production flux $P_m^{\mathrm{mix}}$ arising from sign-mismatched flux components (see Supplementary Note 3 for details), in reference condition. Many metabolites involved in the PPP and in NT metabolism are strongly associated to sign-mismatched flux components, with $P_m^{\mathrm{mix}} \sim P_m$. Metabolites were further classified based on the topology of the associated reactions. Most metabolites belong to either one of two classes: Class I (yellow squares) includes metabolites involved in linear pathways, while class II (red triangles) includes metabolites at key branch points. **(H)** Examples of class I metabolites. The metabolites 6-phospho-D-glucono-1,5-lactone (6pgl) and dihydroxyacetone (dha) are intermediate products of PPP and glycolytic reactions, and constrain the flux components of the producing and consuming reactions to be the same. **(I)** Examples of class II metabolites. These key metabolites lead to the emergence of sign-mismatched flux components, requiring the presence of anaplerotic reactions keeping their concentrations stable under changes in the demand fluxes. Consider for example the electron carrier NADPH. It is produced by the TCA cycle reaction ICDHyr (isocitrate dehydrogenase), and consumed by a variety of biosynthetic reactions, in particular GLUDy (glutamate dehydrogenase). The PPP reactions G6PDH2r (glucose-6-phosphate dehydrogenase) and GND (phosphogluconate dehydrogenase) have to balance not only the overall fluxes, but also the energetic and biosynthetic flux components separately, leading to sign-mismatch. Similarly, the glycolytic reaction DHAPT (dihidroxyacetone phosphotransferase) stabilizes the pool of phosphoenolpyruvate, which is produced and consumed in varying amounts by glycolytic reactions and the glucose PTS transporter.

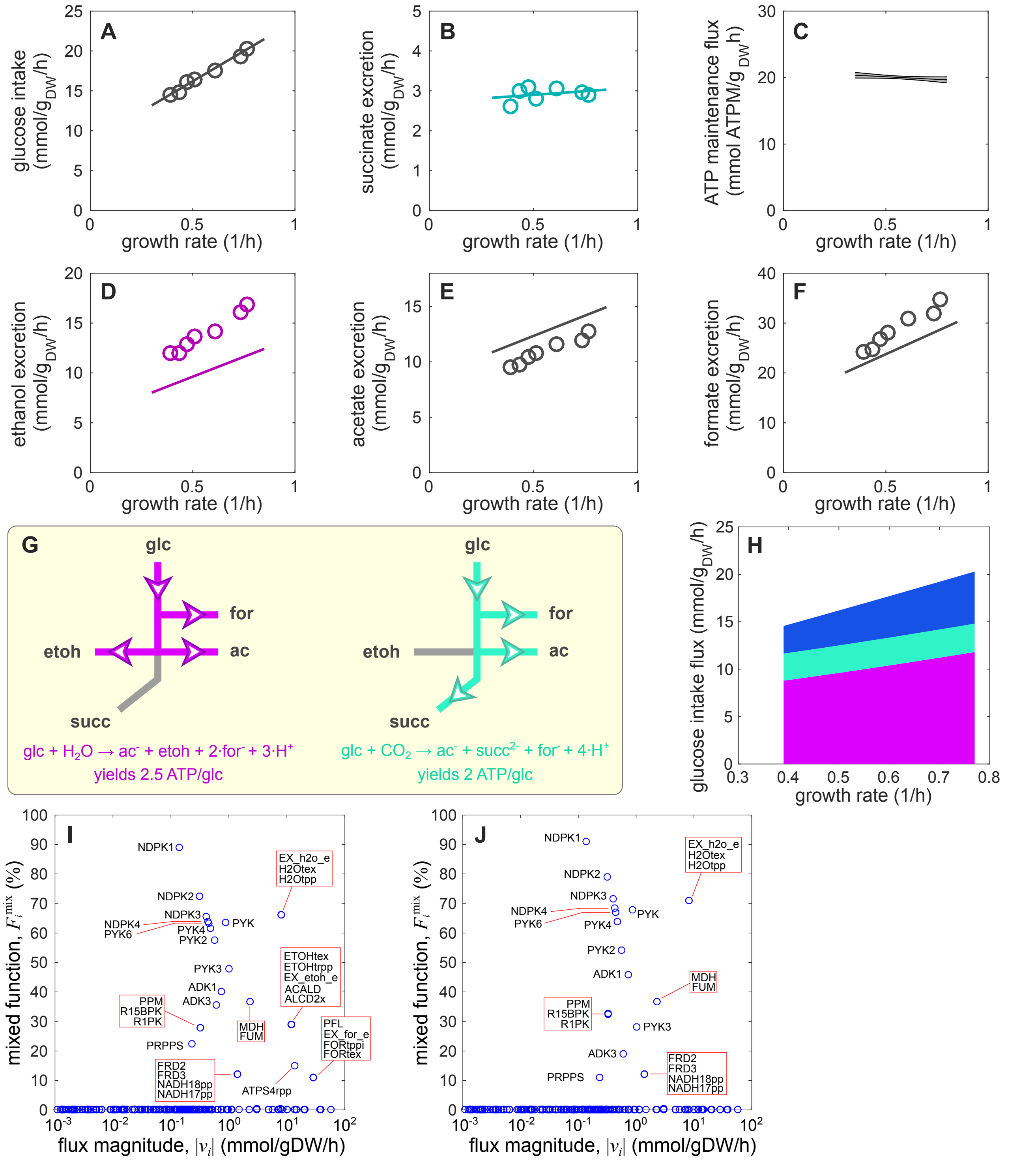

**Figure S3. Metabolic fluxes for cells grown in carbon-limited, anaerobic conditions.** *(See caption in the next page.)*

**Metabolic fluxes for cells grown in carbon-limited, anaerobic conditions.** (*See figure in the previous page.*) **(A-C)** FBA fluxes for anaerobically, glucose-limited *E. coli* cells were estimated by minimizing the glucose intake subject to constraints on succinate excretion and ATP maintenance fluxes (C). Both the succinate excretion and the ATP maintenance flux were modeled as linear functions of the growth rate. The former is estimated from the data (panel B), while the latter was fitted by maximizing the agreement between modeled (panel A, line) and experimental glucose intake rates. **(D-F)** Main fermentation products (symbols), with solid lines indicate the model prediction, without further adjustment compared to the steps outlined in panels (A-C). **(G)** The two flux components corresponding to the ATP maintenance and succinate production constraints were linearly combined to yield two functional components corresponding to distinct mixed fermentation pathways. The first (and dominant) functional component corresponds to the conversion of glucose into acetate, ethanol and formate with an energetic yield of 2.5 ATP/glucose (purple); the second mode corresponds to the conversion of glucose and $CO_2$ into acetate, ethanol and succinate, with an energetic yield of 2.0 ATP/glucose (cyan). **(H)** Prevalence of mixed functional components using the flux components associated to the flux constraints (similar to Supp. Fig. S2A-C) for wild-type cells grown in anaerobic glucose conditions. **(I)** Mixed functional components after the flux coupling procedure (similar to Supp. Fig. S2D-F). The flux-coupling procedure for the anaerobic case is described in Supp. Note 2. **(J)** Functional decomposition of the glucose intake flux in anaerobic conditions. Most of the glucose intake is used for the production of energy via the two fermentation pathways shown in panel (G). About 25% of the flux is used for the production of biomass precursors (in blue). The flux decomposition data for anaerobic growth con glucose at two different growth rates can be found in Supp. File S5.

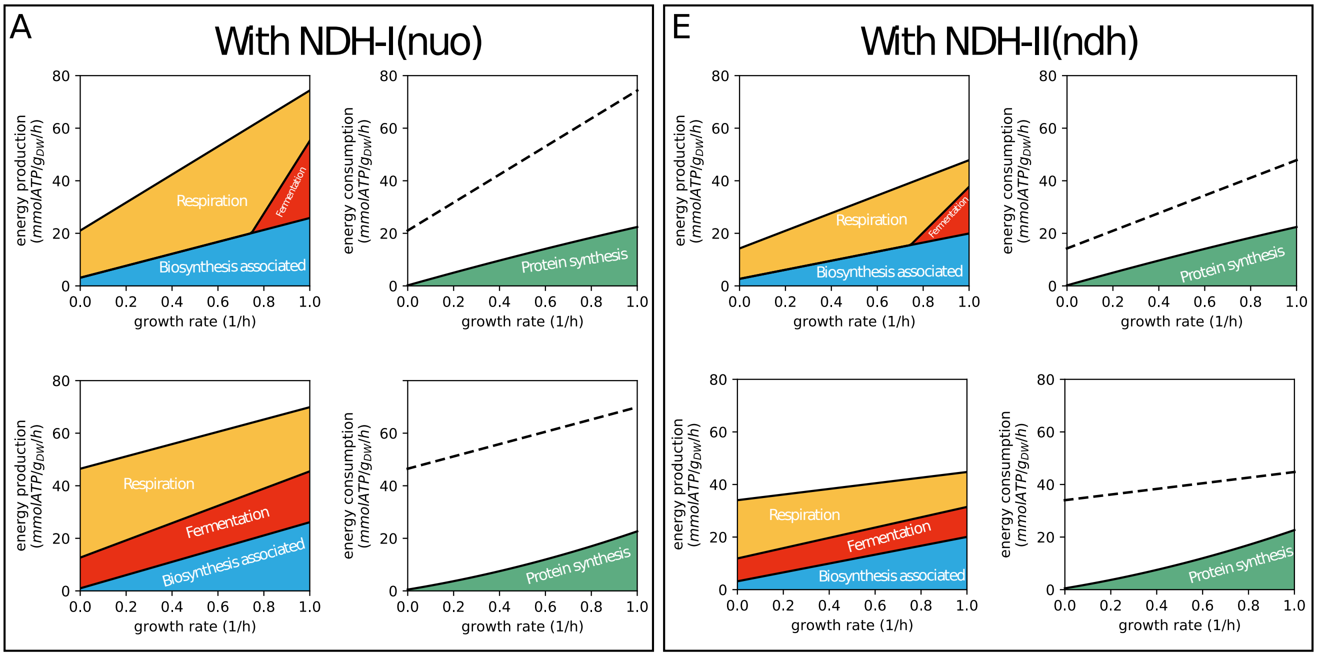

B

| ATP equiv. | NDH-I | NDH-II |
|---|---|---|
| NADH | 2 | 1.25 |
| NADPH | 2.25 | 1.7 |
| FADH2 | 1.75 | 1 |

C

| ATP/glucose | NDH-I | NDH-II |
|---|---|---|
| Respiration | 24 | 17.4 |
| Fermentation | 11 | 8.5 |

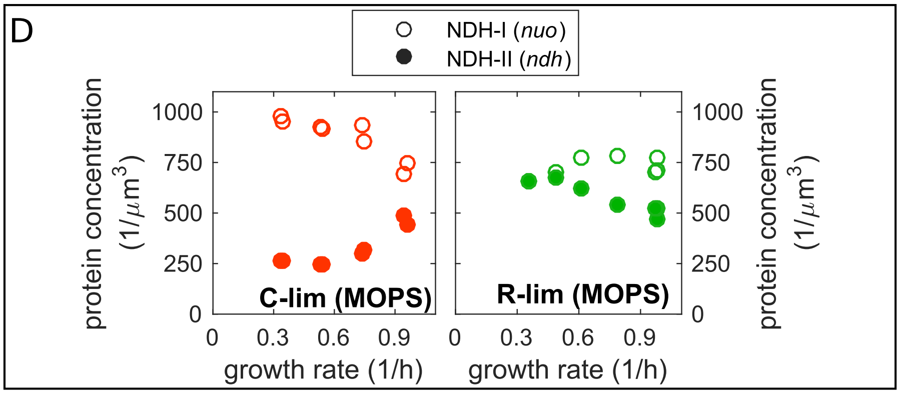

**Figure S4. ATP flux balance and NADH dehydrogenases.** *(See caption in the next page.)*

**ATP flux balance and NADH dehydrogenases.** (*See figure in the previous page.*) **(A)** Summary of the energy balance for C-limited (top) or R-limited (bottom) cells. The plots on the left represent the breakdown of the total ATP synthesis flux (equivalent in this case to the ATP maintenance flux) into ATP producing processes: respiration (yellow), fermentation (red) and the overall ATP production resulting from the synthesis of biomass building blocks on glucose media (blue). The plots on the right compare the same energy flux (black dashed line) to the ATP cost of protein synthesis, assumed to be 4 ATP equivalents per polymerized residue. The difference of the two cannot easily associated to quantifiable energy consuming processes. **(B)** ATP equivalents of various electron carriers when using the more efficient NADH dehydrogenase I (NDH-I, $nuo^+$ $ndh^-$) or the smaller NADH dehydrogenase II (NDH-II, $nuo^-$ $ndh^+$), computed as detailed in Table S3. **(C)** Overall ATP molar yield (ATP produced per glucose molecule) for the respiration and aerobic fermentation pathways using the NDH-I or NDH-II enzymes. **(D)** Concentration of NDH-I and NDH-II enzymes across growth rates in carbon- or translation-limited growth conditions, obtained from experimental mass spectrometry data [3, 20]. The concentration of the NDH-I is estimated as the median concentration of NDH-I subunits. NDH-I is most abundant in slow, carbon-limited growth, and is 3 to 4 times more abundant than NDH-II. However, the concentrations of the two dehydrogenases are comparable in other conditions, in particular in slow, R-limited growth. **(E)** Same as panel (A), but setting to zero the flux through the NDH-I dehydrogenase. Since the ATP yields are lower, the fitted ATP maintenance fluxes are reduced, and so is the difference between the total ATP produced (dashed lines) and the ATP cost of protein synthesis.

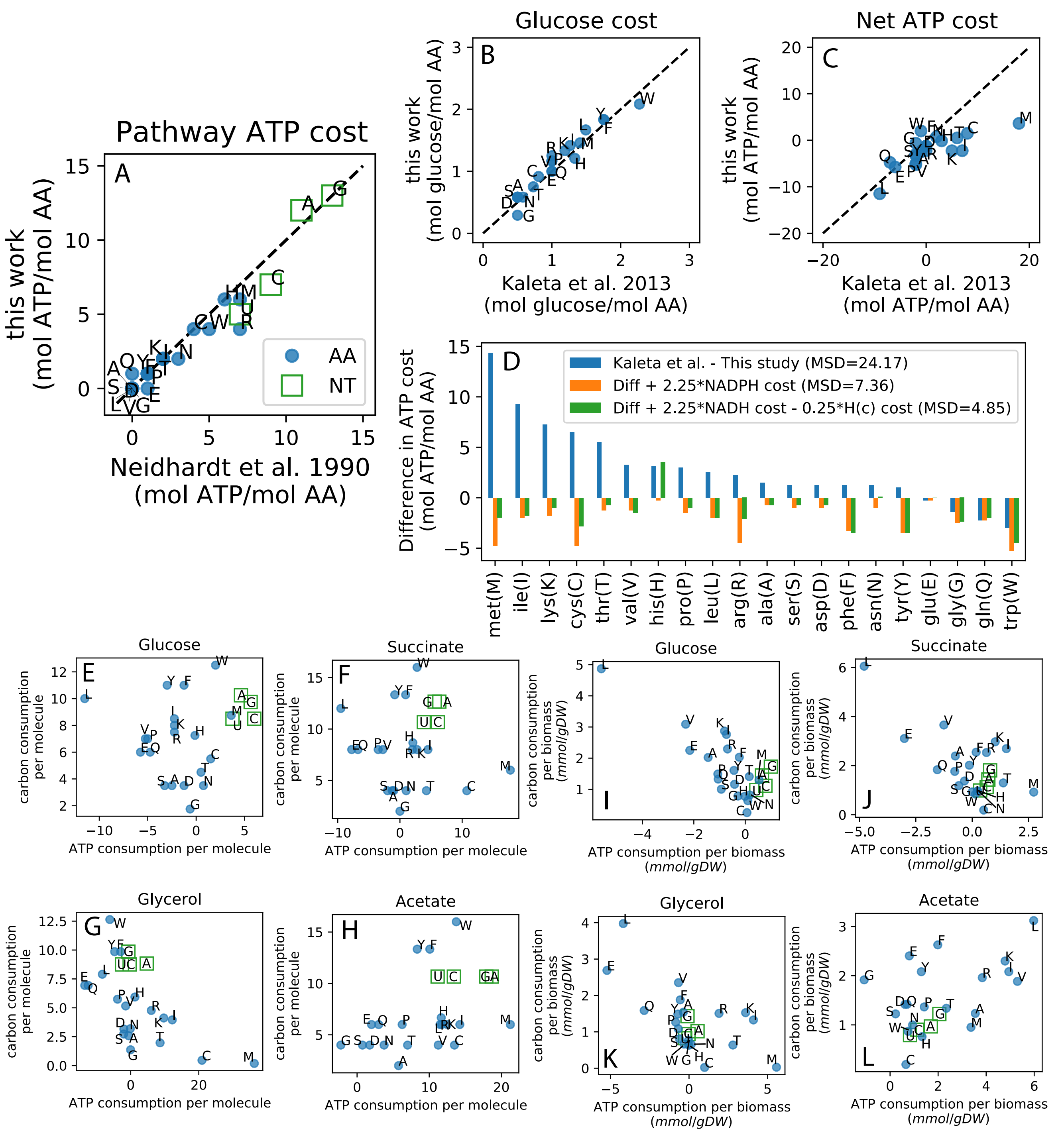

**Figure S5. Carbon and energy costs for the biosynthesis of amino acids and nucleotides.** *(See caption in the next page.)*

**Carbon and energy costs for the biosynthesis of amino acids and nucleotides.** (*See figure in the previous page.*) **(A)** Comparison of anabolic ATP costs for amino acid and nucleotides with those reported in Ref. [8] for glucose minimal media. The costs were computed by summing the contribution of biosynthetic reactions to the ATP balance, excluding reactions in the central carbon pathways (PFK, PFK_3, PGK, PYK, ACKr), as well as ATP synthase (ATPS4rpp) and the ATP maintenance reactions. **(B)** Comparison between carbon costs per amino acid computed in this work with those computed by Kaleta et al. [27] using the "manual" method. The costs computed with other methods, namely "LP standard" and "LP unlimited energy" compare similarly (not shown). **(C)** Comparison between ATP costs per AA computed in this work with those from Ref. [27]. Large differences are seen for several amino acids, with costs from Ref. [27] much larger than those computed with FDM. **(D)** To investigate the difference between the ATP costs observed in panel (C), we first computed the ATP equivalents for various electron carriers in our network (Table S3), obtaining similar values to those assumed in Kaleta et al. [27]. We then inspected the mass balance of the currency metabolites in each metabolic pathway (Supp. File S6). Whereas Kaleta *et al.* directly associated the consumption of one NADPH molecule to the consumption of 2 ATP molecules, our flux solutions balanced the NADPH demand by additional flux through the PPP. Indeed, the difference among our ATP costs is greatly reduced by counting the NADPH produced in the PPP in our flux solutions, and increasing the ATP/AA costs by their ATP equivalent (compare the blue and orange bars). The agreement is further improved when *removing* the ATP contribution due to the production of cytoplasmic protons, which is instead neglected in Kaleta et al. [27] owing to the simplified nature of the network. **(E-H)** Molar carbon (C-atoms) and ATP costs for the synthesis of amino acids and nucleotides for cells grown in glucose (E), succinate (F), glycerol (G) and acetate (H). **(I-L)** Carbon (C-atoms) and ATP requirements per gram of dry biomass produced, for individual building blocks. Panels correspond to cells grown in glucose (I), succinate (J), glycerol (K) and acetate (L). The flux decomposition data for various carbon sources can be found in Supp. File S5.

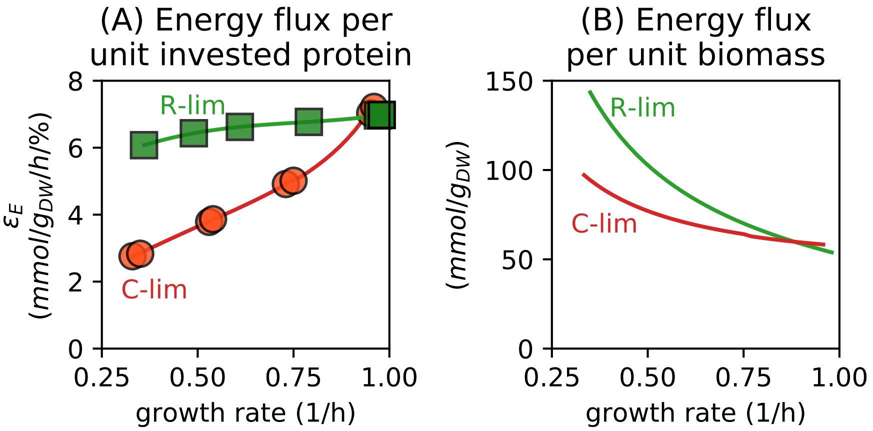

**Figure S6. Efficiency of energy production across conditions.** **(A)** Ratio between the energy flux $J_E$ (same as in Fig. 3AB) and the protein share associated to energy production $\phi_E$ in C- and R-limited growth (red circles and green squares, respectively; solid lines are eye guides). **(B)** ATP demand per unit of biomass, $J_E/\mu$. At slow growth ($\sim$0.3/h), cells grown in R-limited condition (chloramphenicol stress) require much larger ATP per unit of biomass synthesized.

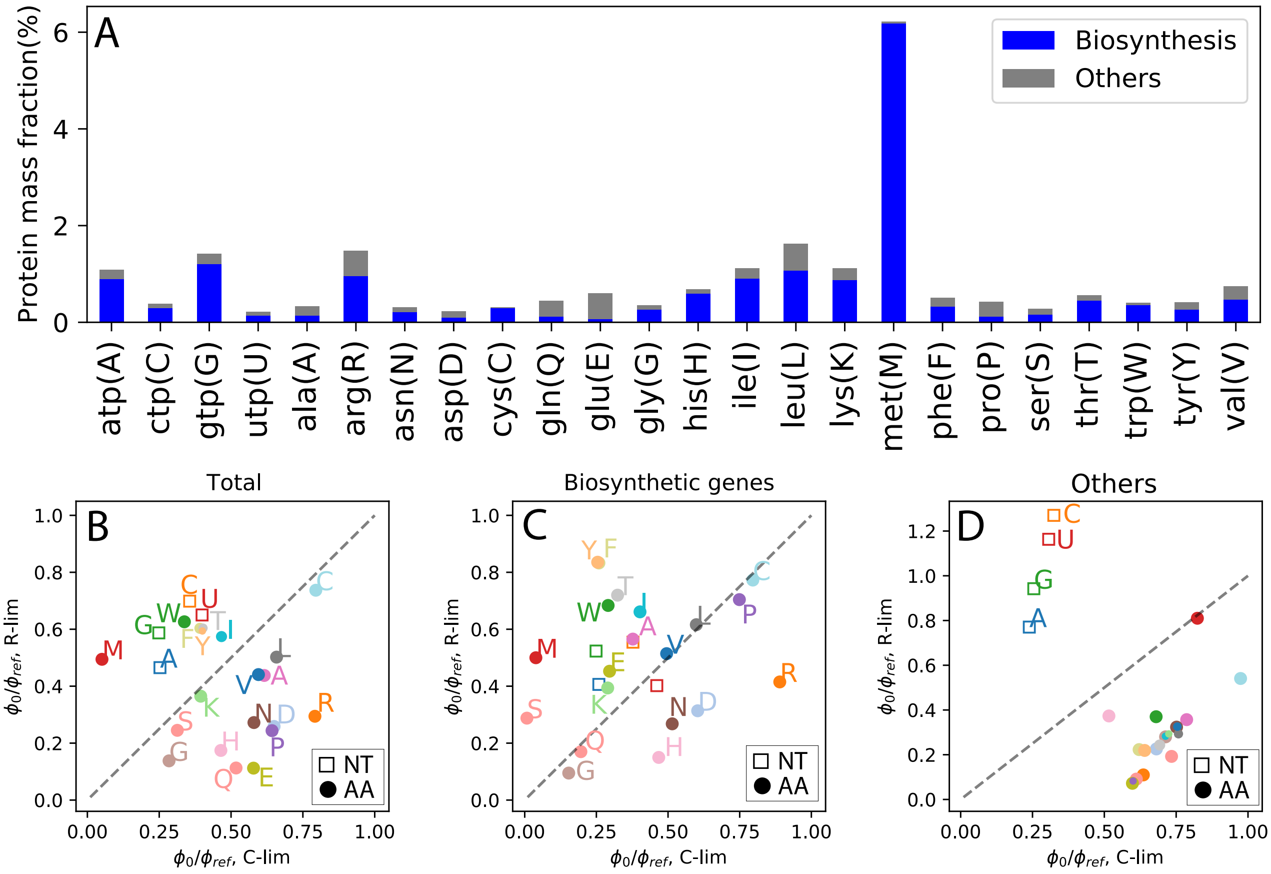

**Figure S7. Fraction of protein costs originating from biosynthetic enzymes. (A)** The protein mass fractions associated to individual amino acids and nucleotides in reference condition (total height of the bars) is broken down into the contribution from biosynthetic enzymes (blue) and the other enzymes (grey), including e.g. central carbon pathways. The proteome allocated to the biosynthesis of most building blocks originates mostly from the biosynthetic enzymes. Instead, the cost for glutamate, glutamine and proline is dominated by the protein cost associated to the non-biosynthetic enzymes. **(B)** For each amino acid and nucleotide, we modeled the relation between the allocated protein mass fraction $\phi$ and the corresponding demand flux as a linear relation. The ratio between its value extrapolated at zero flux, $\phi_0$, and the value in reference condition, $\phi_{\rm ref}$, measures the "flatness" of the protein allocation for different demand fluxes. In this panel, we compared these ratios in C-limited and R-limited growth. For most amino acids, the proteome response was steeper in C-limited condition (below the diagonal, indicated by the dashed line), while for all nucleotides and a few amino acids (methionine, isoleucine, and the aromatic amino acids) the response was steeper in R-limitation. **(C)** When focusing on the contribution from the non-biosynthetic genes (grey bars in panel A), opposite patterns are observed for nucleotides and amino acids. This is consistent with the additional nucleotide production flux required in R-limitation compared to C-limitation (due to the increased RNA/protein ratio, Fig. 2A), which leads to greater allocation towards RNA production of the shared pathways. **(D)** When considering the linear relations for the biosynthetic genes only, the clear patterns observed in panel (B) is much less evident, indicating heterogeneity in the regulation of the biosynthetic proteins for each building block.

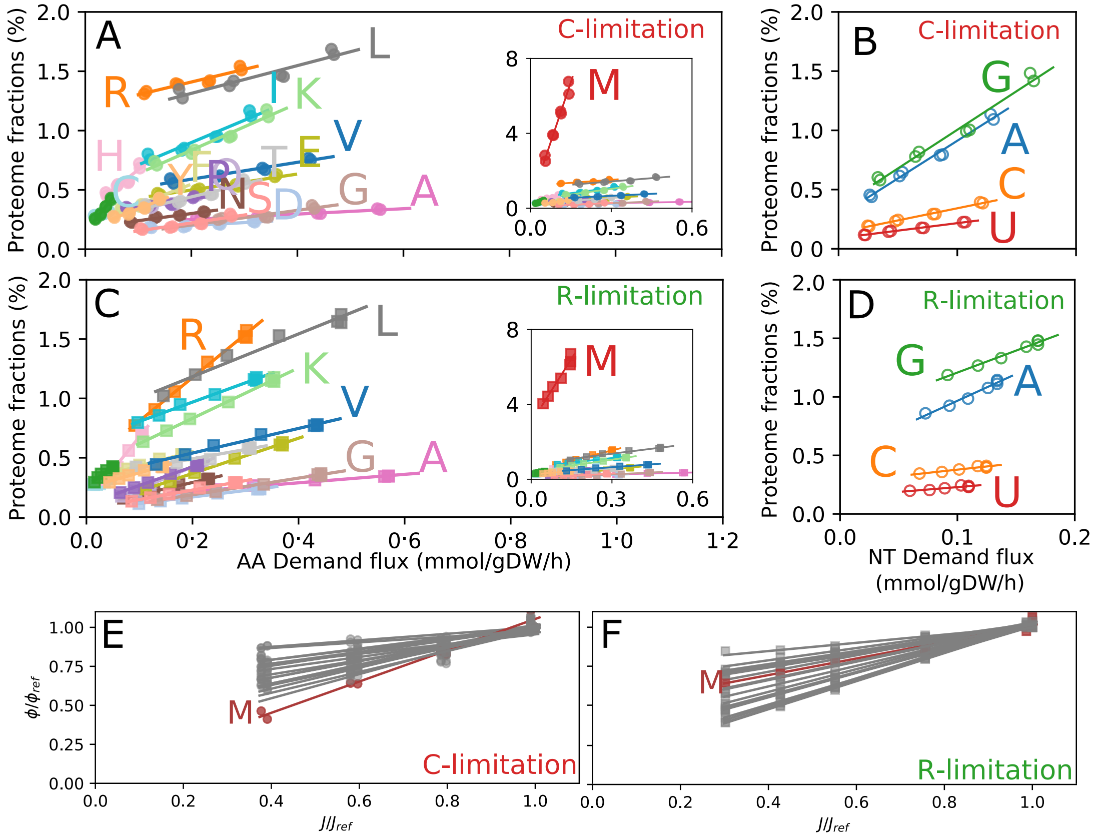

**Figure S8. Protein allocation towards the synthesis of amino acids and nucleotides. (A)** Total protein abundances associated with to the biosynthesis of each amino acid as a function of the amino acid demand flux, in C-limited conditions (same as Main Text Fig. 5I). Solid lines indicate linear fits; their parameters are reported in Supp. File S7. **(B)** Total protein abundances associated with to the *de-novo* biosynthesis of each nucleotide as a function of the corresponding demand flux, in C-limited conditions. **(C)** Same as panel (A), but for R-limited conditions. **(D)** Same as panel (B), but for R-limited conditions. **(E)** Enzyme-flux relationship for AA biosynthesis in carbon-limited conditions, where both the proteins and the demand fluxes associated to each amino acids are normalized to their values in reference condition. (Same data as in panel (A), except for the normalization.) The data and linear relations corresponding to methionine are highlighted in red. **(F)** Same as the previous panel, but for R-limitation data. The change in the protein offset of methionine is clearly visible when comparing with the previous panel.

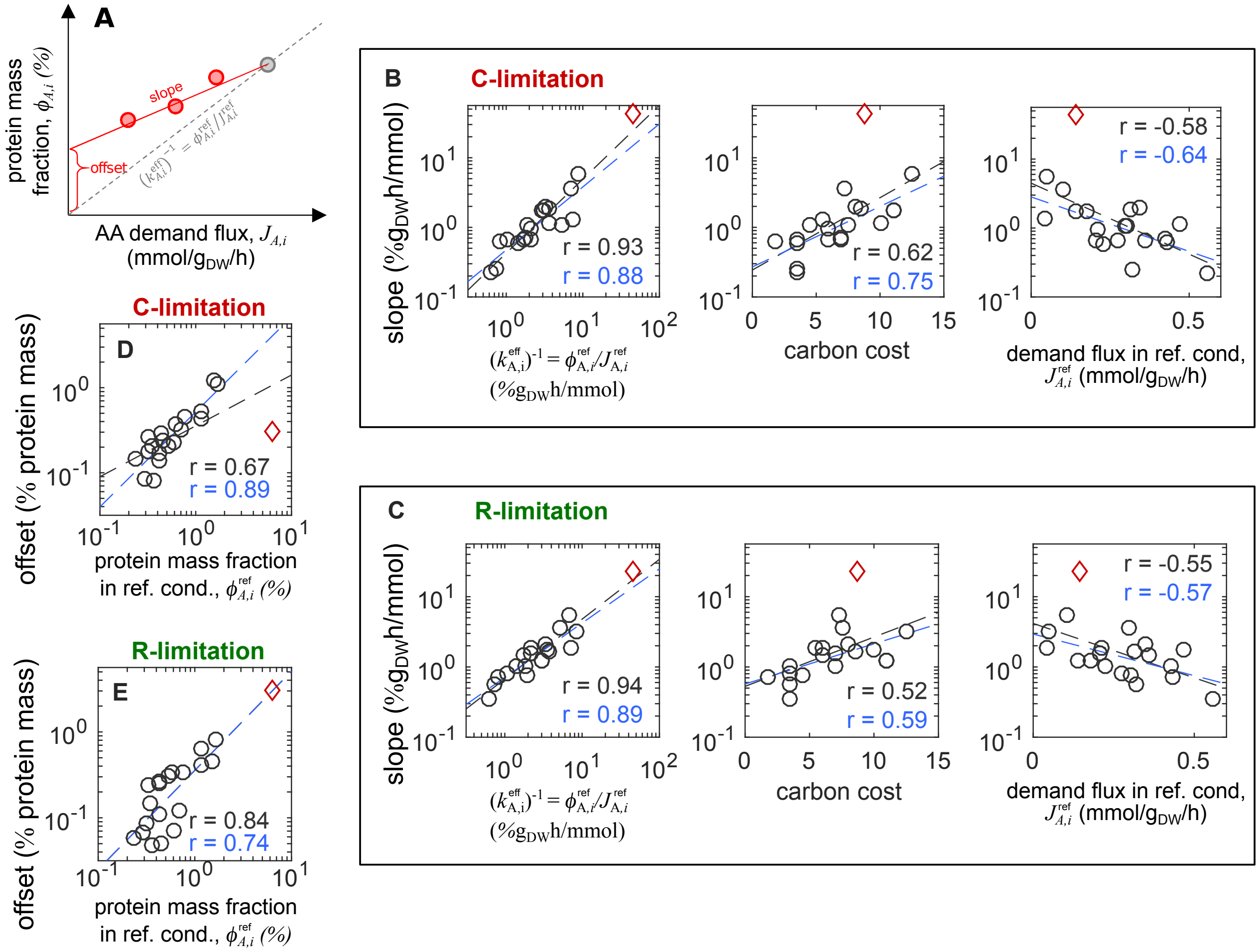

**Figure S9. Comparison between protein allocation and costs across amino acids.** (*See figure in the previous page.*) **(A)** In order to describe the expression of biosynthetic genes across growth rates in a given growth limitation series (see Supp.Fig. S8), we modeled the protein mass fractions $\phi_{A,i}$ associated to amino acid $i$, as a linear function of the demand flux $J_{A,i}$, as indicated by the sketch. In addition, the values of the demand flux and the allocated protein share in reference condition can be used to define the effective turnover number $k^{\text{eff}}_{A,i} = J^{\text{ref}}_{A,i}/\phi^{\text{ref}}_{A,i}$. This is shown in the figure as the inverse of the slope of the dashed grey line. **(B)** The scatter plots compare the slopes of the linear protein-flux relationships obtained in C-limitation to the inverse of the effective turnover number in reference condition, $\left(k^{\text{eff}}_{A,i}\right)^{-1}$, the carbon cost (number of carbon atoms) and the demand flux $J^{\text{ref}}_{A,i}$ in reference condition. The red diamond indicates methionine, which is an evident outlier across all amino acids. Black lines indicate linear fits to the 20 amino acids, while blue lines indicate fits excluding methionine. Pearson correlation coefficients are reported in the same colors. **(C)** Same as the previous panel, but showing the slopes associated to R-limited growth. **(D)** Comparison of the offsets obtained in C-limitation and the protein abundance in reference condition. **(E)** Same as in the previous panel, but for the R-limitation offsets.

**A**

| | C↑ R↓ | C↓ R↑ | C↓ R↓ | C↑ R↑ | Sum |
|---|---|---|---|---|---|
| Energy | 5.66 | 0.93 | 1.64 | 1.15 | 9.38 |
| Biomass | 4.62 | 4.15 | 12.31 | 3.07 | 24.15 |
| Foraging | 5.81 | 0.48 | 0.83 | 1.65 | 8.78 |
| Translation | 0.43 | 23.20 | 1.58 | 0.90 | 26.11 |
| House keeping | 10.07 | 9.12 | 3.82 | 8.56 | 31.57 |
| Sum | 26.59 | 37.88 | 20.19 | 15.34 | 100.00 |

**B**

| | C↑ R↓ | C↓ R↑ | C↓ R↓ | C↑ R↑ | Sum |
|---|---|---|---|---|---|
| Energy | 60.36 | 9.89 | 17.45 | 12.31 | 100 |
| Biomass | 19.12 | 17.19 | 50.98 | 12.71 | 100 |
| Foraging | 66.16 | 5.49 | 9.51 | 18.84 | 100 |
| Translation | 1.66 | 88.83 | 6.06 | 3.46 | 100 |
| House keeping | 31.88 | 28.89 | 12.11 | 27.12 | 100 |

**C**

| | C↑ R↓ | C↓ R↑ | C↓ R↓ | C↑ R↑ |
|---|---|---|---|---|
| Energy | 21.30 | 2.45 | 8.11 | 7.53 |
| Biomass | 17.36 | 10.96 | 60.99 | 20.01 |
| Foraging | 21.85 | 1.27 | 4.14 | 10.78 |
| Translation | 1.63 | 61.24 | 7.84 | 5.88 |
| House keeping | 37.85 | 24.08 | 18.93 | 55.80 |
| Sum | 100.00 | 100.00 | 100.00 | 100.00 |

**D**

| | C | R | A | U | S | C′ | A′ | S′ | X |
|---|---|---|---|---|---|---|---|---|---|
| C-lim | ↑ | ↓ | ↓ | ↓ | ↑ | ↑ | ↓ | ↑ | Not Classified |
| R-lim | ↓ | ↑ | ↓ | ↓ | ↓ | ↑ | ↑ | ↑ | |
| A-lim | ↓ | ↓ | ↑ | ↓ | ↑ | ↓ | ↑ | ↑ | |

**E**

| | C | A | R | U | S | C' | A' | S' | X | Sum |
|---|---|---|---|---|---|---|---|---|---|---|
| Energy | 2.68 | 1.54 | 0.05 | 0.10 | 2.95 | 0.72 | 0.88 | 0.43 | 0.03 | 9.38 |
| Biomass | 1.87 | 5.39 | 1.39 | 6.92 | 2.38 | 0.87 | 2.76 | 2.20 | 0.37 | 24.15 |
| Foraging | 3.11 | 0.48 | 0.41 | 0.35 | 2.22 | 1.29 | 0.07 | 0.37 | 0.48 | 8.78 |
| Translation | 0.00 | 0.41 | 22.98 | 1.18 | 0.31 | 0.07 | 0.21 | 0.83 | 0.12 | 26.11 |
| House keeping | 4.03 | 2.19 | 5.87 | 1.63 | 5.02 | 1.48 | 3.25 | 7.08 | 1.01 | 31.57 |
| Sum | 11.69 | 10.02 | 30.70 | 10.17 | 12.89 | 4.43 | 7.17 | 10.91 | 2.01 | 100.00 |

**F**

| | C | A | R | U | S | C' | A' | S' | X | Sum |
|---|---|---|---|---|---|---|---|---|---|---|
| Energy | 28.53 | 16.43 | 0.50 | 1.02 | 31.48 | 7.67 | 9.38 | 4.63 | 0.35 | 100 |
| Biomass | 7.73 | 22.34 | 5.76 | 28.64 | 9.87 | 3.61 | 11.43 | 9.10 | 1.52 | 100 |
| Foraging | 35.40 | 5.51 | 4.68 | 3.99 | 25.26 | 14.68 | 0.81 | 4.16 | 5.50 | 100 |
| Translation | 0.00 | 1.56 | 88.01 | 4.50 | 1.20 | 0.27 | 0.82 | 3.18 | 0.46 | 100 |
| House keeping | 12.77 | 6.94 | 18.60 | 5.17 | 15.90 | 4.69 | 10.29 | 22.43 | 3.21 | 100 |

**G**

| | C | A | R | U | S | C' | A' | S' | X |
|---|---|---|---|---|---|---|---|---|---|
| Energy | 22.91 | 15.39 | 0.15 | 0.94 | 22.92 | 16.25 | 12.27 | 3.98 | 1.65 |
| Biomass | 15.98 | 53.85 | 4.53 | 68.02 | 18.49 | 19.69 | 38.47 | 20.14 | 18.22 |
| Foraging | 26.60 | 4.83 | 1.34 | 3.45 | 17.21 | 29.08 | 0.99 | 3.35 | 23.97 |
| Translation | 0.00 | 4.07 | 74.85 | 11.55 | 2.43 | 1.59 | 2.97 | 7.62 | 5.94 |
| House keeping | 34.51 | 21.86 | 19.12 | 16.04 | 38.95 | 33.38 | 45.29 | 64.90 | 50.22 |
| Sum | 100 | 100 | 100 | 100 | 100 | 100 | 100 | 100 | 100 |

**Figure S10. Comparison between function-based and regulation-based protein sectors. (A-C)** Intersection of the five functional sectors defined in Fig. 6 in terms of regulation-based groups, in which proteins are distinguished based on they up- or down-regulation in either C- or R-limitation, as indicated by the arrows. Panel (A) is the same as Fig. 6C. Panels (B) shows the breakdown of the functional sectors as percentages of the regulation-based sectors; panel (C) indicates the breakdown of regulation-based sectors in terms of the functional sectors. **(D)** The coarse-grained scheme defined in Mori et al. [3] (a slight extension of the sectors introduced in Hui et al. [25]) split each of the four regulation sectors defined above according to the protein's response in anabolic (A-) limitation, corresponding to the titration of nitrogen uptake, for a total of eight protein sectors. The three C', A' and S' sectors mostly include mildly varying proteins that were associated to the same O-sector in Hui et al. [25]. Additionally, an X-sector includes proteins which could not be consistently classified because of missing data. **(E-G)** Same as in panels A through C, except that in this case we describe the intersection of functional sectors with the regulation-based sectors defined in panel (D).

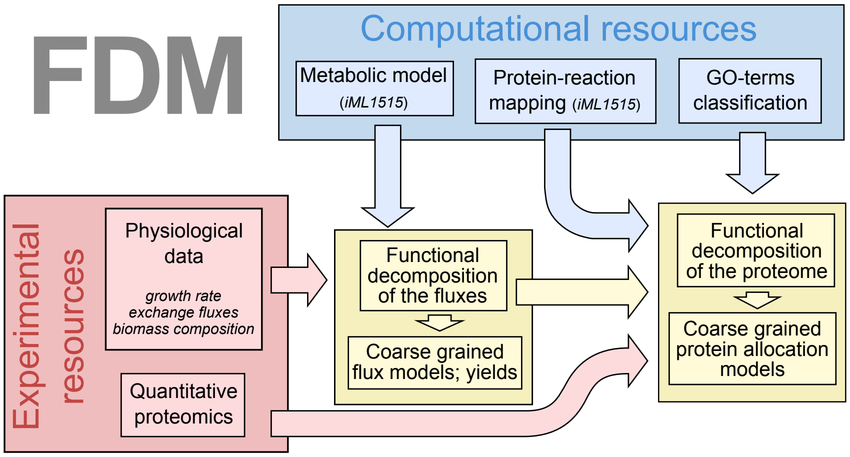

**Figure S11. Workflow of the study of cellular metabolism and gene expression using the Functional Decomposition of Metabolism.** This scheme summarizes how the Functional Decomposition of Metabolism (FDM) combines information from experimental and computational resources. The combination of physiological data (growth rates, exchange fluxes, biomass composition) and *in-silico* models of metabolism allows to estimate intracellular fluxes, define the functional decomposition of the metabolic network, and to calculate yields and costs for the biosynthesis of individual biomass components. Experimental protein abundances can be studied via FDM using the protein-reaction associations provided by the *in-silico* models, as well as available GO-term classifications for non-metabolic proteins.

### Supplementary Tables

| Strains used | Genotype | Description | Source |
|---|---|---|---|
| NCM3722 | *E. coli* K-12 | Wild type | Ref. [28–30] |
| NQ1243 | *ycaD*::FRT:$P_{tet}$:*xylR* $P_{ptsG}$::*kan*:Pu:*ptsG* | Titratable *ptsG* | Ref. [18] |
| NQ1390 | *ycaD*::FRT:$P_{lacIq}$:*xylR* $P_{ptsG}$::*kan*:Pu:*ptsG* | Titratable *ptsG* | Ref. [17] |
| NQ1261 | *ptsG*::FRT | *ptsG* deletion | Ref. [21] |
| NQ1448 | *ycaD*::FRT:$P_{lacIq}$:*xylR* $P_{ptsG}$::Pu:*ptsG* | Titratable *ptsG* | Ref. [31] |
| NQ1554 | *ycaD*::FRT:$P_{tet}$:*xylR* FRT:Pu:*ptsG* ΔmanX::*kan* | Titratable *ptsG* | This work (from NQ1243) |

**Table S1. Strains used in this work.** To construct NQ1554, NQ1243 was transformed with the Pcp20 plasmid [32] to flip out the kanamycin marker. The resulting strain was then used as a recipient strain for P1 transduction with P1 lysate prepared from the Keio collection [33] to create the $\Delta manX$::*kan* mutant, NQ1554.

| Condition | $\sigma$(mmolATP/$g_{DW}$) | $\sigma_0$(mmolATP/$g_{DW}h$) |
|---|---|---|
| C-limited | $53 \pm 13$ | $21 \pm 10$ |
| R-limited | $23 \pm 12$ | $46 \pm 8$ |
| C-limited, anaerobic | $20.9 \pm 1.0$ | $-1.5 \pm 1.7$ |

**Table S2. Fitted ATP maintenance parameters.** The ATP maintenance (ATPM) flux is modeled as a linear function of the growth rate $\mu$ as $J_{ATPM} = \sigma_0 + \sigma\mu$. The two parameters $\sigma$ and $\sigma_0$, termed growth-dependent ATP maintenance (GAM) and non growth-dependent ATP maintenance (NGAM), respectively, are obtained by fitting experimentally determined glucose uptake flux (see Methods). The results are reported in Main Text Fig. 1D and Supp. Fig. S3C.

| Metabolite | dissipation reaction | ATP equivalent cost |
|---|---|---|
| NADH | $NADH + \frac{1}{2}O_2 + H^+ \longrightarrow NAD^+ + \frac{1}{2}H_2O$ | 2 |
| NADPH | $NADPH + \frac{1}{2}O_2 + H^+ \longrightarrow NADP^+ + \frac{1}{2}H_2O$ | 2.25 |
| $FADH_2$ | $FADH_2 + \frac{1}{2}O_2 \longrightarrow FAD + H_2O$ | 1.75 |
| $H^+$(c) | $H^+(c) \longrightarrow H^+(p)$ | 0.25 |

**Table S3. Conversion factors between electron carriers and ATP.** Costs for the electron carriers NADH, NADPH and $FADH_2$ were computed by adding "dissipation" reactions describing the oxidation of the electron carriers, thus "wasting" the electrons. The ATP equivalent cost was computed as the increase in ATP demand (change in ATPM reaction) associated to a small flux through each dissipation reaction. Similarly, the leakage of protons from the cytoplasm (c) to the periplasm (p) is balanced by an increase in ATPM flux. (See Supplementary Note 4 for details.)

| Parameter | Best fit |
|---|---|
| $\phi_{\mathrm{for},0}$ | $0.023 \pm 0.010$ |
| $\phi_{\mathrm{tsl},0}$ | $0.137 \pm 0.008$ |
| $\phi_{\mathrm{bm},0}$ | $0.185 \pm 0.007$ |
| $\phi_{\mathrm{E},0}$ | $0.019 \pm 0.042$ |
| $\phi_{\mathrm{hk},0}$ | $0.306 \pm 0.002$ |
| $\kappa_c$ | $0.727 \pm 0.228$ |
| $\kappa_r$ | $0.220 \pm 0.199$ |
| $\nu_{\mathrm{bm}}$ | $8.142 \pm 0.597$ |
| $\nu_{\mathrm{C}}$ (ref. cond., $\mu = 0.97/h$) | $24.411 \pm 7.68$ |
| $\nu_{\mathrm{R}}$ (ref. cond., $\mu = 0.97/h$) | $8.220 \pm 0.7$ |
| $\nu_{\mathrm{C}}$ (C-lim, $\mu = 0.37/h$) | $2.886 \pm 0.231$ |
| $\nu_{\mathrm{R}}$ (R-lim, $\mu = 0.34/h$) | $1.565 \pm 0.091$ |

**Table S4. Best fit parameter values.** Best fit parameters for the protein allocation model applied to the data in Fig. 6D. The model is explained in detail in Supp. Note 6, while the fitting procedure is explained in the Methods.

#### Supplementary Files

- **Supplementary File S1.** Set of modified iML1515 models with condition-dependent biomass composition, provided as 3 *json* files zipped. The three *json* files correspond to the FBA models of *E. coli* K-12 strain NCM3722 under glucose minimal MOPS media (reference condition, growth rate = 0.96/h), under glucose minimal MOPS media with chloramphenicol treatments (R-limitation, growth rate = 0.35/h), and glucose-limited MOPS growth (C-limitation, growth rate = 0.33/h).

- **Supplementary File S2.** Growth rates and metabolic fluxes used to constrain the models in different conditions, using the conversion factor 1 $OD_{600} \cdot L = 0.5\ g_{DW}$ [5].

- **Supplementary File S3.** The *xlsx* file includes fluxes and the functional decomposition of each reaction into the functional component for the 3 FBA models in Supplementary File S1.

- **Supplementary File S4.** The *xlsx* file includes hierarchical clustering of different reactions based on their functional decomposition. Two sheets are in the file, which include: (1) the reactions and metabolic functions that are included in Main Text Fig. 4A; (2) all the metabolic functions and active reactions.

- **Supplementary File S5.** The *xlsx* file includes fluxes and the functional decomposition of each reaction into the functional component for anaerobic growth on glucose at two different growth rates, as well as aerobic growth on a variety of carbon sources.

- **Supplementary File S6.** The *xlsx* file includes three sheets: (1) the growth rates and ATP yields under different carbon sources; (2) energy pathways of balancing protons or electron carriers (NADH, NADPH, FADH2); (3) biosynthesis reactions for each biomass precursor that involve in energy production/consumption.

- **Supplementary File S7.** The *xlsx* file includes glucose and ATP costs, as well as protein allocation parameters (slopes and intercepts for the protein mass fractions, versus the demand flux for C- and R-limited growth) for individual amino acids and nucleotides.

[8] Frederick Carl Neidhardt, John L Ingraham, and Moselio Schaechter. *Physiology of the bacterial cell: a molecular approach.* Sinauer Associates Sunderland, MA, 1990.

[9] G Bratbak and I Dundas. Bacterial dry matter content and biomass estimations. *Appl. Environ. Microbiol.*, 48(4):755–757, October 1984.

[10] William W Baldwin, Richard Myer, Teresa Kung, Erika Anderson, and Arthur L Koch. Growth and

buoyant density of Escherichia coli at very low osmolarities. *Journal of Bacteriology*, 177(1):235–237, 1995.

[11] Fangwei Si, Dongyang Li, Sarah E Cox, John T Sauls, Omid Azizi, Cindy Sou, Amy B Schwartz, Michael J Erickstad, Yonggun Jun, Xintian Li, and Suckjoon Jun. Invariance of initiation mass and predictability of cell size in escherichia coli. *Curr. Biol.*, 27(9):1278–1287, May 2017.

[12] Junyoung O Park, Sara A Rubin, Yi-Fan Xu, Daniel Amador-Noguez, Jing Fan, Tomer Shlomi, and Joshua D Rabinowitz. Metabolite concentrations, fluxes and free energies imply efficient enzyme usage. *Nature chemical biology*, 12(7):482–489, 2016.

[13] Daniele De Martino, Fabrizio Capuani, Matteo Mori, Andrea De Martino, and Enzo Marinari. Counting and correcting thermodynamically infeasible flux cycles in genome-scale metabolic networks. *Metabolites*, 3(4):946–966, 2013.

[14] Robert Cummins. Functional analysis. *The Journal of Philosophy*, 72(20):741–765, 1975. doi: 10.2307/2024640.

[15] Philippe Huneman. *Functions: selection and mechanisms.* Springer, 2013.

[16] Matteo Mori, Enzo Marinari, and Andrea De Martino. A yield-cost tradeoff governs Escherichia coli's decision between fermentation and respiration in carbon-limited growth. *npj Systems Biology and Applications*, 5(1):16, May 2019. doi: 10.1038/s41540-019-0093-4.

[17] Karl Kochanowski, Hiroyuki Okano, Vadim Patsalo, James Williamson, Uwe Sauer, and Terence Hwa. Global coordination of metabolic pathways in escherichia coli by active and passive regulation. *Molecular systems biology*, 17:e10064, 2021.

[18] Markus Basan, Sheng Hui, Hiroyuki Okano, Zhongge Zhang, Yang Shen, James R Williamson, and Terence Hwa. Overflow metabolism in Escherichia coli results from efficient proteome allocation. *Nature*, 528(7580):99–104, 2015.

[19] Daniel A Beard and Hong Qian. *Chemical biophysics: Quantitative analysis of cellular systems.* Cambridge University Press, 2008.

[20] Chenhao Wu, Matteo Mori, Zhongge Zhang, Amir Banaei-Esfahani, Hiroyuki Okano, Ruedi Aebersold, Christina Ludwig, and Terence Hwa. Proteome reserves define the onset of enzyme recovery across biosynthesis pathways. *Accepted for publication*, 2022.

[21] Xiongfeng Dai, Manlu Zhu, Mya Warren, Rohan Balakrishnan, Vadim Patsalo, Hiroyuki Okano, James R Williamson, Kurt Fredrick, Yi-Ping Wang, and Terence Hwa. Reduction of translating ribosomes enables Escherichia coli to maintain elongation rates during slow growth. *Nature microbiology*, 2(2):16231, 2017.

[22] Conghui You, Hiroyuki Okano, Sheng Hui, Zhongge Zhang, Minsu Kim, Carl W Gunderson, Yi-Ping Wang, Peter Lenz, Dalai Yan, and Terence Hwa. Coordination of bacterial proteome with metabolism by cyclic AMP signalling. *Nature*, 500(7462):301–306, 2013.

[23] Matthew Scott, Carl W Gunderson, Eduard M Mateescu, Zhongge Zhang, and Terence Hwa. Interdependence of cell growth and gene expression: origins and consequences. *Science*, 330(6007): 1099–1102, 2010.

[24] Rohan Balakrishnan, Matteo Mori, Igor Segota, Zhongge Zhang, Ruedi Aebersold, Christina Ludwig, and Terence Hwa. Principles of gene regulation quantitatively connect dna to rna and proteins in bacteria. *bioRxiv*, 2021.
